# Synergistic roles of Aquaporin 5 and Intra- and Extracellular Carbonic Anhydrases in promoting CO_2_ Diffusion across the *Xenopus* Oocyte Plasma Membrane

**DOI:** 10.1101/2025.04.08.647833

**Authors:** Deng-Ke Wang, Fraser J. Moss, Walter F. Boron

## Abstract

**Key Points:**

- According to Fick’s law, transmembrane CO_2_ flux (*J*CO_2_) is the product of membrane permeability (*P*_M,CO2_) and transmembrane concentration gradient (Δ[CO_2_]): *J*CO_2_=*P*_M,CO2_Δ[CO_2_]. Previous work separately showed that (1) human aquaporin-5 (hAQP5) enhances *P*_M,CO2_, and (2) intracellular and (3) extracellular carbonic anhydrases (CAs) enhance Δ[CO_2_] by consuming accumulated or replenishing lost CO_2_. We now examine interactio ns among #1–#3.
- We assess CO_2_ fluxes—produced by addition/removal of extracellular CO_2_/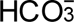—using microelectrodes to monitor extracellular-surface pH (pH_S_) and intracellular pH (pH_i_) of *Xenopus* oocytes heterologously expressing hAQP5, injected with human CA II (hCA II), and/or exposed to extracellular bovine CA (bCA).
- Enhancing effects on CO_2_ fluxes are synergistic among hAQP5, hCA II, and bCA, any of which can become rate limiting, depending on the status of the other two.
- CO_2_/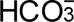 addition transiently increases pH_S_ (ΔpH_S_), hCA II augments ΔpH_S_ (ΔΔpH_S_), and hAQP5 enhances ΔΔpH_S_ (ΔΔΔpH_S_)—a novel tool to assess potential CO_2_ channels.

CO_2_ diffusion across plasma membranes depends on both membrane CO_2_ permeability (*P*_M,CO2_) and transmembrane CO_2_ concentration gradient (Δ[CO_2_])—Fick’s law. Human aquaporin-5 (hAQP5) accelerates CO_2_ diffusion by increasing *P*_M,CO2_, whereas carbonic anhydrases (CAs) accelerate CO_2_ diffusion by enhancing CO_2_ consumption/production and thus Δ[CO_2_]. Here, we systematically assess functional interactions among a gas channel and intra-/extracellular CAs. On Day 1, we inject *Xenopus* oocytes with cRNA encoding hAQP5 (control: H_2_O). On Day 4, we inject hCA II protein in “Tris” buffer (control: “Tris”). We assess CO_2_ fluxes by introducing extracellular 1.5% CO_2_/10 mM 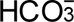 and using microelectrodes to measure (1) maximal extracellular-surface pH increase ΔpH_S_, (2) maximal rate of pH_S_ relaxation (dpH_S_/dt)_Max_, and (3) maximal rate of intracellular-pH decrease (dpH_i_/dt)_Max_. By itself, hCA II minimally increases ΔpH_S_—measured “trans” to added cytosolic CA (CA_i_)—even at highest doses (100 ng/oocyte). However, hAQP5 alone triples ΔpH_S_, an effect further doubled by increasing hCA II. By itself, bovine erythrocyte CA (bCA) in the extracellular fluid doubles (dpH_i_/dt)_Max_ magnitude—measured “trans” to added extracellular CA (CA_o_)—an effect further doubled by hAQP5. Note: pH measurements “cis” to added CAs—pH_S_ for bCA, (dpH_i_/dt)_Max_ for hCA II—are overwhelmed by enzymatical ly-produced/consumed H^+^, and cannot provide intuitive insight into CO_2_ fluxes. Our “trans” pH measurements: (1) confirm synergy between CA_o_ and CA_i_; establish synergy between hAQP5 and both (2) CA_o_ and (3) CA_i_; and show that enhancement of ΔpH_S_ by CA_i_ (ΔΔpH_S_) is a useful tool for assessing CO_2_ permeability of membrane proteins (e.g., hAQP5).

## Introduction

Aquaporins (AQPs) are integral membrane proteins named for their ability to conduct H_2_O across biological membranes. The AQPs occur ubiquitously across all kingdoms of life (King *et al*., 2004; Maurel *et al*., 2015; Ishibashi *et al*., 2017; Ni *et al*., 2017). To date, the cDNAs encoding 13 human AQPs have been cloned and characterized (for review, see Li and Wang, 2017). One can divide them into three groups, based on whether or not they can conduct glycerol in addition to H_2_O. The first group—the classical aquaporins—comprises AQP0, 1, 2, 4, 5, 6, and 8. Although some classical aquaporins are also permeable to urea (AQP6 and AQP8) or ions (AQP6), all of them have a high permeability to H_2_O. The second group—the aquaglyceroporins—includes AQP3, 7, 9 and 10 (Ishibashi *et al*., 1997; Heymann & Engel, 1999; Michalek, 2016; Zhang *et al*., 2019; Moss *et al*., 2020). Besides H_2_O, they can mediate the diffus io n of glycerol, urea, and other non-volatile solutes. Finally, the third group consists of AQP11 and AQP12, which have the lowest homologies to the other aquaporins, and uncertain functions (Ishibashi, 2009; Calvanese *et al*., 2013).

In addition to conducting H_2_O, glycerol, urea, and other small solutes, some AQPs serve as conduits for the highly volatile solutes CO_2_ and NH_3_ (for review, see Boron, 2010), and probably O_2_ (Moss *et al*., 2025; Occhipinti *et al*., 2025; Zhao *et al*., 2025). Nakhoul *et al*., (1998) and Cooper & Boron, (1998) used the rate of intracellular acidification (dpH_i_/dt) to examine the effects of expressing human AQP1 on CO_2_ diffusion across the plasma membrane of *Xenopus laevis* oocytes. They were the first to observe that a membrane protein can act as a channel for a dissolved gas. Later, Endeward *et al*., (2006) confirmed the CO_2_ permeability of AQP1 in human red blood cells, using mass spectrometry to monitor the disappearance of C^18^O^16^O. As part of the Endeward study, Musa-Aziz introduced the technique of using the rapid, transient increase of surface-pH (pH_S_)—during the application of CO_2_/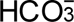—as a semiquantitative index of the CO_2_ influx across the oocyte membrane. This pH_S_ approach is technica lly simpler than the others, and its use led to the elucidation of the roles that several other AQPs—including AQP5—play as CO_2_ channels (Musa-Aziz *et al*., 2009; Geyer *et al*., 2013). Mathematical modeling has provided quantitative insight into the events surrounding CO_2_ influx, including the time course of pH_S_ (Somersalo *et al*., 2012; Musa-Aziz *et al*., 2014*a*, 2014*b*; Occhipinti *et al*., 2014; Calvetti *et al*., 2020).

Raina *et al*., (1995) were the first to clone the cDNA encoding AQP5, obtaining the clone from a library of rat submandibular gland. Others localized AQP5 to the apical membrane of acinar cells in secretory glands, such as salivary glands (Nielsen *et al*., 1997; Ma *et al*., 1999; Steinfeld *et al*., 2001; Gresz *et al*., 2001; Larsen *et al*., 2011; Yoshimura *et al*., 2016) and lacrimal glands (Ishida *et al*., 1997; Tsubota *et al*., 2001). AQP5 also is present in the apical membranes of secretory cells of pyloric glands and duodenal glands (Parvin *et al*., 2002; Matsuzaki *et al*., 2003), as well as sweat glands (Nejsum *et al*., 2002). In the lung, AQP5 in rats and humans is localized to apical membrane of alveolar type I pneumocytes, but not type II pneumocytes (Nielsen *et al*., 1997; Funaki *et al*., 1998; Kreda *et al*., 2001). However, in mice, AQP5 is present in the apical membranes of both type I and type II pneumocytes (Krane *et al*., 2001; Matsuzaki *et al*., 2009). In the kidney, AQP5 co-localizes with pendrin at the apical membrane of β-intercalated cells (β-ICs) that secrete 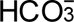 into the lumen of cortical collecting duct (Procino *et al*., 2011). In the aforementioned cells, AQP5 presumably plays an important role in transepithelial fluid and electrolyte transport either because of its role as a H_2_O channel or, perhaps in the case of β-intercalated cells, as a CO_2_ channel that promotes 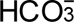 secretion.

Besides the above glandular tissues, AQP5 is present in astrocytes, where it may play a role during metabolic and traumatic injuries (Chai *et al*., 2013). In the eye, AQP5 is present in the epithelia layer (Funaki *et al*., 1998) and stromal keratocytes (Kumari *et al*., 2012) of the cornea, as well as in the epithelia l and fiber cells of the lens (Kumari *et al*., 2012). In the palmar epidermis, AQP5 is expressed in keratinocytes (Blaydon *et al*., 2013). In the aforementioned cells and elsewhere in the body, AQP5 could play a role in cell-volume regulation and osmo-sensing.

Governing CO_2_ diffusion across a membrane is Fick’s law, a simplified and integrated form of which was introduced by Wroblewski (1879):

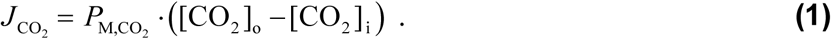

Here, *J*CO_2_ is flux across the membrane, the *P*_M,CO2_ is the macroscopic membrane CO_2_ permeability, and the term in parentheses indicates the transmembrane CO_2_ concentration gradient expressed in terms of bulk extracellular and intracellular.

For our purposes, it is more informative to consider events immediately adjacent to the membrane:

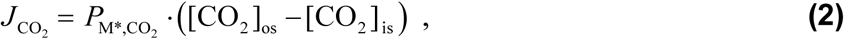

where [CO_2_]_os_ refers to a thin film of aqueous solution on the outer surface of the plasma membrane, [CO_2_]_is_ refers to a thin film on the inner surface, and *P*_M*,CO2_ (with an *) is the corresponding permeability.

Eqn (2) better reflects the reaction-diffusion mathematical models to which we refer above. Stated simply, *J*CO_2_ depends both on *P*_M*,CO2_, which can be augmented by certain AQPs and other protein channels, and factors that can influence the nanoscopic concentration gradient near the membrane. Among these latter factors are the carbonic anhydrases (CAs), enzymes that reversibly catalyze the hydration of CO_2_ and the dehydration of H_2_CO_3_ (for review, see Boone et al., 2014). Among the 12 enzymatica l ly active human (h) α-carbonic anhydrases (hCA), hCA II has one of the highest catalytic rates (see Purkerson and Schwartz, 2007). It is expressed broadly throughout the body, and is especially abundant in cells that engage in substantial CO_2_/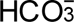 transport, including red blood cells, the renal proximal tubule, and alveolar type I cells (Chen *et al*., 2008). Moreover, CA II appears to play a role in CO_2_ elimination by the lung (Heming *et al*., 1986; Lien & Lai, 1998; Taki *et al*., 1999).

A long-established role of CAs is in the facilitated diffusion of CO_2_ within a single compartment (for review, see Occhipinti and Boron, 2019). In the case of transmembrane CO_2_ diffusion, when CA is present on both sides of an artificial membrane, the CA can replenish depleted CO_2_ on the membrane surface from which CO_2_ departs, and consume newly arriving CO_2_ on the opposite surface. The result is a magnifica t io n of the transmembrane CO_2_ concentration gradient on a nano-scale (see Eqn (2)), and thus an increase in transmembrane CO_2_ diffusion (Gutknecht *et al*., 1977). Work from our group on *Xenopus* oocytes—in the absence of exogenously expressed CO_2_ channels—has shown that hCA II (in the cytosol) and CA IV (mainly on the extracellular surface) each can enhance transmembrane CO_2_ fluxes, and that, together, the two enzymes produce a supra-additive stimulation (Musa-Aziz *et al*., 2014*a*, 2014*b*; Occhipinti *et al*., 2014)—synergism. Moreover, the group quantitatively accounted for these effects using computer simulations based on three-dimensional reaction-diffusion models. Left unanswered is the question of whether CAs can enhance the effect of CO_2_ channels (the *P*_M*,CO2_ term in Eqn (2)), and vice versa, on transmembrane CO_2_ fluxes. In the present study, we use simultaneous measurements of pH_S_ and intracellular pH (pH_i_) to ask whether hAQP5, cytosolic hCA II, and extracellular bovine CA (bCA; from erythrocytes) augment each other’s effects in accentuating CO_2_ fluxes across *Xenopus* oocyte plasma membranes. We find that the effects of expressing hAQP5 are supra-additive (i.e., synergistic) to those of adding CA to either the intra- or extracellular fluid. Moreover, in experiments in which we measure the increase in cell-surface pH (ΔpH_S_) caused by the influx of CO_2_, the introduction of CA to the cytosol augments the pH_S_ increase—a ΔΔpH_S_—by an amount that CO_2_ channels further augment—a ΔΔΔpH_S_. Thus, this ΔΔΔ approach could be a diagnostic tool for identifying candidate CO_2_ channels.

## Methods

### Ethical approval and animal procedures

The Institutional Animal Care and Use Committee at Case Western Reserve University approved all procedures for the housing and handling of *Xenopus laevis*. Housing, anesthesia and euthanasia were as described by Moss & Boron, (2020).

### Physiological solutions

Our ND96 solution comprises 93.5 mM NaCl, 2 mM KCl, 1 mM MgCl_2_, 1.8 mM CaCl_2_ and 5 mM HEPES (including ∼2.5 mM Na-HEPES after titration of solution pH to 7.50 using NaOH). We adjusted the osmolality to 195±5 mOsm by adding NaCl or H_2_O. Our 1.5% CO_2_/10 mM NaHCO_3_ solution was identical to ND96, except that 10 mM NaHCO_3_ replaced 10 mM NaCl, and we bubbled the solution with 1.5% CO_2_/balance O_2_. Our ND48—identical to ND96, except contained only 48 mM NaCl—was a hypotonic solution (∼100 mOsm) for the osmotic water permeability coefficient (*P*_f_) assay. The 0 Ca^2+^-normal saline (NRS) solution used during oocyte isolation was as described by Musa-Aziz *et al*., (2010).

To study the effect of extracellular carbonic anhydrase on CO_2_ diffusion, we dissolved bCA purified from bovine erythrocytes (Cat#C3934, Sigma-Aldrich, St. Louis, MO, USA)—a mixture of CA I and CA II—in both the ND96 and 1.5% CO_2_/10 mM NaHCO_3_ solutions of an experiment to achieve a concentration of 0.1 mg/ml (Musa-Aziz *et al*., 2014*b*).

OR3 medium as described by Musa-Aziz *et al*., (2010), comprises one sachet of Leibovitz L-15 (Cat#41300, ThermoFisher, Waltham, MA, USA), 100 ml Penicillin-Streptomycin (Cat#15140, ThermoFisher), and 1.785 g HEPES, all dissolved in ∼1.6 L of deionized H_2_O (dH_2_O), with pH titrated to 7.50 and osmolality adjusted to 195±5 mOsm by adding an appropriate volume of dH_2_O.

We confirmed all solution osmolalities with a Wescor model 5520 osmometer.

### Oocyte preparation

For a description of our approach, see Musa-Aziz et al. (2010) and Parker et al. (2008). Briefly, on Day 0, we anesthetized a frog (female *Xenopus laevis*, NASCO, Fort Atkinson, WI, USA) in 0.2% Tricaine (Cat#A5040, Sigma-Aldrich), surgically removed an ovary from the ventral side of the frog, cut the ovary into small pieces (∼0.5 × 0.5 × 0.5 cm), and washed the pieces ×3 with 0 Ca^2+^-NRS in a 50-ml Falcon tube, using a benchtop end-over-end rotator. We then digested the extracellular matrix of the follicles with 2 mg/ml collagenase (Cat#C5138, Sigma-Aldrich) dissolved in 0 Ca^2+^-NRS, washed ×3 in 0 Ca^2+^-NRS, carefully selected stage V and VI oocytes from the processed batch, and then cultured in OR3 medium in an incubator at 18 °C for subsequent injection of complementary RNA (cRNA).

### cDNA construct encoding human AQP5

The cDNA encoding human AQP5 (GenBank# NM_001651.4), subcloned into pGH19 vector, was the same construct as used by Qin and Boron (2012). After confirming the sequence, we linearized the cDNA using Xho I (Cat# R0146, New England Biolabs, Ipswich, MA, USA), transcribed it *in vitro* using a mMESSAGE mMACHINE^TM^ T7 capped RNA transcription kit (Cat#AM1344, ThermoFisher). We purified the cRNA using an RNeasy MinElute Cleanup Kit (Cat#74204, QIAGEN, Hilden, Germany), eluted the resulting cRNA with DNase/RNase-free H_2_O, quantified the cRNA concentration by measuring peak absorbance at λ=260 nm (A_260_) and assessed its purity by measuring the ratio of the λ=260 nm vs. λ=280 nm absorbance (A_260/280_) using a NanoDrop 2000 UV spectrophotometer (ThermoFisher), and diluted the cRNA into 1000 ng/μl aliquots, which we stored at –80°C.

### Expression in *Xenopus* oocytes

On Day 1, we injected oocytes with 25 ng (25 nl) of cRNA encoding hAQP5, or 25 nl of H_2_O as a control. On Day 4, we separated both cRNA-injected and H_2_O-injected oocytes into two subgroups, one pair of subgroups for injection with 25 nl containing 1 ng, 10 ng, or 100 ng of hCA II protein purified from human erythrocytes (Cat #C6165, Sigma-Aldrich) dissolved in “Tris” (i.e., 50 mM Tris base, titrated with HCl) at pH 7.40, and the other pair of subgroups for injection with 25 nl of “Tris” as a control.

### Electrophysiological measurements

#### Solution delivery to the chamber

At room temperature, we put each solution into two 140-ml piston syringes (Cat#8881114030, Covidien, Dublin, Ireland) and delivered the solution using a dual syringe pump (Cat#55-2226, Harvard Apparatus, Holliston, MA, USA) at a constant total flow of 4 ml/min. The solutions flowed through Tygon® tubing (4.76 mm OD × 1.59 mm ID, Cat#ACF00003, Saint-Gobain, Courbevoie, France) to minimize the leak of CO_2_. We used custom-manufactured 5-way valves (Clippard, Cincinnati, OH, USA) actuated by nitrogen pressure to switch solutions between ND96 and 1.5% CO_2_/10 mM NaHCO_3_. We controlled the nitrogen pressure using electrically activated 4-way solenoid valves (R481, Clippard).

#### Construction of microelectrodes

We made *V_m_* electrodes from borosilicate tubing (2.00 mm OD × 1.56 mm ID, with filament; Cat#BF200-156-10, Sutter Instrument, Novato, CA, USA), which we pulled on a Sutter puller (model P-97, Sutter Instrument), and filled with 3 M KCl, to achieve a resistance of 0.5 to 1.0 MΩ. For pH_i_ electrodes, after pulling, we dried the micropipettes at 270°C overnight, treated with *bis*-di-(methylamino)-dimethylsilane (Cat#14755, Sigma-Aldrich) for 20 min, vented the silane vapors, maintained the silanized micropipettes at 270°C overnight, removed them from the oven, filled the tip with H^+^ ionophore I-cocktail B (Cat#95293, Sigma-Aldrich), and then filled with backfill solution (see Musa-Aziz et al., 2010). We made pH_S_ electrodes in a fashion as pH_i_ electrodes, except that, after pulling the tubing (2.00 mm OD × 1.16 mm ID, without filament; Cat#B200-116-10, Sutter Instrument), we used a microforge to break the micropipette tip to give a final inner diameter of 30 μm. The thicker wall for the pH_S_ electrodes produces a better tip break, and the lack of the filament stabilizes the H^+^ cocktail.

### Measurement of pH_S_ and pH_i._

We measured pH_S_ and pH_i_ as previously described (Musa-Aziz *et al*., 2010). Briefly, with all electrodes (see below) in place in the plastic chamber, we placed an oocyte in the chamber, with flowing ND96; X-shaped nylon fibers downstream of the oocyte held the cell in place. We connected the *V*_m_ electrode to an OC-725C oocyte clamp (Warner Instruments, Hamden, CT, USA), and the pH_i_ and pH_S_ electrodes to a HiZ-223 amplifier (Warner Instruments). The electrical ground of the chamber was a bath clamp in which the same OC-725C as above connected to (a) a platinum wire in the chamber and (b) a reference electrode (identical to a *V_m_* electrode, but with its tip broken by dragging on paper) that contacted the ND96 downstream from the oocyte. The reference electrode for the pH_S_ measurements was a longer version of a *V*_m_ microelectrode, but with a carefully broken tip, placed downstream from the oocyte, and connected via a plastic electrode holder and calomel half-cell to a model 750 amplifier (WPI, Sarasota, FL, USA). All of the primary electrical signals fed into custom data-acquisition hardware “Ribbit Box” built in-house by Dale Huffman around a LabJack U6Pro device (LabJack, Lakewood, CO, USA) and connected to a Windows-based computer, controlled by custom software “The Frog Whisperer,” written in house by Dale Huffman.

Our sampling frequency was 3/s. We obtained the pH_i_ value by digitally subtracting (in the computer) the *V_m_* signal (voltage) from the pH_i_-electrode signal (voltage), and converted the difference to a pH_i_ value using the electrode-calibration data (see below). We similarly obtained the pH_S_ value by subtracting the signal of pH_S_ reference (calomel) from that of the pH_S_ electrode. The above custom software performed the calculations and displayed the record of all electrical parameters vs. time on a computer monitor.

### Calibration of pH electrodes

The first time we used a new pH_S_ or pH_i_ electrode—we never used an electrode for more than 1 day— we obtained the electrode slope by using the solution-delivery system (see above) to flow sequentially a pH 6.00 or pH 8.00 buffer (Certified, Cat#SB104-1 and SB112-1, ThermoFisher) through the chamber— before adding the oocyte—and measuring the voltage signal as described above. We accepted an electrode only if it had a slope of at least 55 mV/pH. We flushed the chamber with ND96 before adding the oocyte, and then performed a one-point calibration for each electrode in the ND96 (i.e., we assigned to the subtracted pH_S_ or pH_i_ electrode voltage the pH value of 7.50 in the ND96, and used the slope previously measured).

### Recording of electrophysiological data

In consecutive fashion, with ND96 flowing in the chamber, we impaled the oocyte with a *V_m_* electrode and a pH_i_ electrode, and then positioned the tip of the pH_S_ electrode ∼300 μm from the oocyte surface using a precision remote-controlled micromanipulator (Cat#ROE200, Sutter Instrument), all as previously described (Musa-Aziz *et al*., 2014*a*). While maintaining the flow of ND96, we advanced the pH_S_ electrode tip ∼300 μm to touch the oocyte surface, and an additional 40 μm to create a slight dimple on the oocyte surface. After 1 min, we switched the solution from ND96 to 1.5% CO_2_/10 mM NaHCO_3_. After several minutes, we retracted the pH_S_ electrode 340 μm from the oocyte surface for 1 min, in order to obtain a 1-point calibration of the pH_S_ electrode in the bulk 1.5% CO_2_/10 mM NaHCO_3_ solution. We then re-advanced the pH_S_ electrode 340 μm to resume pH_S_ measurements, before switching back to ND96. Finally, after pH_S_ had stabilized, we retracted the pH_S_ electrode 340 μm again, to obtain a 1-point calibration in the bulk ND96 solution at the end of the experiment.

### Calculation of ΔpH_S_

With its tip in the bulk ND96 solution, the pH_S_ electrode detected, by definition, a pH of 7.50 (i.e., the initial ND96 calibration). Once dimpling the membrane with the oocyte exposed to ND96, the pH_S_ electrode—calibrated for ND96—detected pH_S_. The average pH_S_ over at least the final 30 s—when pH_S_ was stable—we took as pH_S_(ND96_Init_). From the moment that we switched the chamber solution to 1.5% CO_2_/10 mM NaHCO_3_, the initial ND96 calibration of the pH_S_ electrode was no longer valid, which is why we obtained a new 1-point calibration in 1.5% CO_2_/10 mM NaHCO_3_ (see above); we applied this second CO_2_/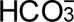 calibration to all pH_S_ data obtained during the CO_2_/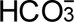 exposure, as pH_S_ rises to a peak and then decays downward. Using this second calibration, we computed the peak pH_S_ after the switch to CO_2_/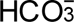, that is, pH_S_(CO_2,Peak_). We define the “upward” ΔpH_S_ for CO_2_ addition as pH_S_(CO_2,Peak_) – pH_S_(ND96_Init_). At the end of the CO_2_/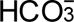 exposure, the average pH_S_ over at least the final 30 s—when pH_S_ was stable—we took as pH_S_(CO_2,T ail_). From the moment that we switched the chamber solution to ND96, the previous 1.5% CO_2_/10 mM NaHCO_3_ calibration of the pH_S_ electrode was no longer valid, which is why we obtained a final (i.e., third) 1-point calibration in ND96 (see above); we applied this final ND96 calibration to all pH_S_ data obtained during the final ND96 exposure, as pH_S_ falls to a trough and then decays upward. Using this third calibration, we computed the lowest pH_S_ after the switch to ND96, namely, pH_S_(ND96_Nadir_). We define the “downward” ΔpH_S_ for CO_2_ removal as pH_S_(ND96_Nadir_) – pH_S_(CO_2,T ail_), which is a negative number.

### Calculation of “maximal” dpH_S_/dt

During CO_2_/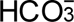 addition (e.g., see Figure 1C), pH_S_ rises rapidly from around 7.50 to a peak (where dpH_S_/dt ≅ 0) and then begins to decline, at first slowly (dpH_S_/dt is slightly negative), later rapidly (dpH_S_/dt reaches a maximally negative value), and finally more slowly again (dpH_S_/dt gradually becomes less and less negative) as pH_S_ relaxes towards an asymptotic value around 7.50. During CO_2_/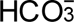 removal, the changes are in the opposite direction. Here we describe the analysis for CO_2_/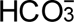 addition, but the approach is similar for CO_2_/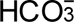 removal. Using software written in house by Dale Huffman, we obtain a running linear fit of (pH_S_, time) data points to generate a plot of (dpH_S_/dt)_Running_ vs. time, which allows us to identify the time (which we define as local time: t_Local_ = 0) of the maximally negative dpH_S_/dt (i.e., minimal dpH_S_/dt). We then obtain a double-exponential (DExp) fit of the (pH_S_, time) data points, beginning at t_Local_ = 0 and extending to the time when we removed the pH_S_ electrode from the cell surface to the bECF (for calibration). Finally, we evaluate the derivative of the DExp function at t_Local_ = 0, and take this value as (dpH_S_/dt)_Max_. During CO_2_/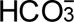 addition, this value is negative (i.e., maximal rate of pH_S_ descent), and during CO_2_/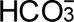 removal (e.g., see Figure 1*C*), this value is positive (i.e., maximal rate of pH_S_ rise).

**Figure 1.**
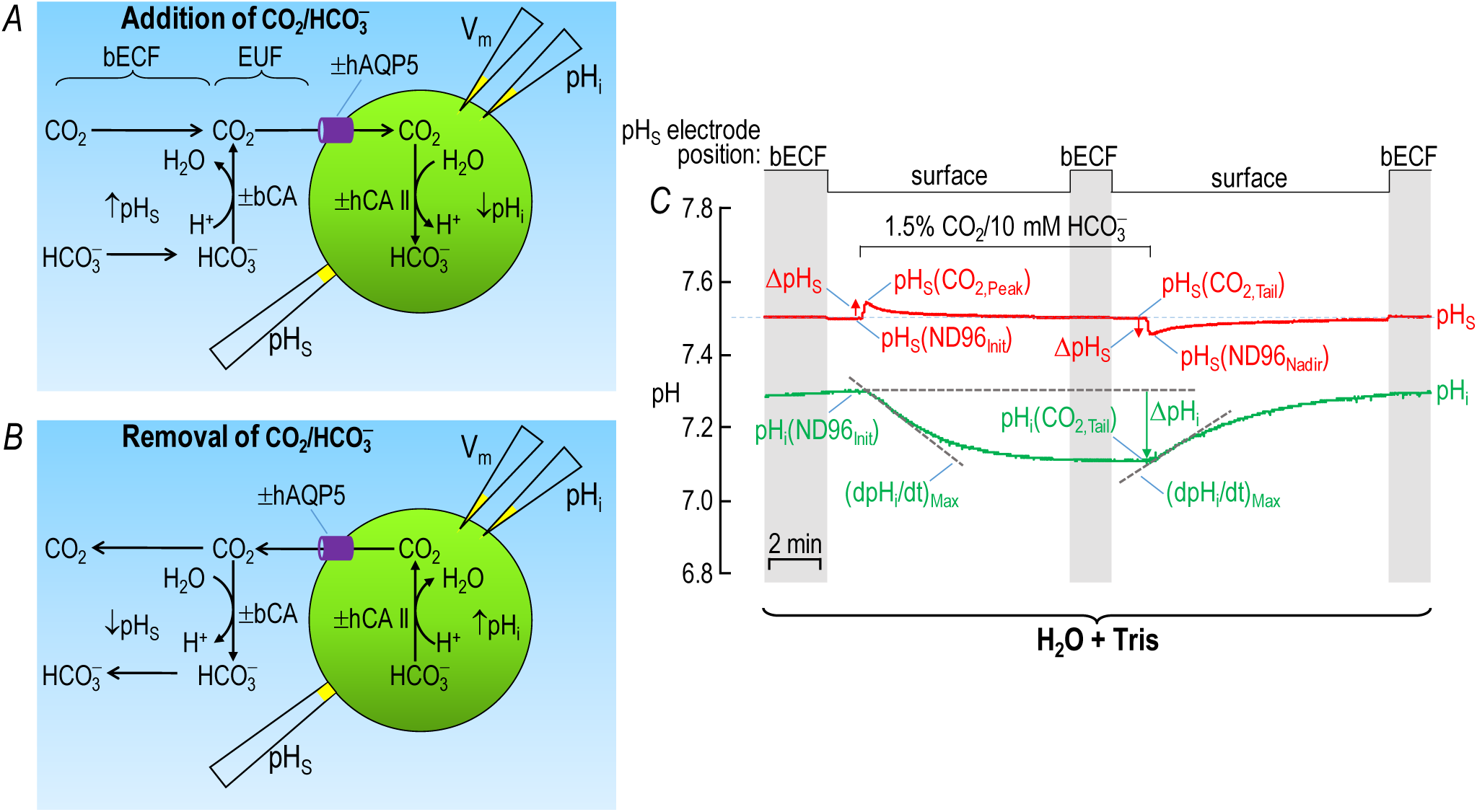
Design of experiments. *A*, model cell showing CO_2_ influx. The green circle represents a *Xenopus laevis* oocyte. Arrows with sharper arrowheads indicate the direction of diffusion. Arrows with blunter arrowheads indicate the direction of a chemical reaction. Upward and downward arrows associated with pH_S_ and pH_i_ indicate the direction of a pH change. The purple cylinder represents a human (h) AQP5 tetramer heterologously expressed in the membrane. We impale a *Xenopus laevis* oocyte with both a pH_i_ and *V_m_* electrode, and used a blunt, polished electrode to monitor pH_S_. ±, presence or absence; bECF, bulk extracellular fluid; bCA, extracellular bovine carbonic anhydrase (a mixture of CA I and CA II); EUF, extracellular unconvected fluid; hCA II, intracellular purified carbonic anhydrase II protein from human erythrocytes. *B*, Model cell showing CO_2_ efflux. All events occur in a direction opposite to that in *A*. *C*, Representative experiment. We dissected the oocyte on Day 0, injected it with H_2_O (as a control for cRNA encoding hAQP5) on Day 1, and then with “Tris” (as a control for hCA II protein) on Day 4. The stair-step line at the top of the panel indicates the position of the tip of the pH_S_ electrode. The vertical shaded bars indicate the time during which the pH_S_-electrode tip is in the bECF for recalibration at pH_o_ = 7.50; at other times, the pH_S_-electrode tip dimples the cell surface. The flowing solution is ND96 except for the period labeled “1.5 CO_2_/10 mM 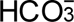.” The red record represents the time course of pH_S_, and the green record, the time course of pH_i_. We provide detailed definitions of the red and green labels in <u>Methods</u>.^15^

### Calculation of “maximal” dpH_i_/dt

After either the addition or removal of CO_2_/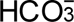, we define (dpHi/dt)_Max_ as the fastest change of pH_i_, which occurs several seconds after the new solution reaches the oocytes (Musa-Aziz *et al*., 2014*a*). For CO_2_/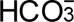 addition, pH_i_ decreases and thus the “downward” (dpH_i_/dt)_Max_ is a negative number; upon CO_2_/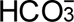 removal the “upward” (dpH_i_/dt)_Max_ is positive. After identifying, by eye, the time of fastest pH_i_ change (i.e., corresponding to the most extreme local value of dpH_i_/dt), we used Origin 2024 to obtain (dpH_i_/dt)_Max_ by a linear fit over the next ∼10 s of (pH_i_, time) data.

### Calculation of intrinsic intracellular buffering power

We define initial pH_i_ in ND96 solution—pH_i_(ND96_Init_)—as the average of at least 90 data points over a period (i.e., 30 s) when the experimenter (in real experimental time) judged pH_i_ to be stable, just before switching from ND96 to the CO_2_/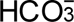 solution. Similarly, we defined the final pH_i_ in the CO_2_/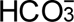 solution—pH_i_(CO_2_,_T ail_)—as the average of at least 90 data points over a period when the experimenter (in real experimental time) judged pH_i_ appeared stable, just before switching from the CO_2_/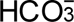 solution to ND96. During CO_2_/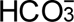 application, we define ΔpH_i_ (a negative number) as pH_i_(CO_2_,_Tail_) – pH_i_(ND96_Init_).

Based on the decrease in steady-state pH_i_ produced by the aforementioned application of CO_2_/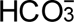, we calculated the intrinsic intracellular buffering (Boron, 1977) power β_I_ (mM/pH) of oocytes using the following equation:

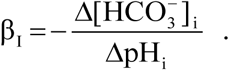

Here, the negative sign arises because we are observing a pH_i_ decrease produced by an intracellular, CO_2_-induced acid load. Because we impose the acid load by the introduction of CO_2_/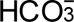, followed by the intracellular reaction CO_2_ + H_2_O → H^+^ + 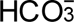, the magnitude of the acid load is Δ[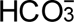] (for a discussion, see Roos and Boron, 1981). Because we assume that [HCO_3_^−^]_i_ is 0 before the exposure to CO_2_/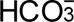, Δ[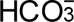] (a positive number) is the same as the final [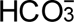] after the introduction of CO_2_/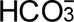, determined at a time when pH_i_ is stable. Because we observed no pH_i_ recovery during the CO_2_/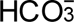 exposure, nor a pH_i_ overshoot after the removal of CO_2_/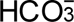, we can conclude that net acid extrusion during the CO_2_/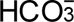 exposure was negligible (Boron & De Weer, 1976). Thus, we computed the final [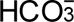] using pH_i_(CO_2_,_T ail_). Because we used 1.5% CO_2_/10 mM 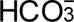 at a pH of 7.5,

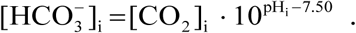

Here we assume that [CO_2_]_i_ = [CO_2_]_o_, where the subscript “o” denotes the bulk extracellular fluid, and that the *pK*_a_ in the intracellular fluid is the same as in the extracellular fluid.

### Measurement of Pf

After recording the pH_S_ and pH_i_, we saved the oocyte, equilibrated it in ND96 for at least 10 min, transferred it to a petri dish containing the hypotonic solution ND48 and a 1.6-mm diameter steel sphere (i.e., a ball bearing), placed the dish under a dissecting microscope (model Stemi 508, ZEISS, Oberkochen, Germany) equipped with a video camera (OptixCam summit series, Microscope LLC, Roanoke, VA, USA) connecting to a computer running proprietary software, and recorded 1 image/s of the swelling oocyte and the nearby steel sphere for 1 min. We used Image J software (U. S. National Institutes of Health, Bethesda, Maryland, USA, https://imagej.nih.gov/ij) to determine the perimeter of the oocyte at each time point, computed the projection area (using the steel sphere as an area standard), computed the oocyte volume (assuming the oocyte to be a sphere), and used the following equation to calculate osmotic water permeability:

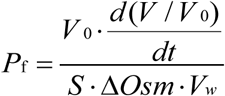

Here, *V*_0_ is the initial oocyte volume, *d*(*V*/*V*_0_)/*dt* is the rate of volume increase during the first minute, Δ*Osm* is the osmotic gradient across the membrane (i.e., 195 mOsm – 100 mOsm), V_W_ is the molar volume of water (18 ml/molar), and S is the oocyte surface area, assumed to be 8-fold greater than the idealized area (Chandy *et al*., 1997).

### Statistical analysis

We present data as mean ± SD. To compare the difference between 2 or more groups, we use a one-way ANOVA followed by the Tukey post-hoc analysis. We perform the analyses using Origin 2024, considering *P*<0.05 as significant.

To compare means of analyzed parameters of oocytes exposed to bECF lacking bCA (gray, black, light- and dark-blue bars in Figure 3 *A–D*) with oocytes for which the bECF contained bCA (Figure 8*A–D*), we perform a one-way ANOVA with Tukey’s means comparisons. In Statistics Tables 8*E–H*, we present descriptive statistics and means comparisons to determine the statistical significance of bCA-dependent differences.

**Figure 2.**
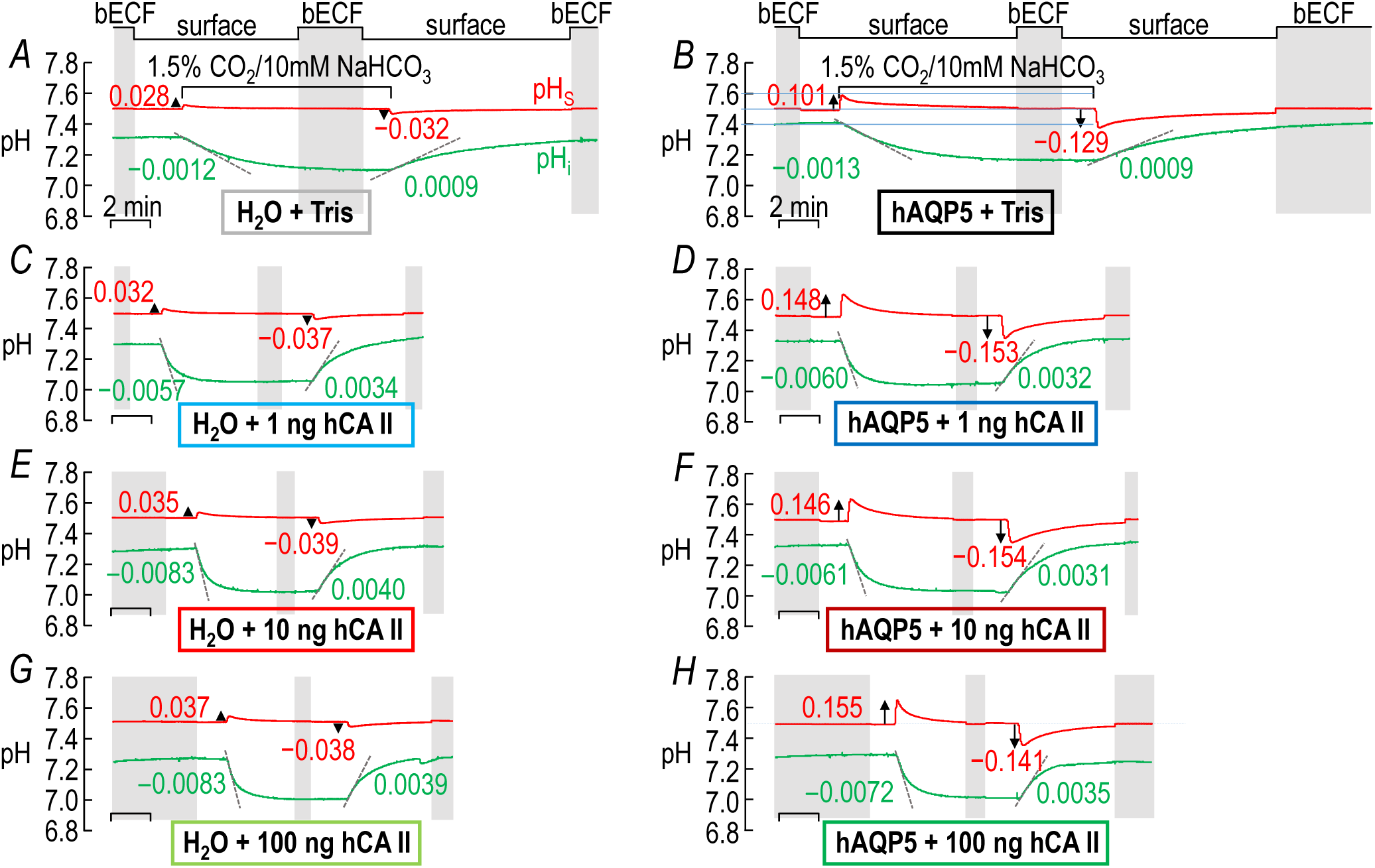
Representative pHS and pHi recordings: ±hAQP5, Δcytosolic hCA II, all in the absence of extracellular bCA (i.e., –bCA). *A*, “H_2_O + Tris”. We injected this oocyte with H_2_O (as a control for cRNA encoding hAQP5) on Day 1, and then with “Tris” (as a control for hCA II) on Day 4. *B*, “hAQP5 + Tris”. This oocyte, we injected on Day 1 with cRNA encoding hAQP5, and then on Day 4 with “Tris”. *C*, “H_2_O + 1 ng hCA II”. Similar to panel *A* except that on Day 4, we injected the oocyte with hCA II enzyme dissolved in “Tris”. *D*, “hAQP5 +1 ng hCA II”. Similar to panel *B* except that on Day 4, we injected the oocyte with hCA II enzyme dissolved in “Tris”. *E*, “H_2_O + 10 ng hCA II”. *F*, “hAQP5 + 10 ng hCA II”. *G*, “H_2_O + 100 ng hCA II”. *H*, “hAQP5 + 100 ng hCA II”. The “Δ” in the title and elsewhere in the paper implies that we choose among several levels of injected hCA II. The numbers in red are ΔpH_s_ and in green are (dpH_i_/dt)_Max_ for the oocyte presented here. The stair-step line at the top of *A* and *B* indicates the position of the pH_S_ electrode. The vertical shaded bars indicate times during which the pH_S_ electrode is in the bulk extracellular fluid (bECF) for recalibration; at other times, the pH_S_ electrode dimples the cell surface for actual pH_S_ measurements. The colors of the rectangles that surround the panel labels (e.g., “H_2_O + Tris”) correspond to the colors of the bars in Figure 3 and later figures.

**Figure 3.**
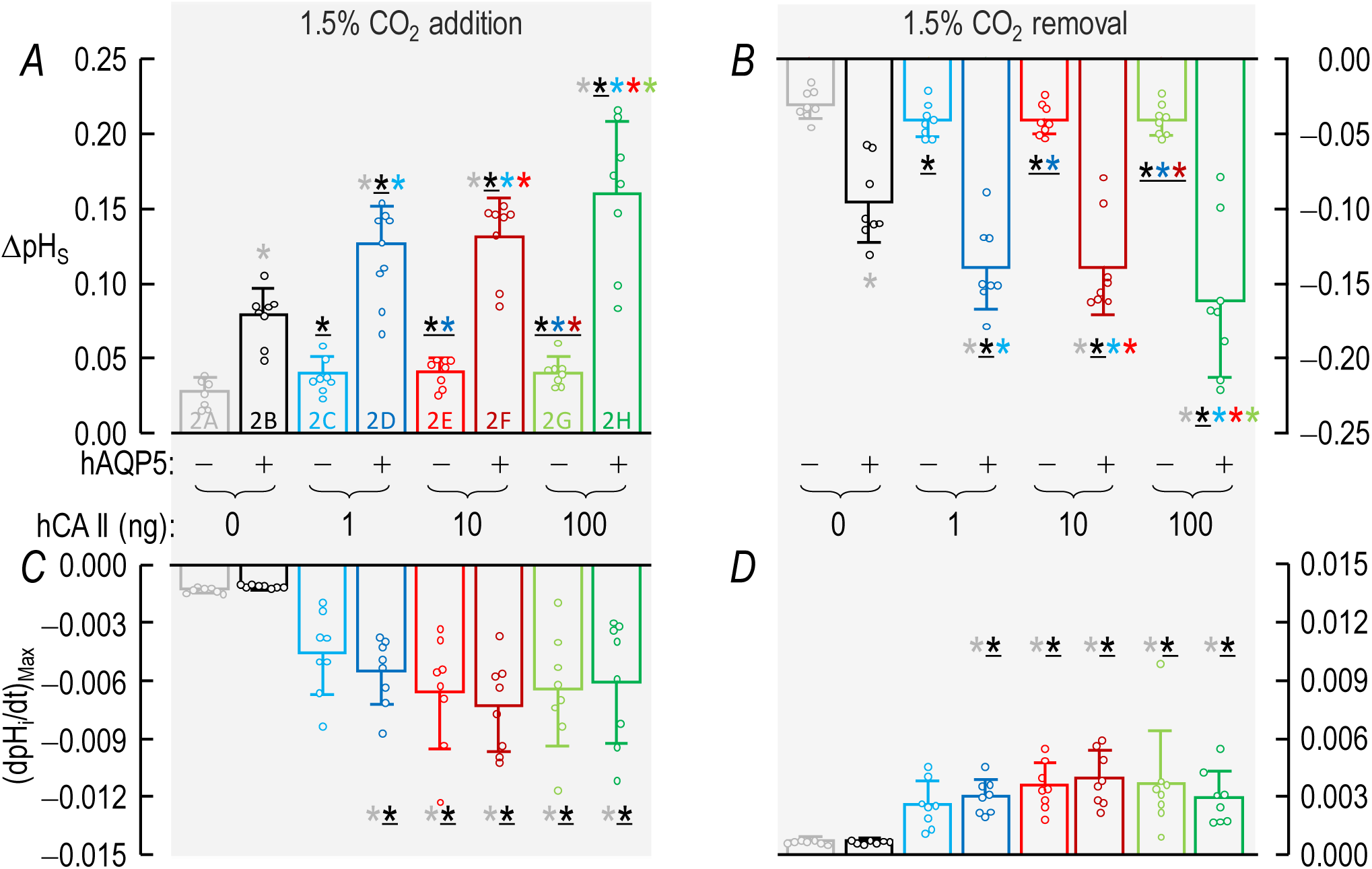
Summary of ΔpHS and (dpHi/dt)_Max_ data from experiments like those in Figure 2: ±hAQP5, Δcytosolic hCA II, all in the absence of extracellular bCA (i.e., –bCA). *A*, Summary of ΔpH_S_ upon addition of 1.5% CO_2_/10 mM 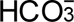. We computed individual ΔpH_S_ values as outlined in “Methods” > “Electrophysiological measurements” > “Calculation of ΔpH_S_”. *B*, Summary of ΔpH_S_ upon removal of 1.5% CO_2_/10 mM 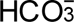. *C*, Summary of (dpH_i_/dt)_Max_ upon addition of 1.5% CO_2_/10 mM 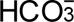. We computed individual ΔpH_S_ values as outlined in “Methods” > “Electrophysiological measurements” > “Calculation of “maximal” dpH_S_/dt”. *D*, Summary of (dpH_i_/dt)_Max_ upon removal of 1.5% CO_2_/10 mM 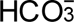. The “Δ” in the title and elsewhere in the paper implies that we choose among several levels of injected hCA II. Data are presented as mean ± SD. “−” and lighter-colored bars indicate that we injected oocytes with H_2_O; “+” and darker-colored bars, with cRNA encoding hAQP5 on Day 1. “0”, “1”, “10” and “100” indicate the amount of hCA II (ng) injected into oocytes on Day 4. Gray/black and light-/dark-blue, red, and green pairs of bars indicate 8 groups of oocytes. These colors correspond to the colors of the rectangles that surround panel labels (e.g., “H_2_O + Tris”) in Figure 2. A star associated with a bar indicates a statistically significant difference when comparing the bar to another bar with the same color as the star. The star/bar combinations are meant to be read right to left; we indicate statistical significance between two bars only once, associating the star with the bar further to the right. Thus, the mean value represented by the dark-green bar in panel *A* differs significantly from the means of the following bars: light gray, black (underscored to indicate darker than gray), light blue, light red, and light green. Conversely, the gray bar, which has no stars, differs significantly from the means of all bars with a gray star. The absence of a star indicates a lack of statistical significance. See Statistics Table 3 for *P*-values.

To compare mean values of various other parameters—initial pH_i_, ΔpH_i_, β_I_, and *P*_f_—from oocytes not exposed to bCA (gray, black, light- and dark-blue bars in Figure 6*A–D*) with their counterparts exposed to bCA (Figure 10*A–D* (bECF +bCA), we use the same approach described in the previous paragraph. In Statistics Tables 10*E–H*, we present descriptive statistics and means comparisons to determine the statistical significance of bCA-dependent differences.

**Figure 4.**
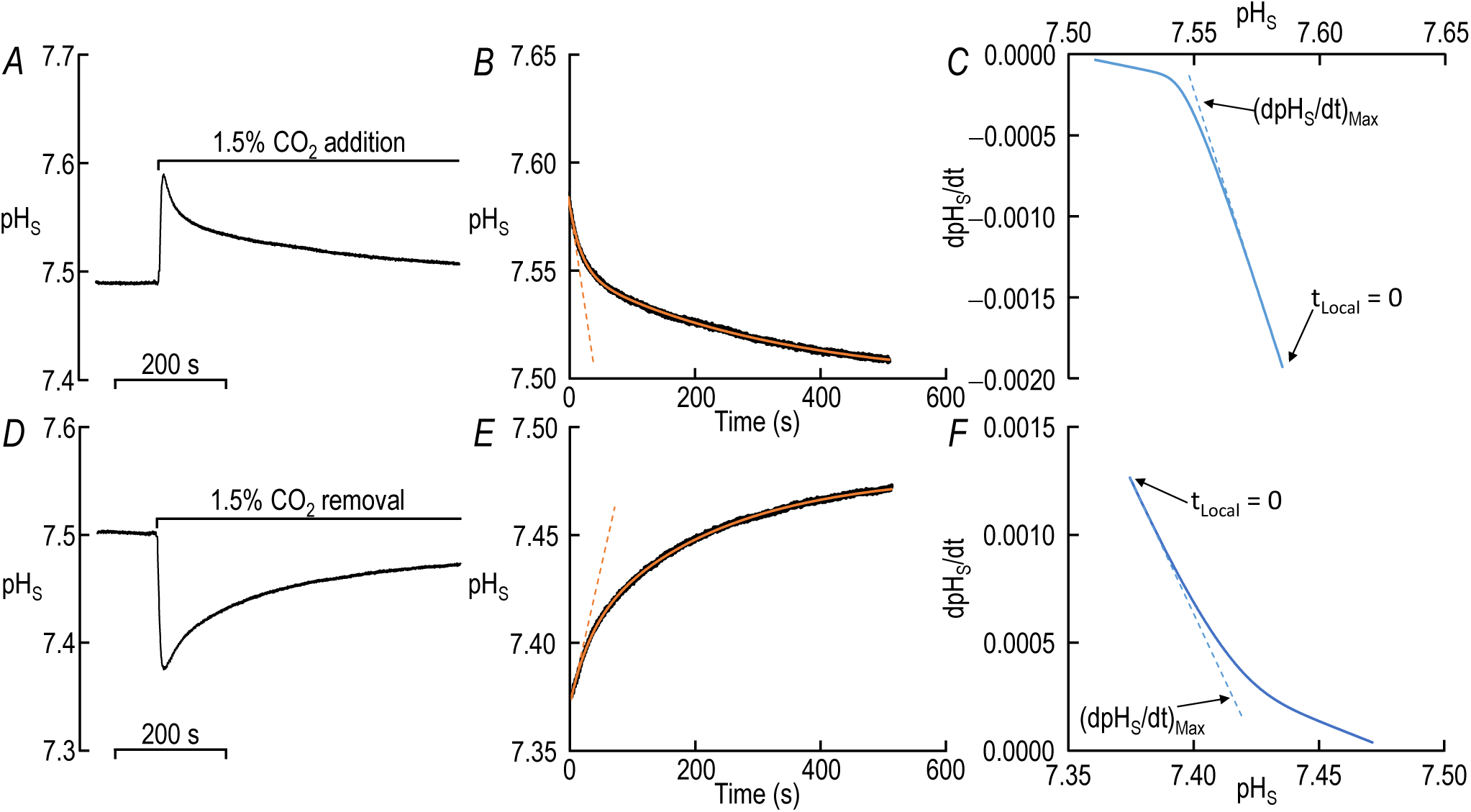
Analysis of (dpH_S_/dt)_Max_, the magnitude of maximal rate of relaxation of pH_S_. *A*, Time course of surface pH (pH_S_) during addition of 1.5% CO_2_/10 mM 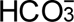. This is a reproduction—with a magnified y-axis—of the red record (during CO_2_ addition) in Figure 2*B*. *B*, Detail of panel *A*, showing only the pH_S_ relaxation, beginning from the time (t_Local_ = 0 s) of fastest pH_S_ descent. The jittery black record is the actual pH_S_ recording; the smooth orange curve is the result of a double-exponential (DExp) curve fit. The dashed red line has a dpH_S_/dt-vs.-time slope that is the derivative of the fitted DExp function at t_Local_ = 0 s. *C*, Dependence of dpH_S_/dt on pH_S_ during addition of CO_2_/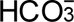 application. We obtained dpH_S_/dt by computing the derivative of the DExp best-fit function at the time that corresponds to the indicated pH_S_. Note that real time runs from right to left. The dashed blue line has a dpH_S_/dt-vs.-pH_S_ slope that corresponds to the value at t_Local_ = 0 s. A perfect single-exponential fit of pH_S_ vs. time would have produced a straight line in this dpH_S_/dt-vs.-pH_S_ plot. Thus, the break in the blue curve near pH_S_ = 7.54 is the demarcation between the dominance of a rapid/initial process and a slower/later process. *D*, Time course of pH_S_ during removal of 1.5% CO_2_/10 mM 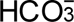. This is a reproduction—wi th a magnified y-axis—of the red record (during CO_2_ removal) in Figure 2*B*. *E*, Detail of panel *D*, processed as in panel *B*. *F*, Dependence of dpH_S_/dt on pH_S_ during CO_2_/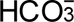 removal, processed as in panel *C*. Note that real time runs from left to right.

**Figure 5.**
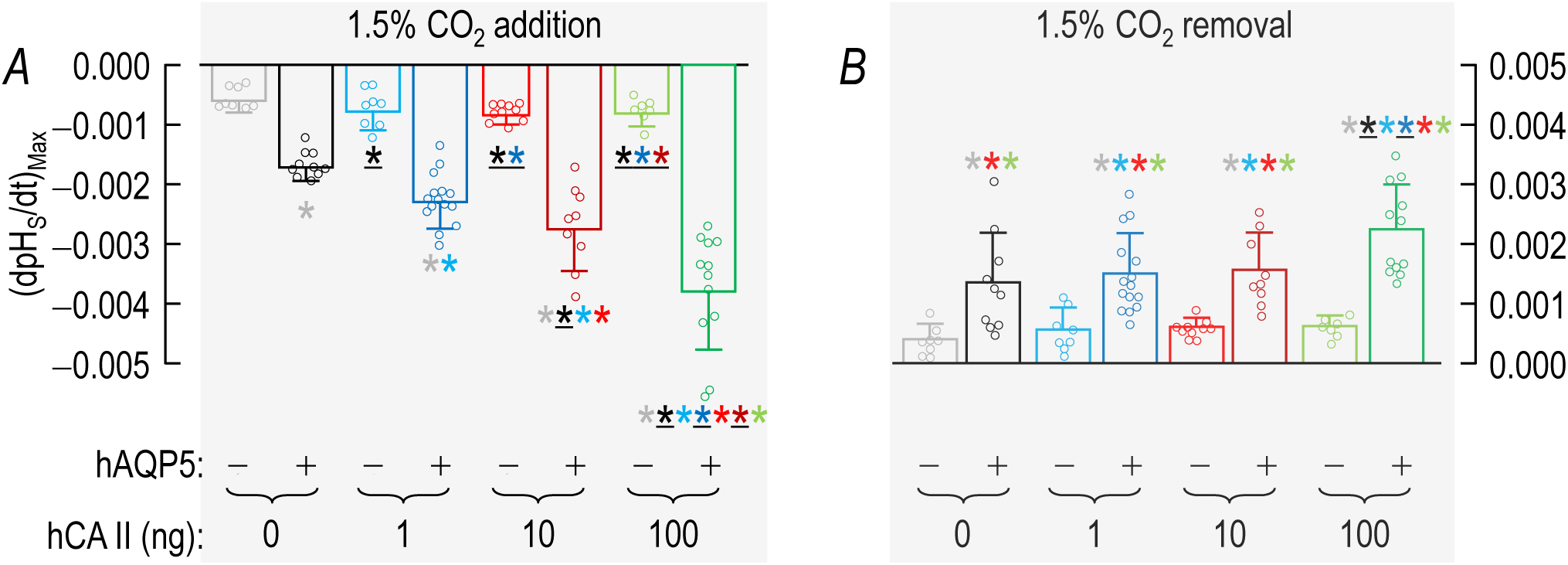
Summary of (dpHS/dt)_Max_ data from experiments like those in Figure 2: ±hAQP5, Δcytosolic hCA II, all in the absence of extracellular bCA (i.e., –bCA). *A*, Summary of (dpH_S_/dt)_Max_ upon addition of 1.5% CO_2_/10 mM 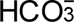. We computed individual (dpH_S_/dt)_Max_ values as described in Figure 4. *B*, Summary of (dpH_S_/dt)_Max_ upon removal of 1.5% CO_2_/10 mM 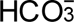. The “Δ” in the title and elsewhere in the paper implies that we choose among several levels of injected hCA II. Data are presented as mean ± SD. “−” and lighter-colored bars indicate that we injected oocytes with H_2_O; “+” and darker-colored bars, with cRNA encoding hAQP5 on Day 1. “0”, “1”, “10” and “100” indicate the amount of hCA II (ng) injected into oocytes on Day 4. Gray/black and light-/dark-blue, red, and green pairs of bars indicate 8 groups of oocytes. These colors correspond to the colors of the rectangles that surround panel labels (e.g., “H_2_O + Tris”) in Figure 2. A star associated with a bar indicates a statistically significant difference when comparing the bar to another bar with the same color as the star. The star/bar combinations are meant to be read right to left; we indicate statistical significance between two bars only once, associating the star with the bar further to the right. Thus, the mean value represented by the dark green bar in panel *A* differs significantly from the means of the following bars: light gray, black (underscored to indicate darker than gray), light blue, dark blue, light red, dark red, and light green. Conversely, the gray bar, which has no stars, differs significantly from the means of all bars with a gray star. The absence of a star indicates a lack of statistical significance. See Statistics Table 5 for *P*-values.

**Figure 6.**
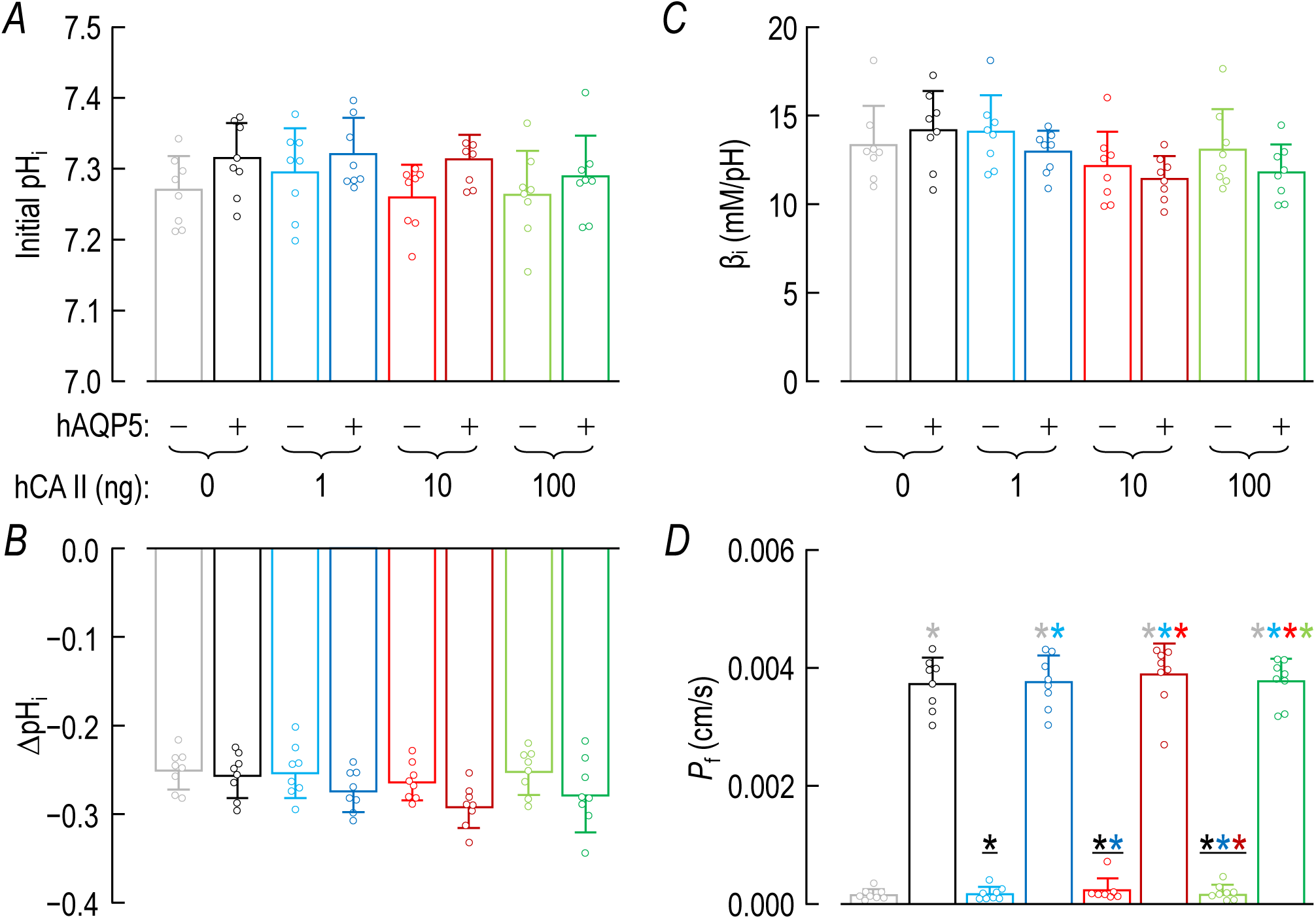
Summary of other oocyte parameters from experiments like those in Figure 2: ±hAQP5, Δcytosolic hCA II, all in the absence of extracellular bCA (i.e., –bCA). *A*, Summary of initial pH_i_ values. We computed individual initial pH_i_, ΔpH_i_ (see panel *B*), and β_I_ (see panel *C*) values as outlined in “Methods” > “Electrophysiological measurements” > “Calculation of intrinsic intracellular buffering power”. *B*, Summary of ΔpH_i_ elicited by addition of 1.5% CO_2_/10 mM 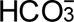. *C*, Summary of intrinsic buffering power (β_I_). *D*, Summary of *P*_f_. We computed individual initial pH_i_, ΔpH_i_ (see panel *B*), and β_I_ (see panel *C*) values as outlined in “Methods” > “Electrophysiological measurements” > “Measurement of *P*_f_”. The “Δ” in the title and elsewhere in the paper implies that we choose among several levels of injected hCA II. Data are presented as mean ± SD. “−” and lighter-colored bars indicate that we injected oocytes injected with H_2_O; “+” and darker-colored bars, with cRNA encoding hAQP5 on Day 1. “0”, “1”, “10” and “100” indicate the amount of hCA II (ng) injected into oocytes on Day 4. Gray/black and light-/dark-blue, red, and green pairs of bars indicate 8 groups of oocytes. These colors correspond to the colors of the rectangles that surround panel labels (e.g., “H_2_O + Tris”) in Figure 2. A star associated with a bar indicates a statistically significant difference when comparing the bar to another bar with the same color as the star. The star/bar combinations are meant to be read right to left; we indicate statistical significance between two bars only once, associating the star with the bar further to the right. Thus, the mean value represented by the dark green bar in panel *D* differs significantly from the means of the following bars: light gray, light blue, light red, and light green. Conversely, the gray bar, which has no stars, differs significantly from the means of all bars with a gray star. The absence of a star indicates a lack of statistical significance. See Statistics Table 6 for *P*-values.

In Figure 11 we determine the mean ΔΔpH_S,Base_ by subtracting the mean ΔpH_S_ for –hAQP5/–hCA II oocytes from each measured ΔpH_S_ of –hAQP5/+hCA II oocytes. We determine the mean ΔΔpH_S,hAQP5_ by subtracting the mean (ΔpH_S_) for +hAQP5/–hCA II oocytes from each measured ΔpH_S_ of +hAQP5/+hCA II oocytes. We then performed a one-way ANOVA followed by the Tukey post-hoc analysis on the data for ΔΔpH_S,Base_ vs. ΔΔpH_S,hAQP5_.

In Figure 12 we determine the mean Δ(dpH_i_/dt)_Max,Base_ by subtracting the mean (dpH_i_/dt)_Max_ for – hAQP5/–bCA oocytes from each measured (dpH_i_/dt)_Max_ of –hAQP5/+bCA oocytes. We determine the mean Δ(dpH_i_/dt)_Max,hAQP5_ by subtracting the mean Δ(dpH_i_/dt)_Max_ for +hAQP5/–bCA oocytes from each measured Δ(dpH_i_/dt)_Max_ of +hAQP5/+bCA oocytes. We then perform a one-way ANOVA followed by the Tukey post-hoc analysis on the data for Δ(dpH_i_/dt)_Max,Base_ vs. Δ(dpH_i_/dt)_Max,hAQP5_.

## Results

### General protocol

Figure 1A and B are schematic representations suggesting how hAQP5, cytosolic hCA II, and extracellular ^b^C^A^, would affect the influx (panel *A*) or efflux of CO_2_ (panel *B*) and thus pH_S_ and pH_i_ of an *Xenopus* oocyte. Imagine that we switch the solution—the bulk extracellular fluid (bECF^1^)—that flows through our chamber from one that nominally lacks, to one that contains CO_2_/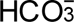. Imagine also that we have a cell whose membrane is initially impermeable to CO_2_ and 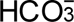. Thus, a short time after we switch solutions, CO_2_ and 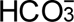 will have diffused throughout the system so that [CO_2_] and [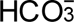] will be uniform through the bECF (an “infinite reservoir”) and extracellular unconvected fluid (EUF), including a thin layer of fluid at the oocyte surface (S). Suddenly, we now increase *P*_M,CO2_, which is where Figure 1A picks up the narrative. Now CO_2_ diffuses into the cell (Figure 1A), leading to the depletion of CO_2_ at the cell surface. The replenishment of this surface CO_2_ occurs via both CO_2_ diffusion from bECF and— especially adjacent to the cell surface—the reaction 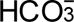 + H^+^ → CO_2_ + H_2_O. The reaction causes a rise in pH_S_, the maximum magnitude of which is ΔpH_S_; assuming that cell-surface CA activity is fixed, ΔpH_S_ is a semiquantitative index of the CO_2_ influx (see Eqn (2)). Meanwhile, beneath the membrane, the CO_2_ that has entered the cell undergoes the reaction CO_2_ + H_2_O → H^+^+ 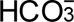, thus causing a fall in pH_i_. The maximal rate of intracellular acidification—assuming that cytosolic CA activity is fixed—is also an index of the CO_2_ influx. However, (dpH_i_/dt)_Max_ is a far less sensitive measure than ΔpH_S_ (Musa-Aziz *et al*., 2014*b*).

If we nominally remove CO_2_/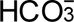 from the solution flowing through the chamber, all of the above processes, including pH_S_ and pH_i_ changes reverse (Figure 1B).

Figure 1C shows typical pH_S_ and pH_i_ records of a control oocyte—that is, one injected with H_2_O (rather than cDNA dissolved in H_2_O encoding hAQP5) on Day 1, and then “Tris” (rather than hCA II dissolved in “Tris”) on Day 4—during the application and removal of CO_2_/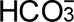. No bCA is present in the bECF. The shaded vertical bars represent periods of time during which the pH_S_ electrode is ∼300 μm from the cell surface, in the bECF, for electrode calibration. At other times (i.e., between shaded bars), the electrode tip dimples the oocyte surface. Switching the solution from ND96 to 1.5% CO_2_/10 mM NaHCO_3_ causes an upward pH_S_ transient that soon reaches a maximum (left upward red arrow), and a pH_i_ decrease, the maximum rate of which is indicated by the left dashed line. The pH_S_ signal (after the pH_S_ peak) relaxes and pH_i_ signal declines with similar time constants (Musa-Aziz *et al*., 2014*a*, 2014*b*; Occhipinti *et al*., 2014). Later, we switch the solution back to ND96, which elicits a reversal of the previous events. In Methods and the figure legend, we define the red and green labels shown in Figure 1C.

In the following eight sections, we describe the effects—on various parameters measured during our standard protocol in Figure 1C—of various combinations of human AQP5 expressed or not (±hAQP5), human CA II injected or not (±hCA II), and bovine CA added to the bulk extracellular fluid or not (±bCA). In the first four sections, we examine oocytes in the absence of bCA, but with various combinations of ±hAQP5 and ±hCA II. In the second group of four, we study oocytes in the presence of extracellular bCA.

### ±hAQP5^2^ in absence of exogenous CAs: Effects on ΔpHS & (dpHi/dt)Max

Our overall goal is to investigate the relative roles of hAQP5, cytosolic hCA II, and extracellular bCA in promoting CO_2_ fluxes across the plasma membrane of a *Xenopus* oocyte. In this first section, we begin by verifying previous data that addressed the role of hAQP5 in promoting CO_2_ diffusion in the absence of added CAs. We injected oocytes on Day 1 with either “H_2_O” or H_2_O + cRNA encoding “hAQP5” and then, on Day 4, with “Tris” (as a control for “Tris + hCA II”).

**pH_S_** The red record in Figure 2A shows the pH_S_ data for a representative “H_2_O + Tris” oocyte. The red record in Figure 2B shows the comparable data for an “hAQP5 + Tris” oocyte. Consistent with previous results (Musa-Aziz *et al*., 2009; Geyer *et al*., 2013), the expression of hAQP5 leads to pH_S_ transients that are substantially greater in magnitude—both with the addition of CO_2_/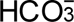 (0.101 vs. 0.028) and with the removal of CO_2_/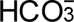 (–0.129 vs. –0.032)—than the mere injection of H_2_O into the oocytes (red records in Figure 2A vs. *B*).

*Summary.* For a larger number of these oocytes studied in the absence of both hCA II and bCA, the bars in Figure 3A provide the mean ΔpH_S_ data for CO_2_/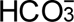 addition in the absence (gray) and presence (black) of hAQP5, respectively. The gray and black bars in Figure 3B summarize the corresponding data for CO_2_/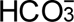 removal.

*Conclusions.* From the statistical analysis of the data summarized in Figure 3A*,B*, we conclude that— in the absence of both hCA II and bCA, both for CO_2_/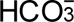 addition and removal—expression of hAQP5 increases the magnitude of ΔpH_S_.

*Interpretation: ΔpH_S_ (gray vs. black bars) in Figure 3A,B; ±hAQP5, –hCA II, –bCA.* Expression of hAQP5 increases *P*_M,CO2_ in Eqn (1).

**dpH_i_/dt** The green records in Figure 2A and B show the pH_i_ data that correspond to the pH_S_ data presented above. The expression of hAQP5 does not produce a remarkable change in either the downward or upward (dpH_i_/dt)_Max_.

*Summary.* The gray and black bars in Figure 3C show the mean downward (dpH_i_/dt)_Max_ for CO_2_/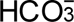 addition, ±hAQP5—in the absence of both hCA II and bCA. The bars in Figure 3D show the analogous data for CO_2_/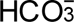 removal.

*Conclusions.* With either CO_2_/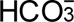 addition or removal—in the absence of injected hCA II and extracellular bCA—the expression of hAQP5 does not significantly affect (dpH_i_/dt)_Max_.

*Interpretation: (dpH_i_/dt)_Max_ (gray vs. black bars) in Figure 3C,D; ±hAQP5, –hCA II, –bCA.* (1) Expression of hAQP5, even though it markedly increases *P*_M,CO2_ as indicated by the ΔpH_S_ signals in Figure 3A*,B*, does not increase the magnitude of (dpH_i_/dt)_Max_. The reason, in part, is that—without hCA II—the cytosolic CO_2_ hydration reaction (during CO_2_ influx) and dehydration reaction (during CO_2_ efflux) are rate limiting, thereby limiting the transmembrane CO_2_ gradient and choking the CO_2_ fluxes (see Eqn (2)). In the second half of Results, we will see that the availability of CO_2_ on the outer surface of the cell (replenished/consumed by bCA) also is rate limiting under the conditions of Figure 2A*,B*. (2) Thus, (dpH_i_/dt)_Max_ is a relatively insensitive indicator of CO_2_ fluxes in the absence of bCA in the bECF. And (3) a contributing factor may be the large diameter of oocytes (∼1.2 mm); a prolonged time for diffus io n to/from the center of the cell limits the dynamic range of (dpH_i_/dt)_Max_. Note that, in the hands of Nakhoul et al. (1998) and Cooper & Boron, (1998), AQP1 expression did not produce a statistically significant effect on (dpH_i_/dt)_Max_ under similar experimental conditions (i.e., in the absence of injected CA enzyme, and with vitelline membrane intact).

### ±hAQP5, ΔhCA II, –bCA^3^: Effects on ΔpHS & (dpHi/dt)Max

To explore the interaction between hAQP5 and intracellular CA (CA_i_) in the diffusion of CO_2_, we injected oocytes on Day 1 with either “H_2_O” or “H_2_O + cRNA encoding hAQP5” and then, on Day 4, we injected oocytes with either “Tris” as a control, or “Tris + hCA II” at hCA II levels of 1, 10 or 100 ng.

**pHs** The red record in Figure 2C shows the pH_S_ data for a representative “H_2_O + 1 ng hCA II” oocyte and, in Figure 2D, the comparable data for an “hAQP5 + 1 ng hCA II” oocyte. Comparing the pH_S_ records between Figure 2C vs. Figure 2A, we see that—in the absence of hAQP5—the addition of 1 ng hCA II appears to increase the magnitude of the pH_S_ transient slightly. Musa-Aziz et al. (2014a) had previously observed that hCA II—presumably with a higher specific activity—significantly increases the magnitude of ΔpH_S_ with both CO_2_/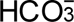 addition and removal. If we compare Figure 2D vs. Figure 2B, we see that—in the presence of hAQP5—the injection of 1 ng hCA II has a much larger effect on the pH_S_ transient. Finally, comparing Figure 2D vs. Figure 2C, and Figure 2B vs. Figure 2A, we see that the hAQP5 expression has a far greater effect on augmenting the magnitude of ΔpH_S_—both with CO_2_/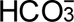 addition and removal—with the injection of 1 ng hCA II than with no added hCA II.

Note that, in the above comparisons, the pH_S_ electrode is “trans”—or on the opposite side of the membrane—with respect to the hCA II that we added to the cytosol, a concept developed by Musa-Aziz et al. (2014a, 2014b) and Occhipinti et al. (2014). In such cases, one can conclude by intuition—b ut supported by mathematical simulations—that the trans-side increase in ΔpH_S_ is indicative of an increased CO_2_ flux when hCA II is present in the cytosol. Thus, based on Figure 2A*–D*, we can conclude that both the expression of hAQP5 alone (a large effect) and the injection of hCA II alone (a small effect), and especially the two in combination, increase CO_2_ fluxes into/out of oocytes.

Examining the effects of injecting larger amounts of hCA II, we continue to see—in the absence of hAQP5—very little effect on the pH_S_ transients (Figure 2E*,G*) compared to the “Tris” control (Figure 2A). In the presence of hAQP5, the higher levels of injected hCA II (Figure 2F*,H*) continue to enhance the pH_S_ transients greatly compared to the “AQP5 + Tris” control (Figure 2B), but the stimulatory effect is little more than with our 1 ng dose of hCA II (Figure 2D).

In their earlier work, Musa-Aziz et al. (2014a) observed a baseline ΔpH_S_ (i.e., no injected CA) of slightly more than 0.04, whereas in the present study, our baseline ΔpH_S_ is only ∼0.03. Thus, the baseline values of CO_2_ permeability, surface CA activity, or cytosolic CA activity in that previous study may have been somewhat greater than in the present study. Moreover, Musa-Aziz et al. (2014a) found that injecting 300 ng of recombinant hCA II approximately doubled the ΔpH_S_, whereas in the present study injecting 100 ng hCA II enzyme (a dose at which the ΔpH_S_ seemingly had already plateaued) increased ΔpH_S_ only by about 1/3 (compare Figure 2G vs. *A*). We presume that our commercially obtained hCA II, purified from red blood cells (RBCs), had a lower specific activity than the recombinant hCA II in the previous study. We abandoned attempts at injecting >100 ng/oocyte because these higher doses seemed to have deleterious effects on the oocytes, perhaps because of contaminants.

*Summary.* The lighter-colored blue, red and green bars in Figure 3A display the mean ΔpH_S_ data for CO_2_/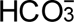 addition with increasing doses of hCA II, all in the absence of hAQP5 and bCA. The darker bars in Figure 3A summarize comparable data, but in the presence of hAQP5. For CO_2_/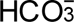 removal, Figure 3B reveals patterns that are similar to those in Figure 3A.

*Conclusions.* For oocytes examined in the absence of bCA, both for the addition and removal of CO_2_/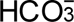: (a)^4^ Expressing hAQP5 (darker vs. lighter bars) increases the magnitude of ΔpH_S_ at every level of injected hCA II. (b) In the absence of hAQP5 (lighter bars), injecting oocytes with increasing amounts of hCA II, despite a modest upward trend, does not have a statistically significant effect on ΔpH_S_ magnitudes, neither for CO_2_/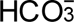 addition nor removal. And (c) in the presence of hAQP5 (darker bars), injection of increasing amounts of hCA II tends to cause graded increases of ΔpH_S_ magnitudes, both for CO_2_ influx and efflux. Relative to no injected hCA II, the effects reach statistical significance at all three levels of injected hCA II, although these three bars are not significantly different from each other.

*Interpretation: ΔpH_S_ (colorful bars) in Figure 3A,B; ±hAQP5, ΔhCA II, –bCA.* (1) In the absence of hAQP5 (4 lighter bars), *P*_M,CO2_ is the major rate-limiting factor in both directions of CO_2_ diffusion, and at all hCA II levels. Expression of hAQP5 augments *P*_M,CO2_ and thereby increases ΔpH_S_ (4 darker bars). (2) During CO_2_/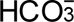 addition in control cells (–hAQP5, –hCA II; gray bars), disposal of incoming CO_2_ (in this case by cytosolic CA) is borderline rate limiting (i.e., with greater CA_i_ activity, the flux would have been modestly greater), as noted previously by Musa-Aziz et al. (2014a). Conversely, during the subsequent CO_2_/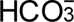 removal, replenishment of outgoing CO_2_ (here, by hCA II), likewise, is borderline rate limiting. And (3) as we will see below in our presentation of data on bCA in the bECF, for control cells (gray bars), the replenishment of CO_2_ on the extracellular surface is also rate limiting during CO_2_ influx. Conversely, disposal of exiting CO_2_ at the cell surface during CO_2_ efflux is also rate limiting. (4) The greater effect of expressing hAQP5 in oocytes injected with 1 ng of hCA II (light-vs. dark-blue bars) than in oocytes injected only with “Tris” (gray vs. black bars) is an example of synergism.

**dpH_i_/dt** The green records in Figure 2C*,E,G and D,F,H* show the pH_i_ data that correspond to the pH_S_ data presented above, with incremental amounts of injected hCA II, without hAQP5 (left side of Figure 2) and with hAQP5 (right side). We observe that injecting even 1 ng hCA II into the cytosol produces a striking increase in (dpH_i_/dt)_Max_, both for CO_2_/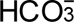 application and removal. Increasing the injected hCA II to 10 ng further increases the downward (dpH_i_/dt)_Max_ but has only a modest effect on the upward (dpH_i_/dt)_Max_ in this example. Increasing injected hCA II to 100 ng has no additional effect. Comparing the left and right sides of Figure 2, we see that hAQP5 expression seemingly has little effect on downward or upward (dpH_i_/dt)_Max_, regardless of the amount of injected hCA II.

Note that, in the above (dpH_i_/dt)_Max_ analyses, the pH_i_ electrode is “cis”—or on the same side of the membrane—with respect to the hCA II that we added to the cytosol, again, a concept developed by Musa-Aziz et al. (2014a, 2014b) and Occhipinti et al. (2014). In such cases, one cannot use intuition to arrive at conclusions that the cis-side acceleration of a pH_i_ change by hCA II is indicative of an increased CO_2_ flux. The reason is that—near the intracellular surface of the plasma membrane—the added hCA II greatly accelerates H^+^ formation during CO_2_ influx, and H^+^ consumption during CO_2_ efflux. The pH_i_ electrode directly senses the changes in [H^+^], which only indirectly reflect CO_2_ fluxes. However, previous mathematical simulations predict that the CO_2_ fluxes must have increased under these conditions (Musa-Aziz et al. (2014a) and Occhipinti et al. (2014).

*Summary.* The lighter/darker pairs of bars (–/+ hAQP5) in Figure 3C*,D* précis the mean (dpH_i_/dt)_Max_ data for CO_2_/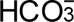 addition and removal.

*Conclusions.* Both for CO_2_/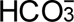 addition and removal: (a) Expression of hAQP5 does not significantly affect (dpH_i_/dt)_Max_ at any level of injected hCA II (4 sets of lighter vs. darker bars). And (b) in the absence of hAQP5 (4 lighter bars), increasing amounts of injected hCA II cause (dpH_i_/dt)_Max_ values to trend toward greater magnitudes. However, the effects do not reach statistical significance (vs. 0 ng) until 10 and 100 ng; the lighter blue, red, and green bars are not significantly different from one another.

*Interpretation: (dpH_i_/dt)_Max_ (colorful bars) in Figure 3C,D; ±hAQP5, ΔhCA II, –bCA.* (1) Because the injected hCA II is “cis” to the pH_i_ electrode, we cannot draw intuitive conclusions about CO_2_ fluxes as we compare the gray (dpH_i_/dt)_Max_ bar (i.e., –hAQP5) to the other three light-colored bars (i.e., increasing amounts of hCA II), or as we compare the black bar (i.e., +hAQP5) to the other three darker-colored bars. (2) Expression of hAQP5 (compare lighter-vs. darker-blue, red, and green bars) does not increase (dpH_i_/dt)_Max_ magnitudes. As noted in our interpretation of ΔpH_S_ data <u>immediately above</u>^5^, this lack of effect probably reflects limited availability of CO_2_ at the extracellular surface during influx and disposal of CO_2_ at the extracellular surface during efflux. Such choking could be mitigated by introducing an extracellular carbonic anhydrase (CA_o_), as we will see below as we discuss the gray vs. black bars in Figure 8C. Also <u>already mentioned</u>^6^, a compounding factor may be the large diameter of the oocyte. (3) Nevertheless, we know that expression of hAQP5 leads to a sizeable increase of *P*_M,CO2_ because of the parallel ΔpH_S_ data summarized by the lighter/darker pairs of bars in Figure 3A*,B*.

### ±hAQP5, ΔhCA II, –bCA: Effects on pHS relaxation

*Theoretical considerations.* Let us assume that the oocyte is a sphere, containing only an aqueous pH buffer, surrounded by a membrane permeable only to CO_2_, and initially devoid of CO_2_/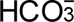. If we now expose the oocyte to a CO_2_/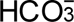 solution, we could compute—given [CO_2_]_o_, initial pH_i_, buffering power, and cell volume—the net number of CO_2_ molecules that would diffuse into the cell before the system came into equilibrium. The speed of this equilibration depends on the parameters that contribute to the description of Fick’s law in Eqns (1) and (2), including [CO_2_]_os_, [CO_2_]_is_, and *P*_M*,CO2_.

In their study of the impact of cytosolic CA II and CA IV (predominantly extracellular) on CO_2_ equilibration, Musa-Aziz et al. (2014a, 2014b) noted that, following the rapid upswing in pH_S_ triggered by CO_2_/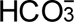 application (or the rapid downswing in pH_S_ triggered by CO_2_/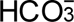 removal), (1) pH_S_ decayed following an approximately single-exponential (SExp) time course, and that the time constant (τ) of this decay decreased markedly either with (2) injection of recombinant hCA II into the cytosol (see fig. 13 in Musa-Aziz et al. (2014a) or (3) expression of hCA IV (see fig. 21 in Musa-Aziz et al. (2014b). Moreover, mathematical simulations based on a reaction-diffusion model (Somersalo *et al*., 2012) corroborated nearly all of the essential observations (Occhipinti *et al*., 2014). In other words, CA II and CA IV increased the speed (reflected by the rate constant = 1/τ) of CO_2_ equilibration, and thus 1/τ is an indirect measure of the transmembrane CO_2_ flux.

In the present study, we note that the pH_S_ relaxations do not so nearly approximate an SExp decay as in the work by Musa-Aziz et al. (2014a, 2014b), probably due to (1) minor differences in the tip of the pH_S_ electrodes, (2) the angles with which the pH_S_ electrode contacted the cell surface, and (3) the precise location of the pH_S_ electrode on the oocyte surface (see figure 1 insets in Musa-Aziz et al. (2014a). Note also that the mathematical simulations suggest that the pH_S_ decays should not be perfectly exponential in the first place (Musa-Aziz *et al*., 2014*a*, 2014*b*; Occhipinti *et al*., 2014). Therefore, in the present study, as a surrogate for 1/τ, we chose to use the maximal initial rate of pH_S_ relaxation.

*Exemplar data.* Figure 4A reproduces the pH_S_ time course for CO_2_/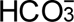 addition in Figure 2B (i.e., “hAQP5 + Tris”). The jittery black record in Figure 4B shows the time course of the actual pH_S_ relaxatio n, beginning at a local time (t_Local_) of 0, which we judged by eye to be time of fastest pH_S_ decay. The smooth orange curve is the result of a double-exponential (DExp) curve fit. The dashed orange line is the tangent to the DExp function, evaluated at t_Local_ = 0. The blue curve in Figure 4C is a plot of dpH_S_/dt, computed as the derivative of the best-fit DExp function, versus pH_S_. Here, time flows from right to left. The break in slopes at pH_S_ ≅ 7.54 is the result of the second, slower exponential process. The dashed blue line is the derivative of the DExp function, again calculated at t_Local_ = 0, but now plotted as a function of pH_S_. Although the choice of t_Local_ = 0 is subject to human error, the effect of misjudgment is likely to be minima l inasmuch as the DExp fit encompasses so many points, and because both the initial time course and the initial plot of dpH_S_/dt vs. pH_S_ are nearly linear.

Figure 4*D–F* are analogous to the top row of panels, except that here we analyze the CO_2_/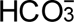-removal step in Figure 2B. Note that all of the pH_S_ records are inverted, and that, in Figure 4F, time runs from left to right.

*Summary.* The layout of the synopses in Figure 5*A,B* for (dpH_S_/dt)_Max_ data—CO_2_ addition/remo val, ±hAQP5, with increasing amounts of injected hCA II—are the same as in Figure 3A for the ΔpH_S_ data.

*Conclusions.* Both for CO_2_/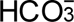 addition and removal: (a) In the absence of hCA II, expression of hAQP5 (gray vs. black bars) increases the magnitude of (dpH_S_/dt)_Max_, and the difference in mean values is statistically significant. The same is true for the expression of hAQP5 at each of the increasing hCA II levels (compare light-vs. dark-blue, red, and green bars). (b) In the absence of hAQP5, injecting hCA II in ever-greater amounts (lighter bars) causes (dpH_S_/dt)_Max_ to trend upward, but is without a statistica l ly significant effect on (dpH_S_/dt)_Max_. And (c) in the presence of hAQP5, injecting increasing amounts of hCA II tends to produce ever-greater magnitudes of (dpH_S_/dt)_Max_, with several of the differences reaching statistical significance, especially for CO_2_/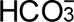 addition. And (d) the effects on (dpH_S_/dt)_Max_ are in the same direction but statistically more robust than for ΔpH_S_ in Figure 3*A,B*.

*Interpretation: (dpH_S_/dt)_Max_ in Figure 5A,B; ±hAQP5, ΔhCA II, –bCA.* Our analysis of these data is similar to those for ΔpH_S_ in <u>sections above</u>.^7^

### ±hAQP5, ΔhCA II, –bCA: Effects on other oocyte parameters

**Initial pH_i_** Figure 6A summarizes the initial pH_i_ values of oocytes subjected to the eight protocols of Figure 2. All of the mean values are within 0.1 pH units of each other. Statistical analyses reveal no significant differences.

**ΔpH_i_** Figure 6B summarizes the ΔpH_i_ values—that is, the decreases in pH_i_—elicited by applicatio n of 1.5% CO_2_/NaHCO_3_ for the eight protocols. These values are all well within 0.1 of each other, and statistical analyses shows no statistically significant differences among the eight ΔpH_i_ mean values.

**β_I_** Figure 6C summarizes the intrinsic buffering power values of the eight groups of oocytes. Statistical analyses show no significant differences among the groups. The modest downward trend from the leftmost bars to the rightmost bars may reflect a faster equilibration of CO_2_ within oocytes with higher injected levels of hCA II, especially with hAQP5 expression. With a slower CO_2_ equilibration (see Figure 2A), the measured ΔpH_i_ would tend to underestimate the value that we would have obtained at t = ∞, and thus lead to an artificially elevated computed β_I_ value.

***P*_f_** After monitoring pH_S_ and pH_i_, we performed the *P*_f_ assay with each oocyte. Figure 6D summar izes our *P*_f_ data. The injection of cRNA encoding hAQP5 significantly increases *P*_f_ for each of the four groups of oocytes previously injected with no or increasing amounts of hCA II (compare lighter vs. darker bars in blue, red, and green). Thus, we can conclude that the oocytes express hAQP5 that traffics normally to the plasma membrane. Because the increase in *P*_f_ induced by hAQP5 was virtually the same for the four doses of hCA II, it is most likely that the CA has no effect on the monomeric pores that are responsible for the osmotic water permeability of hAQP5.

### ±hAQP5, –hCA II, +bCA: Effects on ΔpHS & (dpHi/dt)Max

In the final four sections of Results, we examine oocytes studied in the presence of extracellular bCA and thereby explore the functional interaction between hAQP5 and CA_o_ in the transmembrane diffus io n of CO_2_.

In this first of the final four sections, we study oocytes in the absence of injected hCA II. On Day 1, we injected oocytes with either “H_2_O” or “H_2_O + cRNA” encoding “hAQP5” and then, on Day 4, we injected all oocytes with “Tris”. During the experiment, we augment the extracellular solutions with 0.1 mg/ml of bCA, the same level as used previously by Musa-Aziz *et al*. (2014*b*). This protocol is identical to that of Figure 2AB (–bCA) except for the presence of bCA.

**pH_S_** The red record in Figure 7A shows the pH_S_ data for a representative “H_2_O + Tris” oocyte and, in Figure 7B, the comparable data for an “hAQP5 + Tris” oocyte. Notice that the pH_S_ transients in Figure 7A (+bCA, extracellular) are many fold larger—both with CO_2_/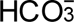 addition and removal—than their counterparts in Figure 2A (–bCA). Musa-Aziz et al. (2014b) had previously made a similar observation with extracellular hCA II (compare their figs. 13g and 15g).

**Figure 7.**
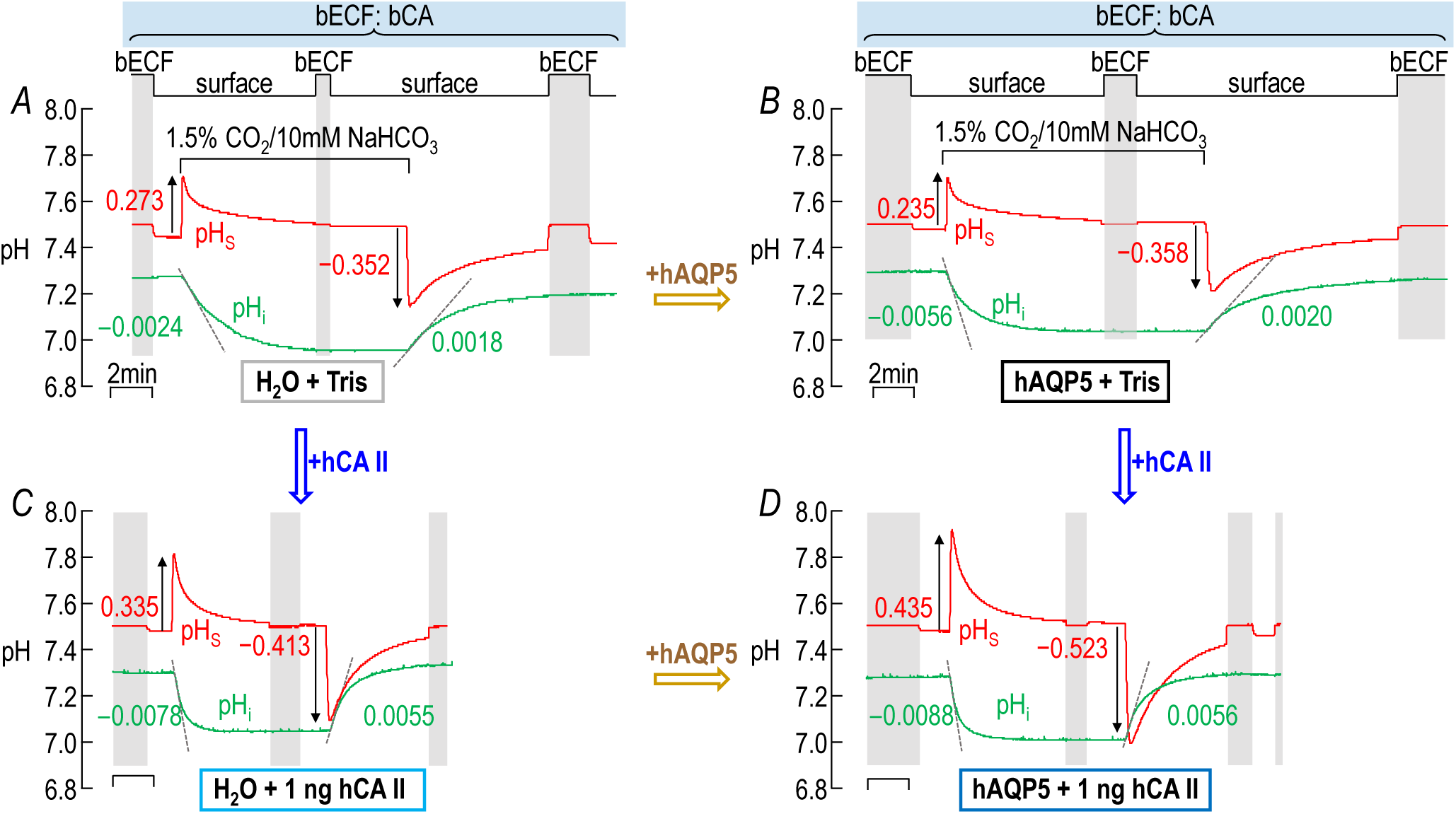
Representative pH_S_ and pH_i_ recordings: ±hAQP5, ±cytosolic hCA II, all in the presence of extracellular bCA (i.e., +bCA). *A*, “H_2_O + Tris”. We injected this oocyte with H_2_O (as a control for cRNA encoding hAQP5) on Day 1, and then with “Tris” (as a control for bCA II) on Day 4. *B*, “hAQP5 + Tris”. This oocyte, we injected on Day 1 with cRNA encoding hAQP5, and then on Day 4 with “Tris”. *C*, H_2_O + 1 ng hCA II”. Similar to panel *A* except that on Day 4, we injected the oocyte with hCA II enzyme dissolved in “Tris”. *D*, “hAQP5 + 1 ng hCA II”. Similar to panel *B* except that on Day 4, we injected the oocyte with hCA II enzyme dissolved in “Tris”. The numbers in red are ΔpH_s_ and in green are (dpH_i_/dt)_Max_ for the oocytes presented here. The stair-step line at the top of *A* and *B* indicates the position of the pH_S_ electrode. The vertical shaded bars indicate times during w_hi_ch the pH_S_ electrode is in the bulk extracellular fluid (bECF) for recalibration; _at_ other times, the pH_S_ electrode dimpl_es_ the cell surface for actual pH_S_ measurements. The colors of the rectangles that surround the panel labels (e.g., “H_2_O + Tris”) correspond to the colors of the bars in Figure 8 and later figures. ECF: bCA indicates that we obtained all measurements while exposing oocytes to solutions containing 0.1 mg/ml of bCA.

*Theoretical considerations.* Note that in comparing pH_S_ transients in Figure 7A vs. Figure 2A, the pH_S_ electrode is “cis” to the added bCA in Figure 7. A major reason that bCA increases the magnitude of ΔpH_S_ is that—near the extracellular surface of the plasma membrane—the added bCA greatly accelerates H^+^ consumption during CO_2_ influx (see Figure 1A), and H^+^ production during CO_2_ efflux (see Figure 1B). Although the pH_S_ electrode directly senses changes in [H^+^]_S_, these only indirectly reflect CO_2_ fluxes. Thus, by comparing ±bCA, we can reach no intuitive conclusions about the possible effect of bCA on transmembrane CO_2_ fluxes from pH_S_ data. However, previous mathematical simulations predict that the CO_2_ fluxes must have increased under these conditions (Musa-Aziz *et al*., 2014*b*; Occhipinti *et al*., 2014), even though in some cases the effects may be too small to measure.

*Exemplar data* in Figure 7A *vs. B.* The pH_S_ records show that—in the presence of extracellular bCA— the expression of hAQP5 does not produce an apparent increase in the magnitudes of the pH_S_ transients. This apparent lack of effect is quite different from the large fractional increases in ΔpH_S_ magnitudes that we observed in the absence of extracellular bCA (i.e., Figure 2B vs. Figure 2A), where the ΔpH_S_ magnitudes are so much smaller.

*Summary.* The gray and black bars in Figure 8A show the mean ΔpH_S_ data for CO_2_/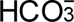 addition, ±hAQP5, all in the absence of hCA II but presence of extracellular bCA. Figure 8B shows corresponding data for CO_2_/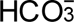 removal.

**Figure 8.**
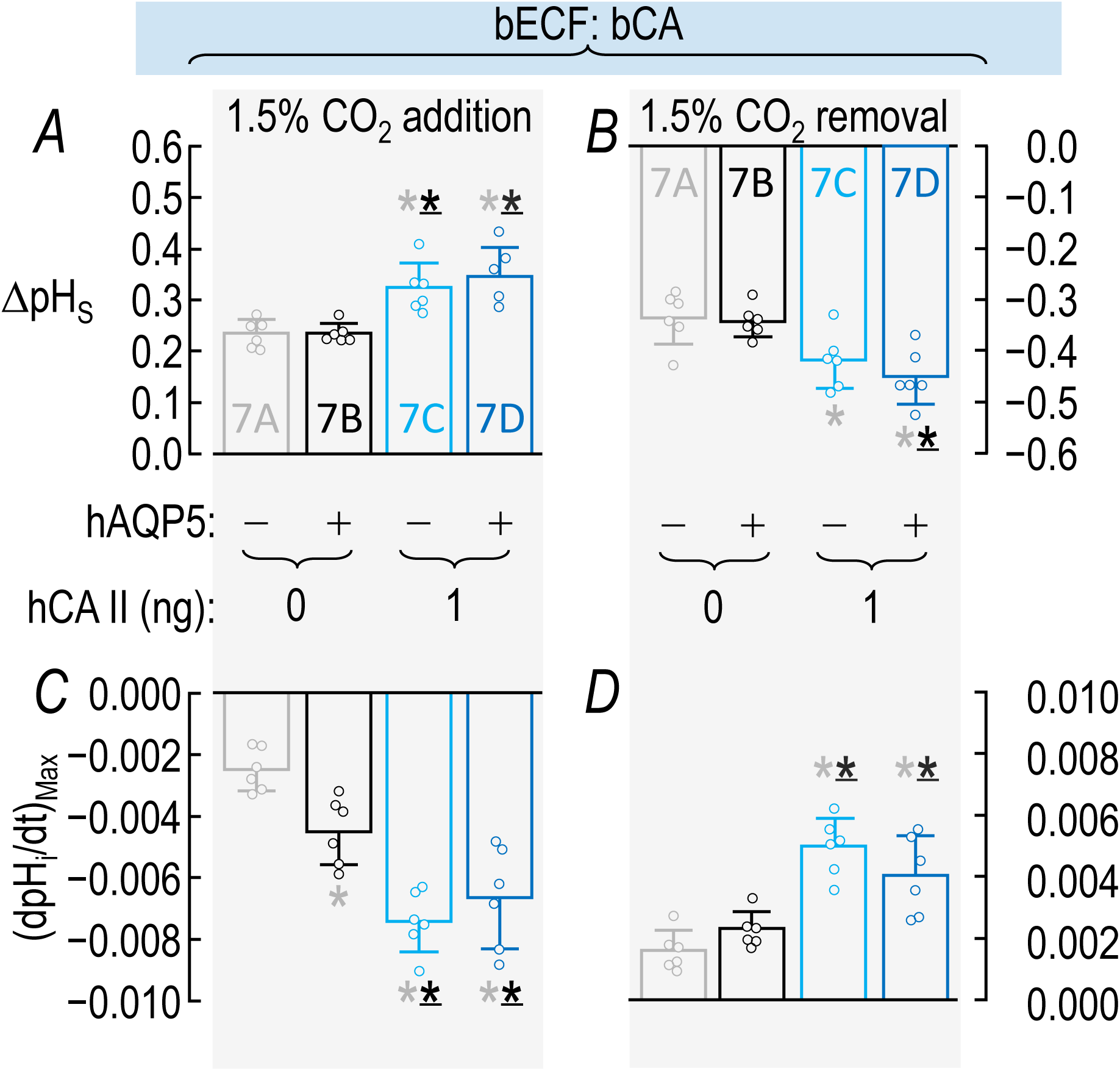
Summary of (pH_S_ and (dpH_i_/dt)_Max_ data from experiments like those in Figure 7: ±hAQP5, ± cytosolic hCA II, all in the presence of extracellular bCA (i.e., +bCA). This figure is analogous to Figure 3, which sum_ma_rized ΔpH_S_ and (dpH_i_/dt)_Max_ data obtained from oocytes in the absence of bCA. *A*, Summary of ΔpH_S_ upon addition of 1.5% CO_2_/10 mM 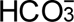. We computed individual ΔpH_S_ values as outlined in “Methods” > “Electrophysiological measurements” > “Calculation of ΔpH_S_”. *B*, Summary of ΔpH_S_ upon removal of 1.5% CO_2_/10 mM 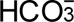. *C*, Summary of (dpH_i_/dt)_Max_ upon addition of 1.5% CO_2_/10 mM 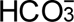. We computed individual ΔpH_S_ values as outlined in “Methods” > “Electrophysiological measurements” > “Calculation of “maximal” dpH_S_ /dt”. *D*, Summary of (dpH_i_/dt)_Max_ upon removal of 1.5% CO_2_/10 mM 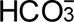. Data are presented as mean ± SD. “−” and lighter-colored bars indicate that we injected oocytes injected with H_2_O; “+” and darker-colored bars, with cRNA encoding hAQP5 on Day 1. “0” and “1”, indicate the amount of hCA II (ng) injected into oocytes on Day 4 Gray/black and light-/dark-blue pairs of bars indicate 4 groups of oocytes. These colors correspond to the colors of the rectangles that surround panel labels (e.g., “H_2_O + Tris”) in Figure 7. A star associated with a bar indicates a statistically significant difference when comparing the bar to another bar with the same color as the star. The star/bar combinations are meant to be read right to left; we indicate statistical significance between two bars only once, associating the star with the bar further to the right. Thus, the mean value represented by the dark-blue bar in panel *A* differs significantly from the means of the light-gray and black (underscored to indicate darker than gray) bars. Conversely, the gray bar, which has no stars, differs significantly from the means of the two bars with a gray star. The absence of a star indicates a lack of statistical significance. See Statistics Table 8 for *P*-values.

*Conclusions.* Considering oocytes examined in the presence of bCA: In the absence of hCA II, expression of hAQP5 (black vs. gray bars) does not significantly affect ΔpH_S_, either for CO_2_/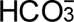 addition or removal.

*Interpretation: ΔpH_S_ (gray vs. black bars) in Figure 8A,B: ±hAQP5, –hCA II, +bCA.* (1) bCA promotes large transmembrane CO_2_ gradients. During CO_2_/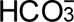 addition, bCA maintains relatively high CO_2_ levels near the extracellular face of the membrane and markedly increases CO_2_ influxes. During the subsequent CO_2_/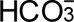 removal, bCA maintains relatively low CO_2_ levels at the cell surface. Thus, even in the absence of hAQP5 (gray bars), the CO_2_ fluxes enabled by bCA are very high. (2) In parallel, in a “cis-side” effect, bCA greatly increases the magnitudes of ΔpH_S_, as we can see by comparing Figure 8A*,B* vs. Figure 3A*,B*. Thus, during CO_2_/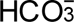 addition, bCA enhances H^+^ consumption, whereas during CO_2_/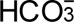 removal, bCA enhances H^+^ production. (3) With expression of hAQP5 (black bars)— combining an increased *P*_M,CO2_ with the capacity of bCA to replenish/consume cell-surface CO_2_— cytosolic CA activity becomes rate limiting. Thus, during CO_2_ addition, CO_2_ rapidly builds up near the inner surface of the membrane, limiting the transmembrane gradient, and choking CO_2_ influx—especia l ly as assessed by a pH_S_ electrode that is “cis” to the bCA. The same is true in reverse during CO_2_/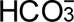 removal. (4) The evidence that hAQP5 does indeed increase *P*_M,CO2_ will come below in our analysis of Figure 8C.

**dpH_i_/dt** The green records in Figure 7A and B show the pH_i_ data collected simultaneously with the red pH_S_ transients (presented above) in these same panels. Note that (dpH_i_/dt)_Max_ values in Figure 7A (+bCA) are substantially greater—both with CO_2_/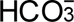 addition and removal—than their counterparts in Figure 2A (–bCA). Musa-Aziz et al. (2014b) had previously made a similar observation with extracellular hCA II (compare their figures 13e and 15e). Note that in comparing pH_i_ changes in Figure 7A vs. Figure 2A, the pH_i_ electrode is “trans” to the bCA that we add to the bECF. During CO_2_/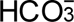 addition, bCA accelerates extracellular CO_2_ formation and thus maintains a relatively high [CO_2_] near the extracellular surface of the membrane. Similarly, during CO_2_/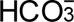 removal, bCA accelerates CO_2_ consumption and thus maintains a relatively low [CO_2_] at the extracellular membrane surface. The greater transmembrane CO_2_ gradients lead to greater CO_2_ fluxes and thus faster “trans-side” pH_i_ changes. Thus, comparing Figure 7A vs. Figure 2A, we can conclude that bCA increases transmembrane CO_2_ fluxes.

If we now compare the pH_i_ records in Figure 7A and Figure 7B, we see that hAQP5 expression produces a substantial acceleration in the rate of pH_i_ decrease during CO_2_/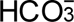 addition, consistent with an hAQP5-dependent increase in CO_2_ influx. During CO_2_/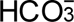 removal, the presence of hAQP5 does not substantially speed the pH_i_ increase.

*Summary.* The gray and black bars in Figure 8C summarize the downward (dpH_i_/dt)_Max_ for CO_2_/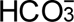 addition, ±hAQP5, all in the presence of extracellular bCA but absence of hCA II. Figure 8D summarizes the corresponding data for CO_2_/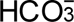 removal.

*Conclusions.* Considering oocytes examined in the absence of hCA II but presence of bCA (Figure 8C*,D*): During CO_2_/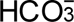 addition, expression of hAQP5 (black vs. gray bars) significantly increases the magnitude of (dpH_i_/dt)_Max_. During CO_2_/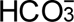 removal, expression of hAQP5 causes (dpH_i_/dt)_Max_ to trend faster, although the difference is not statistically significant.

*Interpretation: (dpH_i_/dt)_Max_ (gray vs. black bars) in Figure 8C,D; ±hAQP5, –hCA II, +bCA.* See our ΔpH_S_ “Interpretation” <u>immediately above</u>.^8^ (1) bCA promotes large transmembrane CO_2_ gradients. (2) In a “trans-side” effect, bCA greatly increases the magnitudes of (dpH_i_/dt)_Max_, as we can see by comparing gray and black bars in Figure 8C*,D* (+bCA) vs. Figure 3C*,D* (–bCA)—consequences of the gradient effect in point #1. (3) Here in Figure 8C (+bCA), hAQP5 increases the magnitude of (dpH_i_/dt)_Max_ during CO_2_/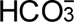 addition, demonstrating the hAQP5 increases *P*_M,CO2_. In Figure 3C (–bCA), the effect of expressing hAQP5 was nil because the limited availability of extracellular CO_2_ choked influx.

### ±hAQP5, +hCA II, +bCA: Effects on ΔpHS & (dpHi/dt)Max

To continue our exploration of the functional interaction of hAQP5 with CAs in the diffusion of CO_2_, in this next series of experiments, we not only augment the extracellular solutions with 0.1 mg/ml bCA, but also inject hCA II. On Day 1, we injected all oocytes with either “H_2_O” or “H_2_O + cRNA” encoding “hAQP5” and then, on Day 4, we injected all oocytes with 1 ng hCA II.

**pH_S_** The red records in Figure 7C (–hAQP5, +hCA II, +bCA) and Figure 7D (+hAQP5, +hCA II, +bCA) are analogous to those in panels *A* and *B*, except for the injection of hCA II. We now make three sets of comparisons:

a. Comparing the red pH_S_ records between Figure 7C (+hCA II) vs. Figure 7A (–hCA II)—having in common –hAQP5, +bCA—we see that addition of 1 ng hCA II increases the magnitudes of the pH_S_ transients—both with CO_2_/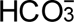 addition and removal. Recall that Musa-Aziz et al. (2014a) made similar ΔpH_S_ observations ±cytosolic hCA II, though in the absence of added CA_o_. Because the pH_S_ electrode is “trans” to the added hCA II (in comparing Figure 7C vs. Figure 7A), we can conclude that the hCA II has increased the transmembrane CO_2_ flux in the continued presence of extracellular bCA.
b. Comparing the pH_S_ records in Figure 7D (+hCA II) vs. Figure 7B (–hCA II)—having in common +hAQP5, +bCA—we see that addition of 1 ng hCA II again increases the magnitudes of the pH_S_ transients.
c. Comparing Figure 7D (+hAQP5) vs. Figure 7C (–hAQP5), we see that—on a background of injected hCA II and extracellular bCA—hAQP5 expression has limited effect.
d. Note that the pH_S_ transients here in Figure 7C*,D* (±hAQP5, +hCA II, +bCA) are much larger than their counterparts in Figure 2C*,D* (±hAQP5, +hCA II, –bCA).

*Summary.* (a) The gray vs. light-blue bars in Figure 8A show the mean ΔpH_S_ data for CO_2_/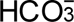 addition, ± hCA II, all in the absence of hAQP5 but presence of bCA. Figure 8B shows the corresponding data for CO_2_/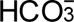 removal. (b) The black vs. dark-blue bars compare ±hCA II, but in the presence of both hAQP5 and bCA. (c) The light-blue vs. dark-blue bars compare ±hAQP5, all in the presence of both hCA II and bCA.

*Conclusions.* (a) The difference in mean values represented by the gray and light-blue bars (±hCA II in the absence of hAQP5) is statistically significant, as is (b) the difference between the black and dark-blue bars (±hCA II in the presence of hAQP5). Note that the effects in ‘a’ and ‘b’ correspond to the previously described synergistic effects of CA_i_ and CA_o_ (Musa-Aziz *et al*., 2014*a*, 2014*b*; Occhipinti *et al*., 2014). (c) The difference between the light-vs. dark-blue bars (±hAQP5 in the presence of hCA II and bCA) is not significant. (d) The statistical analysis summarized in Statistics Tables 8*E,F* compares ΔpH_S_ amplitudes summarized in Figure 8A*,B* (+bCA) vs. the gray/black/light-blue/dark-blue bars in Figure 3A*,B* (bCA). The result is that the effect of bCA is significant.

*Interpretation: ΔpH_S_ (colorful bars) in Figure 8A,B; ±hAQP5, +hCA II, +bCA.* See “Interpretation” for ΔpH_S_ in the <u>previous section</u>.^9^ Namely: (1) the bCA enhances transmembrane CO_2_. (2) The ΔpH_S_ data summarized by the light/dark-blue bars in Figure 8A*,B* (+bCA) are in stark contrast to the analogous data in Figure 3A*,B* (–bCA)—note that this is a “cis” comparison (i.e., ±bCA→pH_S_)—where the ΔpH_S_ values are only about ⅛ to ½ as large during CO_2_/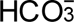 addition. (3) Expression of hAQP5 in the presence of 1 ng injected hCA II (light-vs. dark-blue bars), even with the increase in *P*_M,CO2_, does not increase ΔpH_S_ magnitudes because CO_2_ fluxes are choked by insufficient cytosolic CA activity (i.e., insufficient transmembrane CO_2_ gradient). And (4) recall that hAQP5 does indeed increase *P*_M,O2_ (see Figure 8C, below). In addition to points #1 – #4, which are analogous to the corresponding points made in the previous section: (5) in a “trans-side” effect, injection of 1 ng of hCA II—either in the absence of hAQP5 (gray vs. light-blue bars) or the presence of hAQP5 (black vs. dark-blue bars)—increases the ΔpH_S_ magnit udes because the hCA II is able to increase transmembrane CO_2_ gradients sufficiently under these conditions.

**dpH_i_/dt** The green records in Figure 7C and D show the pH_i_ data that correspond to the pH_S_ data presented above, and lead to three comparisons:

a. Comparing the pH_i_ records between Figure 7C (+hCA II) vs. Figure 7A (–hCA II)—both in the absence of hAQP5—we see that addition of 1 ng hCA II substantially increases the rates of pH_i_ changes, both with CO_2_/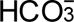 addition and removal. Musa-Aziz et al. (2014a) made similar (dpH_i_/dt)_Max_ observations ±hCA II, though in the absence of added CA_o_.
b. Comparing the pH_i_ records in Figure 7D(+hCA II) vs. Figure 7B (–hCA II)—now both in the presence of hAQP5—we again see that addition of 1 ng hCA II again increases the magnitudes of (dpH_i_/dt)_Max_.
c. Comparing the pH_i_ records in Figure 7D (+hAQP5) vs. Figure 7C (–hAQP5), we see that—on a background of injected hCA II and extracellular bCA—hAQP5 expression has little effect.
d. Finally, the magnitudes of (dpH_i_/dt)_Max_ here Figure 7C*,D* are modestly larger than their Figure 2C*,D* counterparts.

*Summary.* The layout and meaning of the bars in Figure 8C*,D* are the same as in Figure 8A*,B*.

*Conclusions.* As we saw for the ΔpH_S_ data, the differences between (a) the two lighter-colored bars (– hAQP5, ±hCA II,) as well as (b) the two darker-colored bars (+hAQP5, ±hCA II) are statistica l ly significant. (c) The difference between the light- and dark-blue bars (±hAQP5, + hCA II, +bCA) is not.

The statistical analysis summarized in Statistics Tables 8*G,H* compares (dpH_i_/dt)_Max_ magnitudes from Figure 8C*,D* (+bCA) vs. the gray/black/light-blue/dark-blue bars in Figure 3C*,D* (–bCA). The result is that the effect of bCA is significant.

*Interpretation: (dpH_i_/dt)_Max_ (colorful bars) in Figure 8C,D; ±hAQP5, +hCA II, +bCA.* See “Interpretation” for ΔpH_S_ <u>immediately above</u>. ^10^ Namely: (1) bCA magnifies transmembrane CO_2_ gradients. (2) In a “trans” effect, the magnitudes of the light- and dark-blue bars in Figure 8C*,D* (+hCA II, +bCA) are modestly larger than their counterparts in Figure 3C*,D* (+hCA II, –bCA)—a specific example of enhanced transmembrane CO_2_ gradients noted in point #1. (3) The light- and dark-blue bars (i.e., ±hAQP5, +hCA II, +bCA) are not significantly different, presumably because, 1 ng injected hCA II does not raise cytosolic CA activity sufficiently to prevent choking the dominant effects of hAQP5 (↑*P*_M,O2_) and bCA (↑CO_2_ gradient). We predict that, greater CA_i_ activities (e.g., 100 ng) would have alleviated the choke and revealed a much taller +hAQP5 bar. And (4) hAQP5 does indeed increase *P*_M,O2_ as evidenced by the comparison of gray vs. black bars in Figure 8C, above. In addition to points #1 – #4 (analogous to those made in previous section), (5) we note that—because of a “cis” effect—we cannot intuitively interpret the effects of injecting hCA II on (dpH_i_/dt)_Max_, either in the absence of hAQP5 (gray vs. light-blue bars) or the presence of hAQP5 (black vs. dark-blue bars). Even though we cannot intuitively assess ±hCA II effects from (dpH_i_/dt)_Max_, we know from the corresponding ΔpH_S_ data in Figure 8A*,B* that the injection of hCA II—by enhancing transmembrane CO_2_ gradients—must have increased the CO_2_ fluxes.

### ±hAQP5, ±hCA II, +bCA: Effects on pHS relaxation

*Summary.* The layout of Figure 9A*,B* is similar to that of Figure 8A*,B* (i.e., ΔpH_S_), except that here in Figure 9A*,B* we examine (dpH_S_/dt)_Max_ as we did in Figure 5A*,B*.

**Figure 9.**
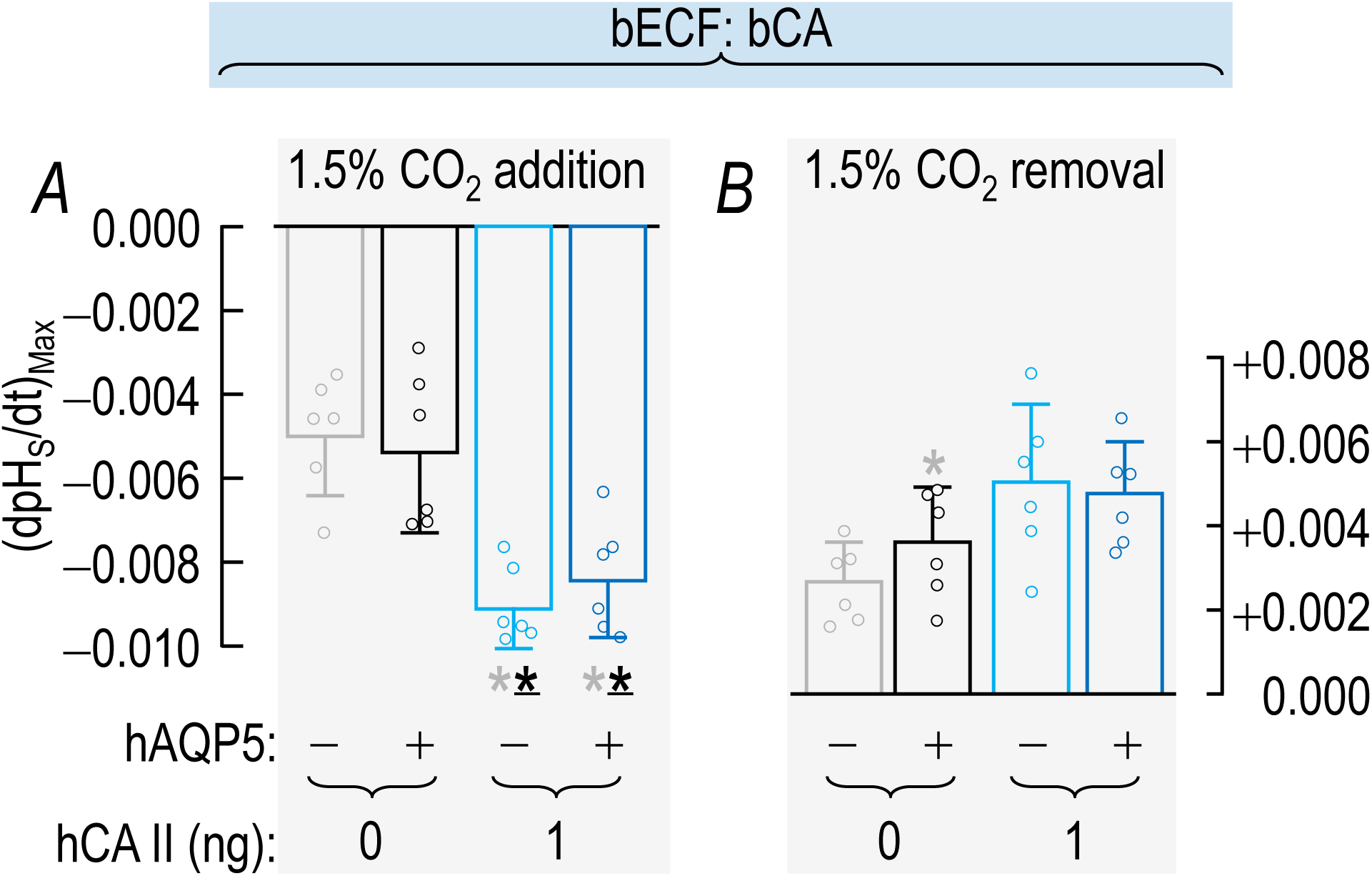
Summary of (dpH_S_/dt)_Max_ data from experiments like those in Figure 8: ±hAQP5, ±cytosolic hCA II, all in the presence of extracellular bCA (i.e., +bCA). This figure is analogous to Figure 5, which summarized (dpH_S_/dt)_Max_ data obtained from oocytes in the absence of bCA. *A*, Summary of (dpH_S_/dt)_Max_ upon addition of 1.5% CO_2_/10 mM 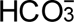. We computed individual (dpH_S_/dt)_Max_ values as described in Figure 4. *B*, Summary of (dpH_S_/dt)_Max_ upon removal of 1.5% CO_2_/10 mM 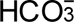. Data are presented as mean ± SD. “−” and lighter-colored bars indicate that we injected oocytes with H_2_O; “+” and darker-colored bars, with cRNA encoding hAQP5 on Day 1. “0” and “1” indicate the amount of hCA II (ng) injected into oocytes on Day 4. Gray/black and light-/dark-blue pairs of bars indicate 4 groups of oocytes. These colors correspond to the colors of the rectangles that surround panel labels (e.g., “H_2_O + Tris”) in Figure 7. A star associated with a bar indicates a statistically significant difference when comparing the bar to another bar with the same color as the star. The star/bar combinations are meant to be read right to left; we indicate statistical significance between two bars only once, associating the star with the bar further to the right. Thus, the mean value represented by the dark blue bar in panel *A* differs significantly from the means of the gray and black (underscored to indicate darker than gray) bars. Conversely, the gray bar, which has no stars, differs significantly from the means of the two bars with a gray star. The absence of a star indicates a lack of statistical significance. See Statistics Table 9 for *P*-values.

*Conclusions.* For oocytes examined in the presence of bCA:

(a) In the absence of hAQP5, injecting hCA II (gray vs. light-blue bars in Figure 9A) produces an increase in (dpH_S_/dt)_Max_ during CO_2_/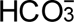 addition; the difference is statistically significant. During CO_2_/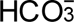 removal, the results trend toward a greater magnitude but do not reach statistical significa nce (Statistics Table 9*B*). These results in Figure 9A (–hAQP5, ±hCA II, +bCA) are in contrast to those in Figure 5A (–hAQP5, ±hCA II, –bCA), where injected hCA II is without statistically significant effect, and even the trends were mild.

And (b) in oocytes expressing hAQP5, injecting hCA II (black vs. dark blue bars) increases the magnitude of (dpH_S_/dt)_Max_ during CO_2_/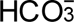 addition, with a statistically significant difference. This treatment produces a slight upward trend during CO_2_/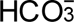 removal. Note that these increments (due to injecting hCA II) are no greater than for oocytes not expressing hAQP5. Viewed differently, expressing hAQP5 significantly increases the magnitude of (dpH_S_/dt)_Max_ in only 1 of 4 cases (CO_2_/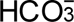 removal in the absence of hCA II, gray vs. black bars in Figure 9B). These results contrast to those in Figure 5 (– bCA) where—in the presence of hAQP5—increasingly large hCA II injections tended to produce graded increases (dpH_S_/dt)_Max_, and for both CO_2_/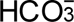 addition and removal. We note, however, that the hCA II injections in Figure 9 were either 0 or 1 ng, whereas in the Figure 5 (–bCA) study, the hCA II we as high as 100 ng.

(c) Expression of hAQP5 in the absence of hCA II (gray vs. black bars in Figure 9A*,B*) or the presence of hCA II (light-vs. dark-blue bars) has no effect on (dpH_S_/dt)_Max_ except during CO_2_/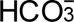 removal in the absence of hCA II (gray vs. black bars, Figure 9B). These results contrast with those of Figure 5 (– bCA), where hAQP5 expression increases (dpH_S_/dt)_Max_ magnitudes robustly and consistently.

(d) The (dpH_S_/dt)_Max_ magnitudes in Figure 9 are ∼2-to ∼3-fold greater than the analogous ones in Figure 5. The differences are greatest in the absence of hCA II, where the lighter-colored bars in Figure 9 are ∼10-fold greater.

*Interpretation: (dpH_S_/dt)_Max_ in Figure 9A,B; ±hAQP5, ±hCA II, +bCA.* Our analysis of these (dpH_S_/dt)_Max_ is similar to that for the ΔpH_S_ data <u>in the previous section</u>.^11^

### ±hAQP5, ±hCA II, +bCA: Effects on other oocyte parameters

The four bars in each of the panels of Figure 10 are analogous to the four leftmost bars of the four panels in Figure 6; the difference is that, here in Figure 10, we summarize experiments in which we exposed oocytes to 0.1 mg/ml extracellular bCA.

**Figure 10.**
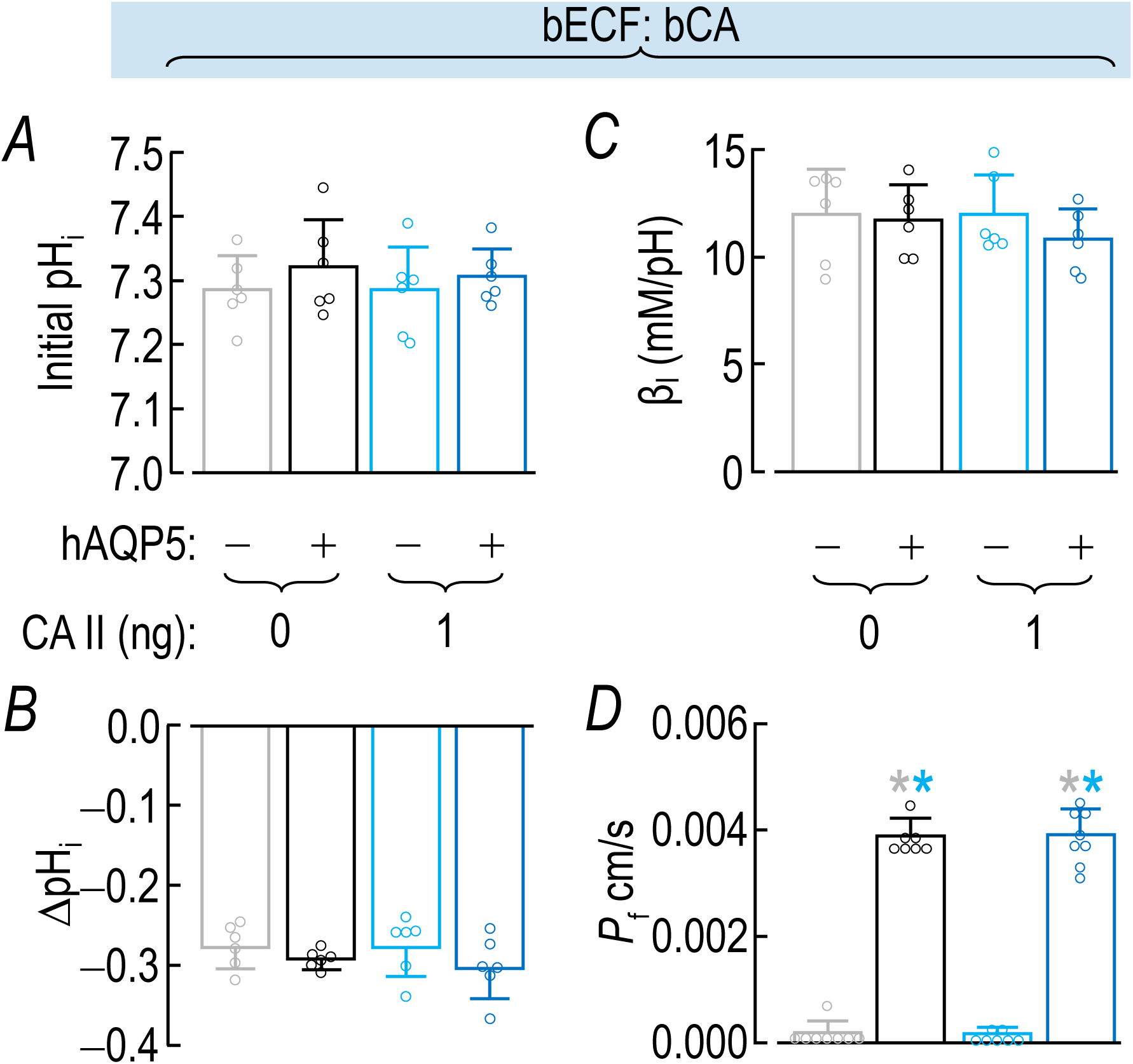
Summary of other oocyte parameters from experiments like those in Figure 7: ±hAQP5, ±cytosolic hCA II, all in the presence of extracellular bCA (i.e., +bCA). This figure is analogous to Figure 6, which summarized comparable data obtained from oocytes in the absence of bCA. *A*, Summary of initial pH_i_ values. We computed individual initial pH_i_, ΔpH_i_ (see panel *B*), and β_I_ (see panel *C*) values as outlined in “Methods” > “Electrophysiological measurements” > “Calculation of intrinsic intracellular buffering power”. *B*, Summary of ΔpH_i_ elicited by addition of 1.5% CO_2_/10 mM 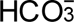. *C*, Summary of intrinsic buffering power (β_i_). *D*, Summary of *P*_f_. We computed individual initial pH_i_, ΔpH_i_ (see panel *B*), and β_I_ (see panel *C*) values as outlined in “Methods” > “Electrophysiological measurements” > “Measurement of *P*_f_”. Data are presented as mean ± SD. “−” and lighter-colored bars indicate that we injected oocytes injected with H_2_O; “+” and darker-colored bars, with cRNA encoding hAQP5 on Day 1. “0” and “1”, indicate the amount of hCA II (ng) injected into oocytes on Day 4. Gray/black and light-/dark-blue pairs of bars indicate 8 groups of oocytes. These colors correspond to the colors of the rectangles that surround panel labels (e.g., “H_2_O + Tris”) in Figure 7. A star associated with a bar indicates a statistically significant difference when comparing the bar to another bar with the same color as the star. A star associated with a bar indicates statistical significance when compared to the bar with the same color as the star. The star/bar combinations are meant to be read right to left; we indicate statistical significance between two bars only once, associating the star with the bar further to the right. Thus, the mean value represented by the dark blue bar in panel *D* differs significantly from the means of the gray and light-blue bars. Conversely, the gray bar, which has no stars, differs significantly from the means of the black and dark-blue bars. The absence of a star indicates a lack of statistical significance. See Statistics Table 10 for *P*-values.

**Initial pH_i_** Statistical analyses, as in the case of Figure 6A, reveal no significant differences among the four mean initial pH_i_ values in Figure 10*A*. Moreover, Statistics Table 10*E*, which summarizes an analysis of values in Figure 10*A* (+bCA) vs. the gray/black/light-blue/dark-blue bars in Figure 6A (–bCA), reveals no significant effect of adding bCA.

**ΔpH_i_** We observe no significant differences among mean ΔpH_i_ values in Figure 10*B*, as observed among groups in Figure 6B. However, Statistics Table 10*F*, which summarizes an analysis of values in Figure 10*B* (+bCA) vs. the gray/black/light-blue/dark-blue bars in Figure 6B (–bCA), reveals a small but significant difference due to the addition of bCA. The magnitudes of ΔpH_i_ values in Figure 6B are somewhat smaller, presumably reflecting the slower transmembrane equilibration of CO_2_ due to the absence of bCA, particularly at lower levels of injected hCA II.

**β _I_** Continuing the trend from the previous two panels, we observe no significant differences among mean β_I_ in Figure 10*C*, as described above among analogous conditions in Figure 6C. However, Statistics Table 10*G*, which summarizes an analysis of values in Figure 10*C* (+bCA) vs. the gray/black/light-blue/dark-blue bars in Figure 6C (–bCA), reveals a small but significant difference—similar to the analysis of the ΔpH_i_ data above. Because their ΔpH_i_ magnitudes tend to be smaller, the β_I_ values in Figure 6C (– bCA) are somewhat larger than those in Figure 10*C* (+bCA).

***P*_f_** As for the previous three panels, the mean values summarized in Figure 10*D* (+bCA)—now for *P*_f_—are indistinguishable from the comparable values in Figure 6D (–bCA). Our statistical analyses in ^S^tatistics Table 10*H* reveals no significant effect of bCA on *P*_f_. Thus, we can conclude that extracellular bCA—a mixture of CA I and CA II—does not interfere with the monomeric pores of hAQP5. Moreover, as we did in our analysis of Figure 6D, we can conclude, from a comparison of the black and dark-blue bars, that the hCA II does not functionally interfere with the monomeric pores.

## Discussion

### Historical context

Although previous papers on CO_2_ diffusion across membranes have addressed the role of channels or the role of CAs, the present paper is the first to undertake a systematic examination of both channels and CAs, as well as their synergistic interaction to enhance CO_2_ fluxes. The underlying principle is Fick’s law of diffusion, which we reproduce from Eqn (2):

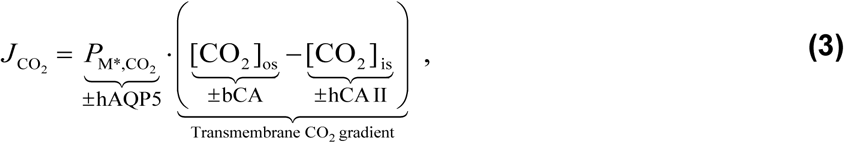

and embellish to emphasize that the key parameters in the present study increase *P*_M*,CO2_ (i.e., hAQP5) and the transmembrane CO_2_ gradient. During CO_2_ influx, bCA (when present) in the extracellular fluid maintains a relatively high [CO_2_]_os_, whereas hCA II (when present) maintains a relatively low [CO_2_]_is_, the result being an enhanced inwardly directed CO_2_ gradient. Conversely, during CO_2_ efflux, bCA in the extracellular fluid maintains a relatively low [CO_2_]_os_ whereas hCA II maintains a relatively high [CO_2_]_is_, the result being an enhanced outwardly directed CO_2_ gradient.

It is perhaps worth noting that similar principles are at work during O_2_ fluxes across the erythrocyte membrane. In that case, according to preliminary results (Moss *et al*., 2025; Occhipinti *et al*., 2025; Zhao *et al*., 2025), it is AQP1, the Rh complex, and as-yet-unidentified channel(s) that make the dominant contribution to *P*_M,O2_. Moreover, it is hemoglobin that maximizes transmembrane O_2_ gradients (ignoring hemoglobin diffusion within the cytoplasm) by serving as a sink or source of O_2_ near the plasma membrane. In the laboratory experiments on O_2_ efflux from RBCs, one can use an extracellular O_2_ scavenger like sodium dithionite to consume O_2_ near the extracellular face of the membrane, in an action that is analogous to that of bCA in the present experiments on CO_2_ efflux.

Working on artificial lipid bilayers, Gutknecht et al. (1977) showed that mobile CA enhances CO_2_ fluxes (measured using ^14^C-labeled CO_2_), but only in the presence of sufficient non-CO_2_/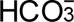 buffers (phosphate, HEPES, “Tris”). Their interpretation is that the CA increases the transmembrane CO_2_ gradient by replenishing CO_2_ on one side of the membrane and consuming it on the other. This replenishment (or consumption) requires that 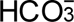 diffuse through the unconvected layers on opposite sides of the membrane to serve as either the source or product of the CA reactions. Moreover, the buffers act as either a source of or sink for H^+^ in these reactions.

The trio of papers by Musa-Aziz et al. (2014a, 2014b) and Occhipinti et al. (2014) extended this line of reasoning to *Xenopus* oocytes, which they injected with recombinant hCA II (to an apparent concentration ∼50% higher than in RBCs) and/or cRNA encoding hCA IV (which contributes to both extracellular-surface and cytosolic CA activity). In addition, they systematically varied [CO_2_]_o_ and [HEPES]_o_ and performed 3-dimensional reaction-diffusion mathematical modeling to enable a quantitative interpretation of the data. As anticipated from the earlier artificial lipid-bilayer work, they found that each CA alone augmented CO_2_ diffusion by increasing transmembrane CO_2_ gradients. However, the combination of the two is not simply additive but highly synergistic. The reason is that, with a CA on just one side of the membrane, net CO_2_ fluxes are limited by the build-up or depletion of CO_2_ on the other—the “choking” (or throttling) effect to which we allude in the Results section of the present paper. They also demonstrated an additional synergism involving an extracellular non-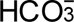 buffer, (i.e., HEPES), assessed by “trans-side” measurements of pH_i_, and examined the impact of increasing [CO_2_]_o_ during CO_2_/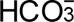 application. In their papers, they did not address the issue of CO_2_ permeability.

The first studies to address the *P*_M,CO2_ term in Eqn (3) were those by Nakhoul et al. (1998) and Cooper & Boron (1998), who showed that AQP1—in addition to being a H_2_O channel—is an effective CO_2_ channel. (Musa-Aziz et al. (2007) and Geyer et al. (2013) later examined a range of mammalian AQPs and showed that AQP5 has the highest CO_2_/H_2_O permeability ratio. The only study to broach channel-CA interactions was that by Nakhoul et al. (1998), who studied CO_2_-induced pH_i_ changes in oocytes with the vitelline membrane intact. They found that the stimulatory effect of AQP1 on CO_2_ influx into oocytes occurs only with bCA injected into the cytoplasm. In the present paper, the analogous experiment is summarized by the light-vs. dark-blue bars in Figure 3C, where hAQP5 expression causes (dpH_i_/dt)_Max_ to trend faster. Perhaps a difference, as suggested by others (Vilas *et al*., 2015), is that CA II binds to AQP1 but not AQP5.

The present paper builds on the work of Musa-Aziz and Occhipinti (Musa-Aziz *et al*., 2014*a*, 2014*b*; Occhipinti *et al*., 2014) and generally uses identical approaches, with three important differences:

First, rather than recombinant hCA II, in the present study we use hCA II purified commercially from RBCs.

Second, rather than heterologously expressing hCA IV, we add bCA to the extracellular fluid. The reasons for this switch are fourfold: (1) hCA IV expression in oocytes not only increases cell-surface CA activity, but also cytosolic CA activity. The oocyte membrane confines bCA to the outside of the cell. (2) The co-expression of hCA IV and hAQP5 adds an extra burden of heterologous expression and also introduces potential competition between the injected cRNAs encoding the two proteins (“ribosome steal”). (3) We also chose to add purified CA protein to the bECF because in principle we know the precise increase in CA_o_ activity, which is difficult to know in the case of protein expression. And (4) we chose bCA because it is a less-expensive combination of bCA I (less active) and bCA II (more active) vs. hCA II; this is an important practical consideration during continuous-flow experiments, which consume considerable volumes of experimental solutions.

Third, rather than working only with oocytes having a background *P*_M,CO2_, we alternated between injections of H_2_O and cRNA encoding hAQP5.

### pH measurements made “cis” vs. “inter” vs. “trans” to the altered parameter

The papers of (Musa-Aziz *et al*., 2014*a*, 2014*b*) and Occhipinti et al. (2014) introduced the concept of pH measurements, used as an indirect measure of something else (e.g., CO_2_ flux), being made “cis” or “trans” to a compartment with an alteration that directly affects pH (e.g., CA activity, non-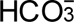 buffering power). Thus, if one wishes to assess the effects on CO_2_ flux of injecting (or not injecting) hCA II into the cytosol, the measurement of pH_i_—“cis” to the added hCA II—does not provide intuitive insight into the effects of hCA II” on transmembrane CO_2_ fluxes. Of course, at the same time as making the pH_i_ measurement, one may also be monitoring of pH_S_—“trans” to the added hCA II in the same cell. Interpreting the data from such a “trans” perspective does provide intuitive insight into CO_2_ fluxes. The same principles apply in the opposite sense if one wishes to assess the effects of adding (or not adding) bCA to the bECF. The measurement of pH_S_—“cis” to the added bCA—does not intuitive insight into the effects of bCA on CO_2_ fluxes, whereas the measurement of pH_i_—“trans”—does.

The reason for the “cis-side” prohibition is that the enzymatic activity of the CA produces or consumes H^+^, and thus can produce large pH changes that are independent of—and could conflated with the interpretation of—changes in transmembrane CO_2_ fluxes. Thus, intuitive interpretations can be extremely difficult, even though mathematical modeling can unravel alterations in “cis-side” pH from alterations in the flux of CO_2_ or other buffer components. Such unraveling was a major component of the studies of Musa-Aziz *et al*. (2014*a*, 2014*b*) and Occhipinti *et al*. (2014), which demonstrated that both cytosolic and extracellular CA increase transmembrane CO_2_ gradients and thus increase CO_2_ fluxes. Of course, avoiding a prohibited “cis-side” interpretation requires proper experimental design. However, the “cis-side” prohibition is more an issue of data interpretation: when altering any parameter that affects acid-base reaction rates in Figure 1A, an intuitive interpretation requires that one examine pH “trans” to the alteration.

Stated simply, “trans-side” pH measurements are valuable for intuitive interpretations because trans-side pH change can occur only as the result of altered CO_2_ fluxes across the membrane.

The experiments summarized by Figure 8 are a useful case study. bCA is present continuously in the bECF. Thus, when we inject hCA II, the pH_S_ measurements—“trans” to the hCA II—do provide intuitive insight into the effect of hCA II on transmembrane CO_2_ fluxes because, in this particular comparison, the bCA status is unchanging. Basing one’s intuition on pH_i_ measurements—“cis” to the added hCA II— would be unwise. However, if we compare Figure 8 (+bCA) with Figure 3 (–bCA), an intuitive assessment of bCA effects must come from pH_i_ measurements—“trans” to the ±bCA condition. Thus, “cis” vs. “trans” is not a matter of data per se, but a matter of perspective during data analysis.

Finally, in Figure 3 and Figure 8 we also assess the effects of expressing the integral membrane protein hAQP5. Both pH_S_ and pH_i_ measurements are neither “cis” nor “trans” to the altered expression of hAQP5 and thus the altered *P*_M,CO2_. One might use the term “inter” (Latin, “within”, “inside”, or “between”) to indicate the position of the protein relative to the membrane. In such an “inter” situation, one could use both pH_S_ and pH_i_ measurements to assess the data intuitively.

### Molecular mechanism of CO2 permeability

Given the suggestion that AQPs may conduct dissolved gases via the hydrophobic central pore of the tetramer (Wang *et al*., 2007; Boron, 2010), it is not surprising that molecular dynamics simulations are consistent with the hypothesis that O_2_ (Zhang & Chen, 2013) and CO_2_ (Alishahi & Kamali, 2019) can diffuse through the central pore of hAQP5. Preliminary data on hAQP5 suggest that (1) mutating amino-acid residues near the outer mouth of the central pore to residues with bulky side chains (e.g., T41F) or (2) creating a divalent-cation binding site (T41H) and then adding Ni^2+^ or Zn^2+^ greatly reduces ΔpH_S_, as supported by crystal structures and molecular dynamics (Shinn *et al*., 2024). In addition, a preliminary report suggests that, with hAQP1, the mercurial pCMBS can block one component of *P*_M,CO2_, the stilbene derivative DIDS can block an equally large component, and the two together than eliminate the CO_2_ permeability of hAQP1 (Musa-Aziz *et al*., 2025). Thus, the emerging picture is that some CO_2_ can permeate the four hydrophilic monomeric pores (at least of hAQP1) and that another component— presumably the major component of CO_2_—moves through the central pore of hAQP5.

### Diagnostic power of ΔΔΔpHS ±hCA II in identifying enhanced CO2 permeability

In Figure 11, we rearrange CO_2_-influx bars from Figure 3A so that we can easily compare the effects— on ΔpH_S_—of adding a CA, namely hCA II, on the side of the membrane “trans” to the pH_S_ measureme nt. The pair of bars on the left of Figure 11*A* shows that, in the absence of hAQP5, injecting 1 ng of hCA II into oocytes has only a minor effect on ΔpH_S_. That is, the baseline ΔΔpH_S_—or ΔΔpH_S,Base_—is small. However, in the presence of a CO_2_ channel hAQP5, ΔΔpH_S_—or ΔΔpH_S,hAQP5_—is substantially greater. The difference between the two ΔΔpH_S_ values—the ΔΔΔpH_S_ due to the presence of hAQP5 or ΔΔΔpH_S,hAQP5_—is statistically significant (Supplemental Table 11*A*).

**Figure 11.**
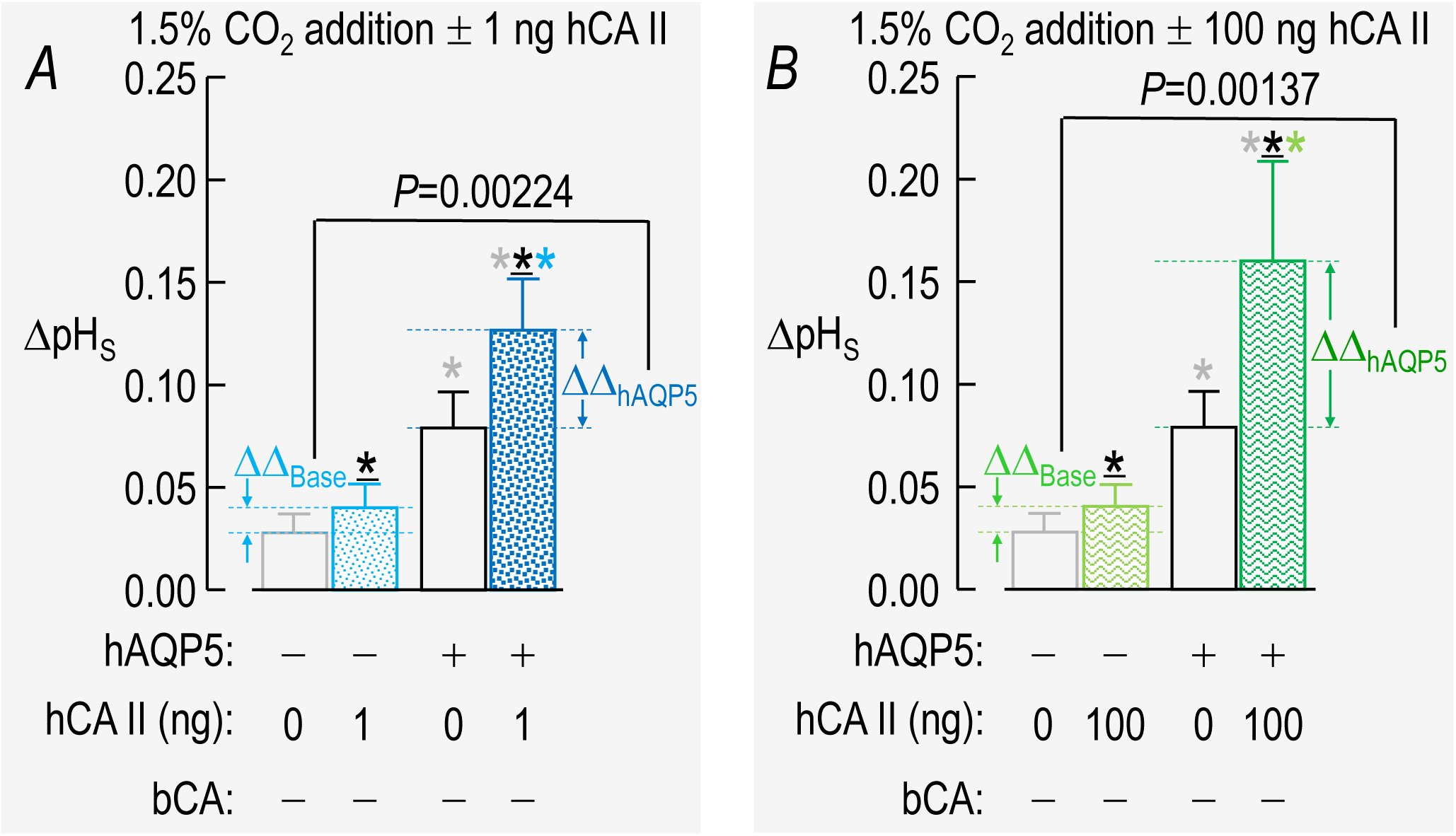
Effect of expressing hAQP5 on ((pH_S_: comparison of selected ΔpH_S_ bars extracted from Figure 3*A* (–bCA). Here we rearrange bars from Figure 3*A* to juxtapose two bars representing the same hAQP5 status (“–”, absent; “+”, heterologously expressed), the left bar in a pair representing the absence of hCA II, and the right bar representing the presence of injected hCA II. *A*, ΔΔpH_S_ determined with ±1 ng of injected hCA II, during addition of CO_2_/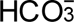. *B*, ΔΔpH_S_ determined with ±100 ng of injected hCA II, during addition of CO_2_/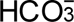. For a pair of bars, the difference between ΔpH_S_ values is ΔΔpH_S_. Thus, ΔΔ_Base_ is difference between ΔpH_S_ values under baseline conditions (i.e., no hAQP5), induced by the addition of the trans-side CA (i.e., CA II); ΔΔ_hAQP5_ is the corresponding difference between ΔpH_S_ values in AQP5-expressing oocytes. Comparing two pairs of bars, the difference between ΔΔ_hAQP5_ and ΔΔ_Base_ values is ΔΔΔpH_S,hAQP5_. Although no bCA is present in any of these experiments, we include bCA status (“–”, absent from bECF) to facilitate comparisons with Figure 12. The stars, which indicate statistical significance among the individual bars, have the same meanings as in Figure 3*A*. The *P*-values indicate statistical significance between the respective ΔΔ_Base_ and ΔΔ_hAQP5_ bar pairs, and reflect the statistical significance of the adding 1 ng hCA II (panel *A*) or 100 ng hCA II (Panel *B*), on the “trans” side of the membrane to the pH_S_ measurement. See Supplemental Table 11*A* and *B* for the statistics summary (including all *P*-values reported by the stars).

Figure 11B is a rearrangement of two pairs of CO_2_-influx bars, but with 100 ng of injected hCA II. The ΔΔpH_S,Base_ is similar to the value in Figure 11*A*. However, here the effect of hAQP5 expression, namely, ΔΔpH_S,hAQP5_ is even greater. Thus, it appears that the greater the level of hCA II, the greater is the effect of hAQP5 expression on ΔΔpH_S_. Although this set of comparisons is of CO_2_ influx, we could reach a similar set of conclusions for CO_2_ efflux (i.e., by rearranging bars in Figure 3B).

If we did not know the identity of the membrane protein expressed in these experiments, we could deduce that the protein must have a significant CO_2_ conductance, assuming that the protein itself lacks significant CA activity. We can imagine that the cytosolic CA, by serving as a sink for CO_2_ during influx, sucks CO_2_ into the cell. If *P*_M,CO2_ is rate limiting, the addition of a CO_2_ channel will lead to increased CO_2_ influx, a greater decrease in [CO_2_] at the extracellular cell surface, and thus a greater ΔpH_S_.

Note that in a cell in which it is impractical to inject or express a cytosolic CA, one might use a drug like acetazolamide to block endogenous CA II, and thereby perform an analogous ΔΔΔpH_S_ assay.

### Diagnostic power of ΔΔ(dpHi/dt)Max ±bCA in identifying enhanced CO2 permeability

In Figure 12, we perform the inverse analysis of Figure 11: We rearrange CO_2_-influx bars from Figure 3C and Figure 8C so that we can easily compare the effects—now on Δ(dpH_i_/dt)_Max_—of adding a CA— now bCA—on the side of the membrane “trans” to the pH_i_ measurement. The pair of bars on the left of Figure 12*A* shows that, in the absence of hAQP5, adding bCA to the bECF produces a modest increase in the magnitude of (dpH_i_/dt)_Max_—the Δ(dpH_i_/dt)_Max,Base_. On the other hand, in the presence of the CO_2_ channel hAQP5, the magnitude of (dpH_i_/dt)_Max_—Δ(dpH_i_/dt)_Max,hAQP5_—is substantially greater. The difference between the two Δ(dpH_i_/dt)_Max_ values—the ΔΔ(dpH_i_/dt)_Max_ due to the presence of hAQP5 or ΔΔ(dpH_i_/dt)_Max,hAQP5_—is statistically significant (Supplemental Table 12*A*).

**Figure 12.**
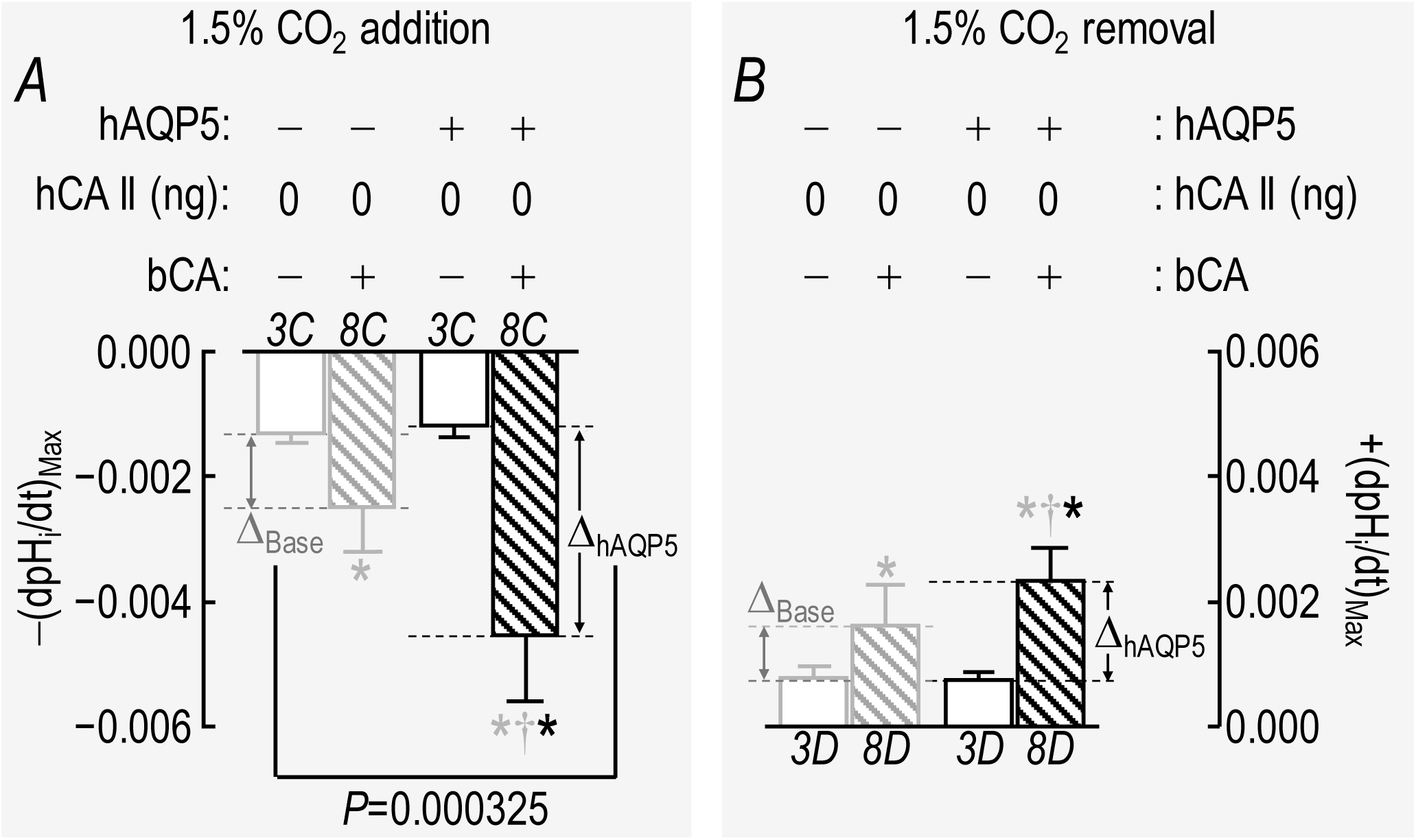
Effect of expressing hAQP5 on ((dpH_i_/dt)_Max_: comparison of selected (dpH_i_/dt)_Max_ bars extracted from Figure 3C vs. Figure 8C and from Figure 3D vs. Figure 8D. Here we rearrange the gray and black bars from Figure 3*C* (–bCA) and the gray and black bars from Figure 8*C* (+bCA) to juxtapose two bars representing the same hAQP5 status (“–”, absent; “+”, heterologously expressed), the left bar in a pair representing the absence of bCA, and the right bar in a pair representing the presence of bCA in the bECF. *A*, Summary of Δ(dpH_i_/dt)_Max_ ±bCA, upon addition of CO_2_/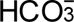. *B*, Summary of Δ(dpH_i_/dt)_Max_ ±bCA, upon removal of CO_2_/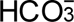. For a pair of bars, the difference between (dpH_i_/dt)_Max_ values is Δ(dpH_i_/dt)_Max_. Thus, Δ_Base_ is difference between (dpH_i_/dt)_Max_ values under baseline conditions (i.e., no hAQP5), induced by the addition of the trans-side CA (i.e., bCA); Δ_hAQP5_ is the corresponding difference between (dpH_i_/dt)_Max_ values in hAQP5-expressing oocytes. Comparing two pairs of bars, the difference between Δ_hAQP5_ and Δ_Base_ values is ΔΔ(dpH_i_/dt)_Max,hAQP5_. Although no hCA II is present in any of these experiments, we include hCA II status (“0 ng”, not injected) to facilitate comparisons with Figure 11. The stars, indicate statistical significance among individual bars; the gray star associated with the gray- and black-hatched bars indicates a significant difference compared to the open gray bar (–hAQP5, –bCA); the black star associated with the black-hatched bars (+AQP5, +bCA) indicates a significant difference compared to the open black bar (+hAQP5, –bCA); the gray dagger symbol associated with the black-hatched bars (+AQP5, +bCA) indicates a significant difference compared to the open gray-hatched bar (–hAQP5, +bCA). See Supplemental Table 12*A* and *B* for the statistics summary (including the all *P*-values reported by the stars).

Figure 12B shows a similar analysis for CO_2_/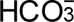 removal. Because the (dpH_i_/dt)_Max_ values are smaller, the Δ(dpH_i_/dt)_Max_ values also are of smaller magnitude than in Figure 12*B*. In this case, the difference between the two Δ(dpH_i_/dt)_Max_ values is not statistically significant. Thus, this assay may not be practical as shown. A future approach could be to raise [CO_2_]_o_, raise [HEPES]_o_, inject 100 ng of hCA II, and raise the bCA level (albeit at the increased cost of the enzyme in a continuously flowing solution). Some combination of these changes would increase the rates of pH_i_ change, and thereby make it easier to detect (dpH_i_/dt)_Max_ differences in both the influx and efflux assays.

### Advances and Limitations

**Synergisms** In the present study, we confirm the strong synergy between extracellular and cytosolic CAs (see Figure 3 and Figure 8). We also identify a strong synergy between a CO_2_ channel (i.e., hAQP5) and a cytosolic CA (i.e., hCA II; see pH_S_ data in Figure 3). Finally, we identify a strong synergy between a CO_2_ channel and an extracellular CA (i.e., bCA; see dpH_i_/dt data in Figure 8).

**“Cis-” vs. “trans-side” effects** Our group has previously pointed out the challenges of pH measurements made “cis” to a CA manipulation (Lu *et al*., 2006). For example, “cis-side” pH_i_ measurements led to the erroneous conclusion that cytosolic CA II binds to and thereby stimulates the Cl-HCO_3_ exchanger AE1, when in fact the added CA II was catalyzing the reaction 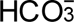 + H^+^ → CO_2_ + H_2_O, and producing a rapid pH_i_ increase because of catalysis, not transport.

The aforementioned trio of papers (Musa-Aziz *et al*., 2014*a*, 2014*b*; Occhipinti *et al*., 2014) systematically deals with the issue of pH measurements made “cis” to a manipulations that impact acid-base chemistry (e.g., modulation of CA activity, alterations in non-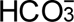 buffering power) and the necessity to focus on “trans” effects. The present paper builds upon and expands these concepts. Note that our modulation of *P*_M,CO2_—an “inter” situation—is neither “cis” nor “trans” to our electrodes and is thus immune from these considerations.

**“Inter” effect** We introduce this term to describe a maneuver that impacts the membrane separating two solutions (e.g. intra-vs. extracellular). For example, the act of introducing, deleting, mutating, or otherwise changing the activity of an integral membrane protein is “in between” “cis” and “trans”. Likewise, altering lipid composition or otherwise altering the chemistry of membrane lipids or their interaction with proteins or other substances would be an “inter” effect.

**pH_S_ relaxation assessed as (dpH_S_/dt)_Max_** Although we have previously assessed pH_S_ relaxation as the time constant of a single-exponential decay, we now introduce a more general tool for describing the initial rate of pH_S_ decay, especially one that that deviates substantially from an SExp time course. Although we implement this tool using a double-exponential curve fit, in principle we could use any function that, near t_Local_ = 0, fits pH_S_ vs. time with a monotonic decay.

**Novel diagnostic paradigms** Our approaches for assessing ΔΔΔpH_S_ and ΔΔ(dpH_i_/dt)_Max_ provide potentially valuable for assessing the CO_2_ permeability of candidate membrane proteins.

**Limitations** A drawback of using bCA rather than hCA IV is that, with bCA added to the bECF, the CA activity in the thin unconvected layer at the oocyte surface is probably less than in a case in which we express hCA IV at the cell surface.

In the present study, recognizing an already large matrix of experimental conditions, we did not explore bCA levels higher than 0.1 mg/ml, levels that may have generated greater (dpH_i_/dt)_Max_ signals. In our bCA study, we did not explore hCA II levels greater than 1 ng, levels that presumably would have generated larger ΔpH_S_ signals.

Unlike the trio of papers noted above, the present study does not include assessments of altered extracellular concentrations of (1) CO_2_ (1.5%, 5%, 10% vs. 1.5% in present study) or (2) non-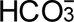 buffers (1, 5, 25 mM HEPES vs. 5 mM here). Including such analyses in the present study, together with ±hAQP5, would have expanded our experimental matrix to unrealistic levels.

Musa-Aziz et al. (2014b) found that (1) increasing [CO_2_]_o_ markedly increased both ΔpH_S_ and (dpH_i_/dt)_Max_, especially in the presence of hCA IV (their figure 10). The effects of increasing [HEPES]_o_ were more complex: (2) With 1.5% CO_2_, raising [HEPES]_o_ predictably decreased ΔpH_S_ (a “cis-side” effect) but did not significantly affect (dpH_i_/dt)_Max_ (their figure 13). (3) When they supplemented the hCA IV with extracellular bCA II, still with 1.5% CO_2_, the effect of raising [HEPES]_o_ on (dpH_i_/dt)_Max_ became stronger but still not significant (their figure 15). However, (4) when they work with 10% CO_2_, the effects of raising [HEPES]_o_ on (dpH_i_/dt)_Max_ were far stronger (their fig 17).

Based on the above observations, we suggest the following for future (dpH_i_/dt)_Max_ experiments on oocytes intended to examine the synergy among hAQP5, CA_i_, and: In ΔΔ(dpH_i_/dt)_Max_ protocols, it would be helpful to: (a) raise [CO_2_]_o_ (e.g., to 10%), (b) explore increasing bCA beyond 0.1 mg/ml, and (c) raise [HEPES]_o_ (e.g., to 25 mM). We also suggest that, in future ΔΔΔpH_S_ protocols, it would be helpful to (a) raise the [CO_2_]_o_ and (b) increase CA_i_ activity (e.g., employing recombinant hCA II, injecting 100 ng or more of purified hCA II).

## Additional information

### Authors’ contributions

D.K.W. contributed to the conception and design of the research, performed the experiments, analyzed data, and interpreted results, wrote the first draft of manuscript, prepared the figures, and edited the manuscript. F.J.M. contributed to the conception and design of the research, interpreted results, performed statistical analyses and edited the figures and manuscript. W.F.B. contributed to the conception and design of the research, interpreted results, and edited the figures and manuscript. All authors approved the final version of the manuscript, all qualify for authorship, and all those who qualify for authorship are listed.

## Acknowledgements

We thank Dale E. Huffman for computer support. We acknowledge the assistance of Gerald T. Babcock in his role as laboratory manager. W.F.B. gratefully acknowledges the support of the Myers/Scarpa endowed chair.

## Statistics Tables

**Statistics Table 3.** For clarity within the figure panel, we present tables of *P*-values for one-way ANOVA with Tukey’s post-hoc means comparison for the data presented in Figure 3. For all tables, α is 0.05 and significant *P*-values are highlighted by bolding. *A, P-*values for means comparisons of ΔpH_S_ upon CO_2_ addition data. *B, P*-values for means comparisons of ΔpH_S_ upon CO_2_ removal data. *C, P-*values for means comparisons of (dpH_i_/dt)_Max_ upon CO_2_ addition data. *D, P-*values for means comparisons of (dpH_i_/dt)_Max_ upon CO_2_ removal data.

| cRNA |  | H<sub>2</sub>O | hAQP5 | H<sub>2</sub>O | hAQP5 | H<sub>2</sub>O | hAQP5 | H<sub>2</sub>O |
| --- | --- | --- | --- | --- | --- | --- | --- | --- |
|  | hCA II (ng) | "Tris" | "Tris" | 1 | 1 | 10 | 10 | 100 |
| hAQP5 | "Tris" | **0.00150** |  |  |  |  |  |  |
| H<sub>2</sub>O | 1 | 0.956 | **0.0391** |  |  |  |  |  |
| hAQP5 | 1 | **4.18×10<sup>-8</sup>** | **0.00247** | **7.19×10<sup>-8</sup>** |  |  |  |  |
| H<sub>2</sub>O | 10 | 0.943 | **0.0456** | 1.00 | **7.72×10<sup>-8</sup>** |  |  |  |
| hAQP5 | 10 | **3.90×10<sup>-8</sup>** | **7.38×10<sup>-4</sup>** | **5.43×10<sup>-8</sup>** | 1.00 | **5.60×10<sup>-8</sup>** |  |  |
| H<sub>2</sub>O | 100 | 0.947 | **0.0437** | 1.00 | **7.56×10<sup>-8</sup>** | 1.00 | **5.55×10<sup>-8</sup>** |  |
| hAQP5 | 100 | **2.21×10<sup>-8</sup>** | **1.34×10<sup>-7</sup>** | **2.93×10<sup>-8</sup>** | 0.0978 | **2.97×10<sup>-8</sup>** | 0.214 | **2.95×10<sup>-8</sup>** |

3B, $\Delta pH_s$ upon $CO_2$ removal
| cRNA |  | H<sub>2</sub>O | hAQP5 | H<sub>2</sub>O | hAQP5 | H<sub>2</sub>O | hAQP5 | H<sub>2</sub>O |
| --- | --- | --- | --- | --- | --- | --- | --- | --- |
|  | hCA II (ng) | "Tris" | "Tris" | 1 | 1 | 10 | 10 | 100 |
| hAQP5 | "Tris" | **2.03×10<sup>-4</sup>** |  |  |  |  |  |  |
| H<sub>2</sub>O | 1 | 0.992 | **0.00309** |  |  |  |  |  |
| hAQP5 | 1 | **4.44×10<sup>-8</sup>** | **0.0321** | **7.16×10<sup>-8</sup>** |  |  |  |  |
| H<sub>2</sub>O | 10 | 0.995 | **0.00249** | 1.00 | **6.67×10<sup>-8</sup>** |  |  |  |
| hAQP5 | 10 | **4.51×10<sup>-8</sup>** | **0.0383** | **7.76×10<sup>-8</sup>** | 1.00 | **7.15×10<sup>-8</sup>** |  |  |
| H<sub>2</sub>O | 100 | 0.994 | **0.00278** | 1.00 | **6.91×10<sup>-8</sup>** | 1.00 | **7.44×10<sup>-8</sup>** |  |
| hAQP5 | 100 | **3.17×10<sup>-8</sup>** | **1.50×10<sup>-4</sup>** | **3.72×10<sup>-8</sup>** | 0.692 | **3.68×10<sup>-8</sup>** | 0.650 | **3.70×10<sup>-8</sup>** |

3C, $(dpH_i/dt)_{Max}$ upon CO<sub>2</sub> addition
| cRNA |  | H<sub>2</sub>O | hAQP5 | H<sub>2</sub>O | hAQP5 | H<sub>2</sub>O | hAQP5 | H<sub>2</sub>O |
| --- | --- | --- | --- | --- | --- | --- | --- | --- |
|  | hCA II<br>(ng) | “Tris” | “Tris” | 1 | 1 | 10 | 10 | 100 |
| hAQP5 | “Tris” | 1.00 |  |  |  |  |  |  |
| H<sub>2</sub>O | 1 | 0.0803 | 0.0744 |  |  |  |  |  |
| hAQP5 | 1 | **0.00798** | **0.00726** | 0.989 |  |  |  |  |
| H<sub>2</sub>O | 10 | **3.62×10<sup>-4</sup>** | **3.25×10<sup>-4</sup>** | 0.624 | 0.979 |  |  |  |
| hAQP5 | 10 | **3.76×10<sup>-5</sup>** | **3.36×10<sup>-5</sup>** | 0.244 | 0.753 | 0.998 |  |  |
| H<sub>2</sub>O | 100 | **5.69×10<sup>-4</sup>** | **5.10×10<sup>-4</sup>** | 0.708 | 0.991 | 1.00 | 0.994 |  |
| hAQP5 | 100 | **0.00210** | **0.00189** | 0.905 | 1.00 | 0.999 | 0.938 | 1.00 |

3D, $(dpH_i/dt)_{Max}$ upon CO<sub>2</sub> removal
| cRNA |  | H<sub>2</sub>O | hAQP5 | H<sub>2</sub>O | hAQP5 | H<sub>2</sub>O | hAQP5 | H<sub>2</sub>O |
| --- | --- | --- | --- | --- | --- | --- | --- | --- |
|  | hCA II<br>(ng) | “Tris” | “Tris” | 1 | 1 | 10 | 10 | 100 |
| hAQP5 | “Tris” | 1.00 |  |  |  |  |  |  |
| H<sub>2</sub>O | 1 | 0.154 | 0.141 |  |  |  |  |  |
| hAQP5 | 1 | **0.0330** | **0.0295** | 0.998 |  |  |  |  |
| H<sub>2</sub>O | 10 | **0.00381** | **0.00335** | 0.865 | 0.995 |  |  |  |
| hAQP5 | 10 | **4.57×10<sup>-5</sup>** | **3.97×10<sup>-4</sup>** | 0.479 | 0.862 | 0.998 |  |  |
| H<sub>2</sub>O | 100 | **0.00160** | **0.00140** | 0.721 | 0.972 | 1.00 | 1.00 |  |
| hAQP5 | 100 | **0.0460** | **0.0413** | 1.00 | 1.00 | 0.988 | 0.801 | 0.947 |

**Statistics Table 5.** For clarity within the figure panel, we present tables of *P*-values for one-way ANOVA with Tukey’s post-hoc means comparison for the data presented in Figure 5. For all tables, α is 0.05 and significant *P*-values are highlighted by bolding. *A, P-*values for means comparisons of (dpH_S_/dt)_Max_ upon CO_2_ addition. *B, P*-values for means comparisons of (dpH_S_/dt)_Max_ upon CO_2_ removal. Where the displayed *P*-values is 0.00, this indicates a values less than <2.22×10^−308^. ^12^

| cRNA |  | H<sub>2</sub>O | hAQP5 | H<sub>2</sub>O | hAQP5 | H<sub>2</sub>O | hAQP5 | H<sub>2</sub>O |
| --- | --- | --- | --- | --- | --- | --- | --- | --- |
|  | hCA II (ng) | "Tris" | "Tris" | 1 | 1 | 10 | 10 | 100 |
| hAQP5 | "Tris" | **$4.79 \times 10^{-4}$** |  |  |  |  |  |  |
| H<sub>2</sub>O | 1 | 0.998 | **0.00518** |  |  |  |  |  |
| hAQP5 | 1 | **$4.73 \times 10^{-8}$** | **0.104** | **$7.84 \times 10^{-8}$** |  |  |  |  |
| H<sub>2</sub>O | 10 | 0.975 | **0.00655** | 1.00 | **$4.77 \times 10^{-8}$** |  |  |  |
| hAQP5 | 10 | **$3.42 \times 10^{-8}$** | **$7.41 \times 10^{-4}$** | **$4.18 \times 10^{-8}$** | 0.408 | **$4.03 \times 10^{-8}$** |  |  |
| H<sub>2</sub>O | 100 | 0.993 | **0.0134** | 1.00 | **$4.89 \times 10^{-7}$** | 1.00 | **$4.78 \times 10^{-8}$** |  |
| hAQP5 | 100 | **$2.85 \times 10^{-8}$** | **0.00** | **$2.85 \times 10^{-8}$** | **$4.74 \times 10^{-8}$** | **0.00** | **$3.95 \times 10^{-4}$** | **0.00** |

5B, (dpH<sub>s</sub>/dt)<sub>Max</sub> upon CO<sub>2</sub> removal
| cRNA |  | H<sub>2</sub>O | hAQP5 | H<sub>2</sub>O | hAQP5 | H<sub>2</sub>O | hAQP5 | H<sub>2</sub>O |
| --- | --- | --- | --- | --- | --- | --- | --- | --- |
|  | hCA II (ng) | "Tris" | "Tris" | 1 | 1 | 10 | 10 | 100 |
| hAQP5 | "Tris" | **0.0151** |  |  |  |  |  |  |
| H<sub>2</sub>O | 1 | 0.999 | 0.0862 |  |  |  |  |  |
| hAQP5 | 1 | **$7.67 \times 10^{-4}$** | 0.998 | **0.00783** |  |  |  |  |
| H<sub>2</sub>O | 10 | 0.995 | 0.0768 | 1.00 | **0.00516** |  |  |  |
| AQP5 | 10 | **0.00116** | 0.992 | **0.0125** | 1.00 | **0.00975** |  |  |
| H<sub>2</sub>O | 100 | 0.995 | 0.165 | 1.00 | **0.00228** | 1.00 | **0.0303** |  |
| AQP5 | 100 | **$3.68 \times 10^{-8}$** | **0.0114** | **$4.03 \times 10^{-7}$** | **0.0280** | **$1.21 \times 10^{-7}$** | 0.144 | **$2.06 \times 10^{-6}$** |

<sup>12</sup> $2.22 \times 10^{-308}$ is the smallest possible value for double type data that a 64-bit system is able to distinguish

**Statistics Table 6.** For clarity within the figure panel, we present tables of *P*-values for one-way ANOVA with Tukey’s post-hoc means comparison for the data presented in Figure 6. For all tables, α is 0.05 and significant *P*-values are highlighted by bolding. *A, P*-values for means comparisons of initial pH_i_ data. *B, P*-values for means comparisons of ΔpH_i_ data. *C, P-* values for means comparisons intrinsic buffering power (β_I_). *D, P*-values for means comparisons of *P*_f_ data.

| cRNA |  | H<sub>2</sub>O | hAQP5 | H<sub>2</sub>O | hAQP5 | H<sub>2</sub>O | hAQP5 | H<sub>2</sub>O |
| --- | --- | --- | --- | --- | --- | --- | --- | --- |
|  | hCA II (ng) | "Tris" | "Tris" | 1 | 1 | 10 | 10 | 100 |
| hAQP5 | "Tris" | 0.701 |  |  |  |  |  |  |
| H<sub>2</sub>O | 1 | 0.971 | 0.997 |  |  |  |  |  |
| hAQP5 | 1 | 0.512 | 1.00 | 0.978 |  |  |  |  |
| H<sub>2</sub>O | 10 | 1.00 | 0.481 | 0.877 | 0.309 |  |  |  |
| hAQP5 | 10 | 0.703 | 1.00 | 0.998 | 1.00 | 0.484 |  |  |
| H<sub>2</sub>O | 100 | 1.00 | 0.569 | 0.925 | 0.385 | 1.00 | 0.572 |  |
| hAQP5 | 100 | 0.997 | 0.972 | 1.00 | 0.898 | 0.970 | 0.973 | 0.987 |

6B ΔpH<sub>i</sub>
| cRNA |  | H<sub>2</sub>O | hAQP5 | H<sub>2</sub>O | hAQP5 | H<sub>2</sub>O | hAQP5 | H<sub>2</sub>O |
| --- | --- | --- | --- | --- | --- | --- | --- | --- |
|  | hCA II (ng) | "Tris" | "Tris" | 1 | 1 | 10 | 10 | 100 |
| hAQP5 | "Tris" | 1.00 |  |  |  |  |  |  |
| H<sub>2</sub>O | 1 | 1.00 | 1.00 |  |  |  |  |  |
| hAQP5 | 1 | 0.679 | 0.909 | 0.757 |  |  |  |  |
| H<sub>2</sub>O | 10 | 0.980 | 1.00 | 0.991 | 0.994 |  |  |  |
| hAQP5 | 10 | 0.0737 | 0.199 | 0.0995 | 0.902 | 0.460 |  |  |
| H<sub>2</sub>O | 100 | 1.00 | 1.00 | 1.00 | 0.748 | 0.990 | 0.0963 |  |
| hAQP5 | 100 | 0.480 | 0.767 | 0.563 | 1.00 | 0.960 | 0.976 | 0.554 |

6C $\beta_i$
| cRNA |  | H<sub>2</sub>O | hAQP5 | H<sub>2</sub>O | hAQP5 | H<sub>2</sub>O | hAQP5 | H<sub>2</sub>O |
| --- | --- | --- | --- | --- | --- | --- | --- | --- |
|  | hCA II (ng) | “Tris” | “Tris” | 1 | 1 | 10 | 10 | 100 |
| hAQP5 | “Tris” | 0.982 |  |  |  |  |  |  |
| H<sub>2</sub>O | 1 | 0.995 | 1.00 |  |  |  |  |  |
| hAQP5 | 1 | 1.00 | 0.869 | 0.934 |  |  |  |  |
| H<sub>2</sub>O | 10 | 0.876 | 0.326 | 0.434 | 0.984 |  |  |  |
| hAQP5 | 10 | 0.504 | 0.0905 | 0.137 | 0.782 | 0.998 |  |  |
| H<sub>2</sub>O | 100 | 1.00 | 0.928 | 0.970 | 1.00 | 0.960 | 0.685 |  |
| hAQP5 | 100 | 0.720 | 0.188 | 0.267 | 0.926 | 1.00 | 1.00 | 0.867 |

6D $P_f$
| cRNA |  | H<sub>2</sub>O | hAQP5 | H<sub>2</sub>O | hAQP5 | H<sub>2</sub>O | hAQP5 | H<sub>2</sub>O |
| --- | --- | --- | --- | --- | --- | --- | --- | --- |
|  | hCA II (ng) | “Tris” | “Tris” | 1 | 1 | 10 | 10 | 100 |
| hAQP5 | “Tris” | **6.86×10<sup>-8</sup>** |  |  |  |  |  |  |
| H<sub>2</sub>O | 1 | 1.00 | **8.93×10<sup>-7</sup>** |  |  |  |  |  |
| hAQP5 | 1 | **1.87×10<sup>-6</sup>** | 1.00 | **2.79×10<sup>-7</sup>** |  |  |  |  |
| H<sub>2</sub>O | 10 | 1.00 | **8.76×10<sup>-7</sup>** | 1.00 | **8.85×10<sup>-7</sup>** |  |  |  |
| hAQP5 | 10 | **7.30×10<sup>-8</sup>** | 0.988 | **7.23×10<sup>-8</sup>** | 0.996 | **7.16×10<sup>-8</sup>** |  |  |
| H<sub>2</sub>O | 100 | 1.00 | **8.96×10<sup>-7</sup>** | 1.00 | **6.86×10<sup>-8</sup>** | 1.00 | **7.25×10<sup>-8</sup>** |  |
| hAQP5 | 100 | **7.08×10<sup>-8</sup>** | 1.00 | **1.86×10<sup>-6</sup>** | 1.00 | **8.93×10<sup>-7</sup>** | 0.999 | **1.87×10<sup>-6</sup>** |

**Statistics Table 8.** For clarity within the figure panel, we present tables of *P*-values for one-way ANOVA with Tukey’s post-hoc means comparison for the data presented in Figure 8. For all tables, α is 0.05 and significant *P*-values are highlighted by bolding. *A, P*-values for means comparisons of initial pH_i_ data. *B, P*-values for means comparisons of ΔpH_i_ data. *C, P-* values for means comparisons of β_I_. *D, P*-values for means comparisons of *P*_f_ data. Panels *E-H* provide descriptive statistics and means comparisons (see Methods, Statistical analysis) for comparisons of data represented by the gray, black light-blue and dark-blue bars in Figure 3 (–bCA) vs. Figure 8 (+bCA). *E*, ΔpH_i_ during addition of CO_2_/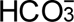. *F*, ΔpH_i_ during removal of CO_2_/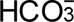. *G*, (dpH_i_/dt)_Max_ during addition of CO_2_/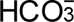. *H*, (dpH_i_/dt)_Max_

| cRNA |  | H<sub>2</sub>O | hAQP5 | H<sub>2</sub>O |
| --- | --- | --- | --- | --- |
|  | hCA II (ng) | “Tris” | “Tris” | 1 |
| hAQP5 | “Tris” | 1.00 |  |  |
| H<sub>2</sub>O | 1 | **0.00582** | **0.00663** |  |
| hAQP5 | 1 | **6.13×10<sup>–4</sup>** | **6.99×10<sup>–4</sup>** | 0.757 |

8B ΔpH<sub>i</sub> CO<sub>2</sub> removal
| cRNA |  | H<sub>2</sub>O | hAQP5 | H<sub>2</sub>O |
| --- | --- | --- | --- | --- |
|  | hCA II (ng) | “Tris” | “Tris” | 1 |
| hAQP5 | “Tris” | 0.996 |  |  |
| H<sub>2</sub>O | 1 | **0.0352** | 0.0560 |  |
| hAQP5 | 1 | **0.00287** | **0.00478** | 0.674 |

8C $(dpH/dt)_{Max}$ CO<sub>2</sub> addition
| cRNA |  | H<sub>2</sub>O | hAQP5 | H<sub>2</sub>O |
| --- | --- | --- | --- | --- |
|  | hCA II (ng) | “Tris” | “Tris” | 1 |
| hAQP5 | “Tris” | **0.0331** |  |  |
| H<sub>2</sub>O | 1 | **2.42×10<sup>-6</sup>** | 0.00171 |  |
| hAQP5 | 1 | **2.33×10<sup>-5</sup>** | 0.0192 | 0.707 |

8D $(dpH/dt)_{Max}$ CO<sub>2</sub> removal
| cRNA |  | H<sub>2</sub>O | hAQP5 | H<sub>2</sub>O |
| --- | --- | --- | --- | --- |
|  | hCA II (ng) | “Tris” | “Tris” | 1 |
| hAQP5 | “Tris” | 0.571 |  |  |
| H<sub>2</sub>O | 1 | **1.42×10<sup>-5</sup>** | **2.49×10<sup>-4</sup>** |  |
| hAQP5 | 1 | **7.08×10<sup>-4</sup>** | **0.0135** | 0.318 |

8E, $\Delta pH_s$ upon CO<sub>2</sub> addition ± bCA in bECF (Figure 3A vs. Figure 8A)
| Descriptive Stats. | n | Mean | S.D. | S.E.M |  |  |  |  |
| --- | --- | --- | --- | --- | --- | --- | --- | --- |
| bECF | 32 | 0.0689 | 0.0422 | 0.00746 |  |  |  |  |
| bECF + bCA | 24 | 0.286 | 0.0641 | 0.0131 |  |  |  |  |
| Means comparison |  | Mean Diff. | S.E.M. | Q-value | P-value | α | LCL | UCL |
| bECF vs. bECF + bCA |  | 0.217 | 0.0142 | 21.6 | 4.53×10<sup>-8</sup> | 0.0500 | 0.188 | 0.245 |

8F, $\Delta pH_s$ upon CO<sub>2</sub> removal ± bCA in bECF (Figure 3B vs. Figure 8B)
| Descriptive Stats. | n | Mean | S.D. | S.E.M |  |  |  |  |
| --- | --- | --- | --- | --- | --- | --- | --- | --- |
| bECF | 32 | -0.0771 | 0.0487 | 0.00861 |  |  |  |  |
| bECF + bCA | 24 | -0.386 | 0.0681 | 0.0139 |  |  |  |  |
| Means comparison | | Mean Diff. | S.E.M. | Q-value | P-value | $\alpha$ | LCL | UCL |
| bECF vs. bECF + bCA | | -0.309 | 0.0156 | 28.0 | **4.53**$\times 10^{-8}$ | 0.0500 | -0.340 | -0.278 |

8G, $(dpH/dt)_{Max}$ upon CO<sub>2</sub> addition ± bCA in bECF (Figure 3C vs. Figure 8C)
| Descriptive Stats. | n | Mean | S.D. | S.E.M |  |  |  |  |
| --- | --- | --- | --- | --- | --- | --- | --- | --- |
| bECF | 32 | −0.00315 | 0.00236 | 4.18×10<sup>−4</sup> |  |  |  |  |
| bECF + bCA | 24 | −0.00528 | 0.00225 | 4.60×10<sup>−4</sup> |  |  |  |  |
| Means comparison |  | Mean Diff. | S.E.M. | Q-value | P-value | α | LCL | UCL |
| bECF vs. bECF + bCA |  | −0.00213 | 6.25×10<sup>−4</sup> | 4.83 | **0.00122** | 0.0500 | −0.00339 | −8.81×10<sup>−4</sup> |

$8H, (dpH/dt)_{Max}$ upon CO<sub>2</sub> removal ± bCA in bECF (Figure 3D vs. Figure 8D)
| **Descriptive Stats.** | **n** | **Mean** | **S.D.** | **S.E.M** |  |  |  |  |
| --- | --- | --- | --- | --- | --- | --- | --- | --- |
| bECF | 32 | 0.00183 | 0.00125 | $2.22 \times 10^{-4}$ | | | | |
| bECF + bCA | 24 | 0.00323 | 0.00161 | $3.29 \times 10^{-4}$ | | | | |
| **Means comparison** |  | **Mean Diff.** | **S.E.M.** | **Q-value** | **P-value** | **$\alpha$** | **LCL** | **UCL** |
| bECF vs. bECF + bCA | | 0.00140 | $3.83 \times 10^{-4}$ | 5.17 | **$5.82 \times 10^{-7}$** | 0.0500 | $6.32 \times 10^{-4}$ | 0.00217 |

**Statistics Table 9.** For clarity within the figure panel, we present tables of *P*-values for one-way ANOVA with Tukey’s post-hoc means comparison for the data presented in Figure 9. For all tables, α is 0.05 and significant *P*-values are highlighted by bolding. *A, P*-values for means comparisons of (dpH_S_/dt)_Max_ on CO_2_ addition. *B, P*-values for means comparisons of (dpH_S_/dt)_Max_ on CO_2_ removal. *C, P-*values for means comparisons of β_i_. *D, P*-values for means comparisons of *P*_f_ data.

| cRNA |  | H<sub>2</sub>O | hAQP5 | H<sub>2</sub>O |
| --- | --- | --- | --- | --- |
|  | hCA II (ng) | "Tris" | "Tris" | 1 |
| hAQP5 | "Tris" | 0.960 |  |  |
| H<sub>2</sub>O | 1 | **$3.38 \times 10^{-4}$** | **0.00103** |  |
| hAQP5 | 1 | **0.00221** | **0.00674** | 0.842 |

9B $(dpH_s/dt)_{Max}$ on CO<sub>2</sub> removal
| cRNA |  | H<sub>2</sub>O | hAQP5 | H<sub>2</sub>O |
| --- | --- | --- | --- | --- |
|  | hCA II (ng) | "Tris" | "Tris" | 1 |
| hAQP5 | "Tris" | **0.0288** |  |  |
| H<sub>2</sub>O | 1 | 0.625 | 0.281 |  |
| hAQP5 | 1 | 0.0604 | 0.984 | 0.461 |

**Statistics Table 10.** For clarity within the figure panel, we present tables of *P*-values for one-way ANOVA with Tukey’ s post-hoc means comparison for the data presented in Figure 10. For all tables, α is 0.05 and significant *P*-values are highlighted by bolding. *A, P*-values for means comparisons of initial pH_i_ data. *B, P*-values for means comparisons of ΔpH_i_ data. *C, P-*values for means comparisons of β_I_. *D, P*-values for means comparisons of *P*_f_ data. Panels *E-H* provide descriptive statistics and means comparisons (see Methods, Statistical analysis) for comparisons of data represented by the gray, black light-blue and dark-blue bars in Figure 6 (–bCA) vs. Figure 10 (+bCA). *E*, Initial pH_i_. *F*, ΔpH_i_ amplitude on addition of CO_2_/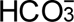. *G*, β_I_. *H*, *P*_f_.

| cRNA |  | H<sub>2</sub>O | hAQP5 | H<sub>2</sub>O |
| --- | --- | --- | --- | --- |
|  | hCA II (ng) | “Tris” | “Tris” | 1 |
| hAQP5 | “Tris” | 0.752 |  |  |
| H<sub>2</sub>O | 1 | 1.00 | 0.727 |  |
| hAQP5 | 1 | 0.939 | 0.974 | 0.925 |

*10B ΔpH<sub>i</sub> data*

| cRNA |  | H<sub>2</sub>O | hAQP5 | H<sub>2</sub>O |
| --- | --- | --- | --- | --- |
|  | hCA II (ng) | “Tris” | “Tris” | 1 |
| hAQP5 | “Tris” | 0.798 |  |  |
| H<sub>2</sub>O | 1 | 1.00 | 0.779 |  |
| hAQP5 | 1 | 0.486 | 0.949 | 0.465 |

10C Buffering power ( $\beta_i$ )
| cRNA |  | H<sub>2</sub>O | hAQP5 | H<sub>2</sub>O |
| --- | --- | --- | --- | --- |
|  | hCA II (ng) | “Tris” | “Tris” | 1 |
| hAQP5 | “Tris” | 0.994 |  |  |
| H<sub>2</sub>O | 1 | 1.00 | 0.994 |  |
| hAQP5 | 1 | 0.652 | 0.799 | 0.659 |

10D $P_i$ data
| cRNA |  | H<sub>2</sub>O | hAQP5 | H<sub>2</sub>O |
| --- | --- | --- | --- | --- |
|  | hCA II (ng) | “Tris” | “Tris” | 1 |
| hAQP5 | “Tris” | $2.10 \times 10^{-12}$ | | |
| H<sub>2</sub>O | 1 | 0.999 | $2.10 \times 10^{-12}$ | |
| hAQP5 | 1 | $2.10 \times 10^{-12}$ | 0.999 | $2.10 \times 10^{-12}$ |

10E, initial $pH_i$ values ± bCA in bECF (Figure 6A vs. Figure 10A)
| Descriptive Stats. | n | Mean | S.D. | S.E.M |  |  |  |  |
| --- | --- | --- | --- | --- | --- | --- | --- | --- |
| bECF | 32 | 7.30 | 0.0551 | 0.00975 |  |  |  |  |
| bECF + bCA | 24 | 7.31 | 0.0603 | 0.0123 |  |  |  |  |
| Means comparison | | Mean Diff. | S.E.M. | Q-value | P-value | $\alpha$ | LCL | UCL |
| bECF vs. bECF + bCA |  | 0.00669 | 0.0155 | 0.610 | 0.668 | 0.0500 | -0.0244 | 0.0378 |

10F, $\Delta pH_i$ upon CO<sub>2</sub> addition ± bCA in bECF (Figure 6B vs. Figure 10B)
| Descriptive Stats. | n | Mean | S.D. | S.E.M |  |  |  |  |
| --- | --- | --- | --- | --- | --- | --- | --- | --- |
| bECF | 32 | -0.258 | 0.0258 | 0.00456 |  |  |  |  |
| bECF + bCA | 24 | -0.285 | 0.0308 | 0.00628 |  |  |  |  |
| Means comparison | | Mean Diff. | S.E.M. | Q-value | P-value | $\alpha$ | LCL | UCL |
| bECF vs. bECF + bCA | | -0.0270 | 0.00756 | 5.04 | $7.74 \times 10^{-4}$ | 0.0500 | 0.0118 | 0.0421 |

10G, $\beta$ values for oocytes ± bCA in bECF (Figure 6C vs. Figure 10C)
| **Descriptive Stats.** | **n** | **Mean** | **S.D.** | **S.E.M** |  |  |  |  |
| --- | --- | --- | --- | --- | --- | --- | --- | --- |
| bECF | 32 | 13.6 | 1.93 | 0.341 |  |  |  |  |
| bECF + bCA | 24 | 11.6 | 1.73 | 0.353 |  |  |  |  |
| **Means comparison** |  | **Mean Diff.** | **S.E.M.** | **Q-value** | **P-value** | **$\alpha$** | **LCL** | **UCL** |
| bECF vs. bECF + bCA |  | -2.03 | 0.498 | 5.75 | **$1.55 \times 10^{-4}$** | 0.0500 | -3.02 | -1.03 |

10H, $P_i \pm$ bCA in bECF (Figure 6D vs. Figure 10D)
| **Descriptive Stats.** | **n** | **Mean** | **S.D.** | **S.E.M** |  |  |  |  |
| --- | --- | --- | --- | --- | --- | --- | --- | --- |
| bECF | 32 | 0.00196 | 0.00184 | $3.25 \times 10^{-4}$ | | | | |
| bECF + bCA | 30 | 0.00204 | 0.00192 | $3.50 \times 10^{-4}$ | | | | |
| **Means comparison** |  | **Mean Diff.** | **S.E.M.** | **Q-value** | **P-value** | **$\alpha$** | **LCL** | **UCL** |
| bECF vs. bECF + bCA | | $8.03 \times 10^{-5}$ | $4.77 \times 10^{-4}$ | 0.238 | 0.867 | 0.0500 | $-8.74 \times 10^{-4}$ | 0.00103 |

## Supplemental Statistics Tables

**Supplemental Table 3A.** Statistics Summary for Figure 3*A*, i) Descriptive Statistics for data presented in Figure 3*A*, ΔpH_S_ upon CO_2_ addition. ii) Overall one-way ANOVA results for data presented in Figure 3*A*. DF is degrees of freedom. iii) Tukey’s means comparison analysis for all data presented in Figure 3*A*. Threshold for significance α=0.05. Standard error of the mean = 0.0116. Mean difference for each comparison (Mean Diff.), lower confidence limit (LCL), upper confidence limit (UCL).

i. Descriptive Statistics
|  | n | Mean | Standard Deviation | SE of Mean |
| --- | --- | --- | --- | --- |
| H<sub>2</sub>O + Tris | 8 | 0.0285 | 0.00895 | 0.00316 |
| hAQP5 + Tris | 8 | 0.0787 | 0.0181 | 0.00641 |
| H<sub>2</sub>O + 1 ng hCA II | 8 | 0.0411 | 0.0115 | 0.00406 |
| hAQP5 + 1 ng hCA II | 8 | 0.127 | 0.0246 | 0.00869 |
| H<sub>2</sub>O + 10 ng hCA II | 8 | 0.0418 | 0.00938 | 0.00332 |
| hAQP5 + 10 ng hCA II | 8 | 0.131 | 0.0262 | 0.00925 |
| H<sub>2</sub>O + 100 ng hCA II | 8 | 0.0416 | 0.00948 | 0.00335 |
| hAQP5 + 100 ng hCA II | 8 | 0.160 | 0.0478 | 0.0169 |

ii. Overall ANOVA Table
| Variation | DF | Sum of Squares | Mean Square | F-value | P-value |
| --- | --- | --- | --- | --- | --- |
| Between Groups | 7 | 0.147 | 0.0210 | 39.2 | $2.44 \times 10^{-19}$ |
| Within Groups | 56 | 0.0301 | $5.37 \times 10^{-4}$ | | |
| Total | 63 | 0.177 |  |  |  |

iii. Tukey Test
| Condition A | Condition B | Mean Diff. | Q-value | P-value | Sig. | LCL | UCL |
| --- | --- | --- | --- | --- | --- | --- | --- |
| hAQP5 + Tris | H<sub>2</sub>O + Tris | 0.0502 | 6.13 | 0.00150 | ✓ | 0.0137 | 0.0867 |
| H<sub>2</sub>O + 1 ng hCA II | H<sub>2</sub>O + Tris | 0.0126 | 1.54 | 0.956 | – | –0.0238 | 0.0491 |
| H<sub>2</sub>O + 1 ng hCA II | hAQP5 + Tris | –0.0376 | 4.59 | 0.0391 | ✓ | –0.0741 | –0.00110 |
| hAQP5 + 1 ng hCA II | H<sub>2</sub>O + Tris | 0.0986 | 12.0 | $4.18 \times 10^{-8}$ | ✓ | 0.0622 | 0.135 |
| hAQP5 + 1 ng hCA II | hAQP5 + Tris | 0.0484 | 5.91 | 0.00247 | ✓ | 0.0120 | 0.0849 |
| hAQP5 + 1 ng hCA II | H<sub>2</sub>O + 1ng hCA II | 0.0860 | 10.5 | $7.19 \times 10^{-8}$ | ✓ | 0.0495 | 0.122 |
| H<sub>2</sub>O + 10 ng hCA II | H<sub>2</sub>O + Tris | 0.0133 | 1.63 | 0.943 | – | –0.0232 | 0.0498 |
| H<sub>2</sub>O + 10 ng hCA II | hAQP5 + Tris | –0.0369 | 4.50 | 0.0456 | ✓ | –0.0734 | $-4.10 \times 10^{-4}$ |
| H<sub>2</sub>O + 10 ng hCA II | H<sub>2</sub>O + 1ng hCA II | $6.88 \times 10^{-4}$ | 0.0839 | 1.00 | – | –0.0358 | 0.0372 |
| H<sub>2</sub>O + 10 ng hCA II | hAQP5 + 1ng hCA II | –0.0853 | 10.4 | $7.72 \times 10^{-8}$ | ✓ | –0.122 | –0.0488 |
| hAQP5 + 10 ng hCA II | H<sub>2</sub>O + Tris | 0.103 | 12.6 | $3.90 \times 10^{-8}$ | ✓ | 0.0664 | 0.139 |
| hAQP5 + 10 ng hCA II | hAQP5 + Tris | 0.0527 | 6.43 | $7.38 \times 10^{-4}$ | ✓ | 0.0162 | 0.0892 |
| hAQP5 + 10 ng hCA II | H<sub>2</sub>O + 1 ng hCA II | 0.0902 | 11.0 | $5.43 \times 10^{-8}$ | ✓ | 0.0538 | 0.127 |
| hAQP5 + 10 ng hCA II | hAQP5 + 1 ng hCA II | 0.00424 | 0.518 | 1.00 | – | –0.0322 | 0.0407 |
| hAQP5 + 10 ng hCA II | H<sub>2</sub>O + 10 ng hCA II | 0.0896 | 10.9 | $5.60 \times 10^{-8}$ | ✓ | 0.0531 | 0.126 |
| H<sub>2</sub>O + 100 ng hCA II | H<sub>2</sub>O + Tris | 0.0131 | 1.60 | 0.947 | – | –0.0234 | 0.0496 |
| H<sub>2</sub>O + 100 ng hCA II | hAQP5 + Tris | –0.0371 | 4.53 | 0.0437 | ✓ | –0.0736 | $-6.01 \times 10^{-4}$ |
| H<sub>2</sub>O + 100 ng hCA II | H<sub>2</sub>O + 1 ng hCA II | $4.96 \times 10^{-4}$ | 0.0606 | 1.00 | – | –0.0360 | 0.0370 |
| H<sub>2</sub>O + 100 ng hCA II | hAQP5 + 1 ng hCA II | –0.0855 | 10.4 | $7.56 \times 10^{-8}$ | ✓ | –0.122 | –0.0490 |
| H<sub>2</sub>O + 100 ng hCA II | H<sub>2</sub>O + 10 ng hCA II | $-1.91 \times 10^{-4}$ | 0.0233 | 1.00 | – | –0.0367 | 0.0363 |
| H<sub>2</sub>O + 100 ng hCA II | hAQP5 + 10 ng hCA II | –0.0898 | 11.0 | $5.55 \times 10^{-8}$ | ✓ | –0.126 | –0.0533 |
| hAQP5 +100 ng hCA II | H<sub>2</sub>O + Tris | 0.132 | 16.1 | $2.21 \times 10^{-8}$ | ✓ | 0.0955 | 0.168 |
| hAQP5 +100 ng hCA II | hAQP5 + Tris | 0.0817 | 9.98 | $1.34 \times 10^{-7}$ | ✓ | 0.0453 | 0.118 |
| hAQP5 +100 ng hCA II | H<sub>2</sub>O + 1 ng hCA II | 0.119 | 14.6 | $2.93 \times 10^{-8}$ | ✓ | 0.0828 | 0.156 |
| hAQP5 +100 ng hCA II | hAQP5 + 1 ng hCA II | 0.0333 | 4.06 | 0.0978 | – | –0.00318 | 0.0698 |
| hAQP5 +100 ng hCA II | H<sub>2</sub>O + 10 ng hCA II | 0.119 | 14.5 | $2.97 \times 10^{-8}$ | ✓ | 0.0821 | 0.155 |
| hAQP5 +100 ng hCA II | hAQP5 + 10 ng hCA II | 0.0291 | 3.55 | 0.214 | – | –0.00742 | 0.0655 |
| hAQP5 +100 ng hCA II | H<sub>2</sub>O + 100 ng hCA II | 0.119 | 14.5 | $2.95 \times 10^{-8}$ | ✓ | 0.0823 | 0.155 |

**Supplemental Table 3B.** Statistics Summary for **Figure 3*B*, i**) Descriptive Statistics for data presented in Figure 3*B*, ΔpH_S_ upon CO_2_ removal. ii) Overall one-way ANOVA results for data presented in Figure 3*B*. DF is degrees of freedom. iii) Tukey’s means comparison analysis for all data presented in Figure 3*B*. Threshold for significance α=0.05. Standard error of the mean = 0.0132. Mean difference for each comparison (Mean Diff.), lower confidence limit (LCL), upper confidence limit (UCL).

|  | n | Mean | Standard Deviation | SE of Mean |
| --- | --- | --- | --- | --- |
| H<sub>2</sub>O + Tris | 8 | -0.03093 | 0.00994 | 0.00352 |
| hAQP5 + Tris | 8 | -0.09598 | 0.02688 | 0.0095 |
| H<sub>2</sub>O + 1 ng hCA II | 8 | -0.04167 | 0.01131 | 0.004 |
| hAQP5 + 1 ng hCA II | 8 | -0.13982 | 0.02817 | 0.00996 |
| H<sub>2</sub>O + 10 ng hCA II | 8 | -0.04078 | 0.01032 | 0.00365 |
| hAQP5 + 10 ng hCA II | 8 | -0.13894 | 0.03249 | 0.01149 |
| H<sub>2</sub>O + 100 ng hCA II | 8 | -0.04123 | 0.0107 | 0.00378 |
| hAQP5 + 100 ng hCA II | 8 | -0.16218 | 0.05069 | 0.01792 |

*ii. Overall ANOVA*
| Variation | DF | Sum of Squares | Mean Square | F-value | P-value |
| --- | --- | --- | --- | --- | --- |
| Between Groups | 7 | 0.165 | 0.0236 | 33.8 | $6.93 \times 10^{-18}$ |
| Within Groups | 56 | 0.0391 | $6.99 \times 10^{-4}$ | | |
| Total | 63 | 0.204 |  |  |  |

*iii. Tukey Test*
| Condition A | Condition B | Mean Diff. | Q-value | P-value | Sig. | LCL | UCL |
| --- | --- | --- | --- | --- | --- | --- | --- |
| hAQP5 + Tris | H<sub>2</sub>O + Tris | -0.0650 | 6.96 | $2.03 \times 10^{-4}$ | ✓ | -0.107 | -0.0234 |
| H<sub>2</sub>O + 1 ng hCA II | H<sub>2</sub>O + Tris | -0.0107 | 1.15 | 0.992 | – | -0.0524 | 0.0309 |
| H<sub>2</sub>O + 1 ng hCA II | hAQP5 + Tris | 0.0543 | 5.81 | 0.00309 | ✓ | 0.0127 | 0.0959 |
| hAQP5 + 1 ng hCA II | H<sub>2</sub>O + Tris | -0.109 | 11.7 | $4.44 \times 10^{-8}$ | ✓ | -0.150 | -0.0673 |
| hAQP5 + 1 ng hCA II | hAQP5 + Tris | -0.0438 | 4.69 | 0.0321 | ✓ | -0.0854 | -0.00223 |
| hAQP5 + 1 ng hCA II | H<sub>2</sub>O + 1 ng hCA II | -0.0981 | 10.5 | $7.16 \times 10^{-8}$ | ✓ | -0.140 | -0.0565 |
| H<sub>2</sub>O + 10 ng hCA II | H<sub>2</sub>O + Tris | -0.00985 | 1.05 | 0.995 | – | -0.0515 | 0.0318 |
| H<sub>2</sub>O + 10 ng hCA II | hAQP5 + Tris | 0.0552 | 5.91 | 0.00249 | ✓ | 0.0136 | 0.0968 |
| H<sub>2</sub>O + 10 ng hCA II | H<sub>2</sub>O + 1 ng hCA II | $8.96 \times 10^{-4}$ | 0.0959 | 1.00 | – | -0.0407 | 0.0425 |
| H<sub>2</sub>O + 10 ng hCA II | hAQP5 + 1 ng hCA II | 0.0990 | 10.6 | $6.67 \times 10^{-8}$ | ✓ | 0.0574 | 0.141 |
| hAQP5 + 10 ng hCA II | H<sub>2</sub>O + Tris | -0.108 | 11.6 | $4.51 \times 10^{-8}$ | ✓ | -0.150 | -0.0664 |
| hAQP5 + 10 ng hCA II | hAQP5 + Tris | -0.0430 | 4.60 | 0.0383 | ✓ | -0.0846 | -0.00136 |
| hAQP5 + 10 ng hCA II | H<sub>2</sub>O + 1 ng hCA II | -0.0973 | 10.4 | $7.76 \times 10^{-8}$ | ✓ | -0.139 | -0.0557 |
| hAQP5 + 10 ng hCA II | hAQP5 + 1 ng hCA II | $8.74 \times 10^{-4}$ | 0.0935 | 1.00 | - | -0.0407 | 0.0425 |
| hAQP5 + 10 ng hCA II | H<sub>2</sub>O + 10 ng hCA II | -0.0982 | 10.5 | $7.15 \times 10^{-8}$ | ✓ | -0.140 | -0.0566 |
| H<sub>2</sub>O + 100 ng hCA II | H<sub>2</sub>O + Tris | -0.0103 | 1.10 | 0.994 | - | -0.0519 | 0.0313 |
| H<sub>2</sub>O + 100 ng hCA II | hAQP5 + Tris | 0.0547 | 5.86 | 0.00278 | ✓ | 0.0131 | 0.0964 |
| H<sub>2</sub>O + 100 ng hCA II | H<sub>2</sub>O + 1 ng hCA II | $4.40 \times 10^{-4}$ | 0.0471 | 1.00 | - | -0.0412 | 0.0420 |
| H<sub>2</sub>O + 100 ng hCA II | hAQP5 + 1 ng hCA II | 0.0986 | 10.5 | $6.91 \times 10^{-8}$ | ✓ | 0.0570 | 0.140 |
| H<sub>2</sub>O + 100 ng hCA II | H<sub>2</sub>O + 10 ng hCA II | $-4.56 \times 10^{-4}$ | 0.0488 | 1.00 | - | -0.0421 | 0.0412 |
| H<sub>2</sub>O + 100 ng hCA II | hAQP5 + 10ng hCA II | 0.0977 | 10.5 | $7.44 \times 10^{-8}$ | ✓ | 0.0561 | 0.139 |
| hAQP5 +100 ng hCAII | H<sub>2</sub>O + Tris | -0.131 | 14.0 | $3.17 \times 10^{-8}$ | ✓ | -0.173 | -0.0896 |
| hAQP5 +100 ng hCA II | hAQP5 + Tris | -0.0662 | 7.08 | $1.50 \times 10^{-4}$ | ✓ | -0.108 | -0.0246 |
| hAQP5 +100 ng hCA II | H<sub>2</sub>O + 1 ng hCA II | -0.121 | 12.9 | $3.72 \times 10^{-8}$ | ✓ | -0.162 | -0.0789 |
| hAQP5 +100 ng hCAII | hAQP5 + 1 ng hCA II | -0.0224 | 2.39 | 0.692 | - | -0.0640 | 0.0192 |
| hAQP5 +100 ng hCA II | H<sub>2</sub>O + 10 ng hCA II | -0.121 | 13.0 | $3.68 \times 10^{-8}$ | ✓ | -0.163 | -0.0798 |
| hAQP5 +100 ng hCA II | hAQP5 + 10 ng hCA II | -0.0232 | 2.49 | 0.650 | - | -0.0648 | 0.0184 |
| hAQP5 +100 ng hCA II | H<sub>2</sub>O + 100 ng hCA II | -0.121 | 12.9 | $3.70 \times 10^{-8}$ | ✓ | -0.163 | -0.0793 |

**Supplemental Table 3C.**
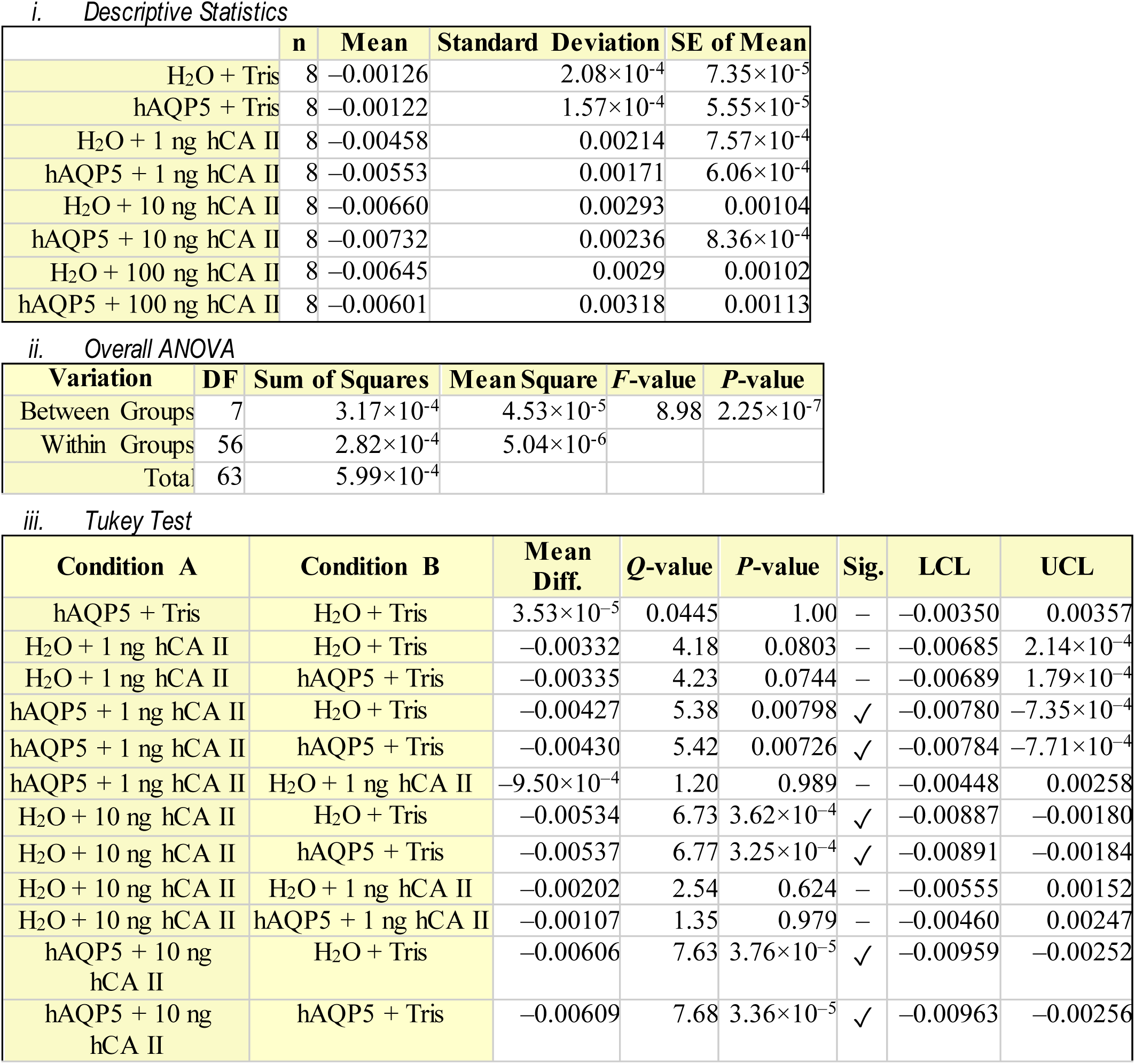
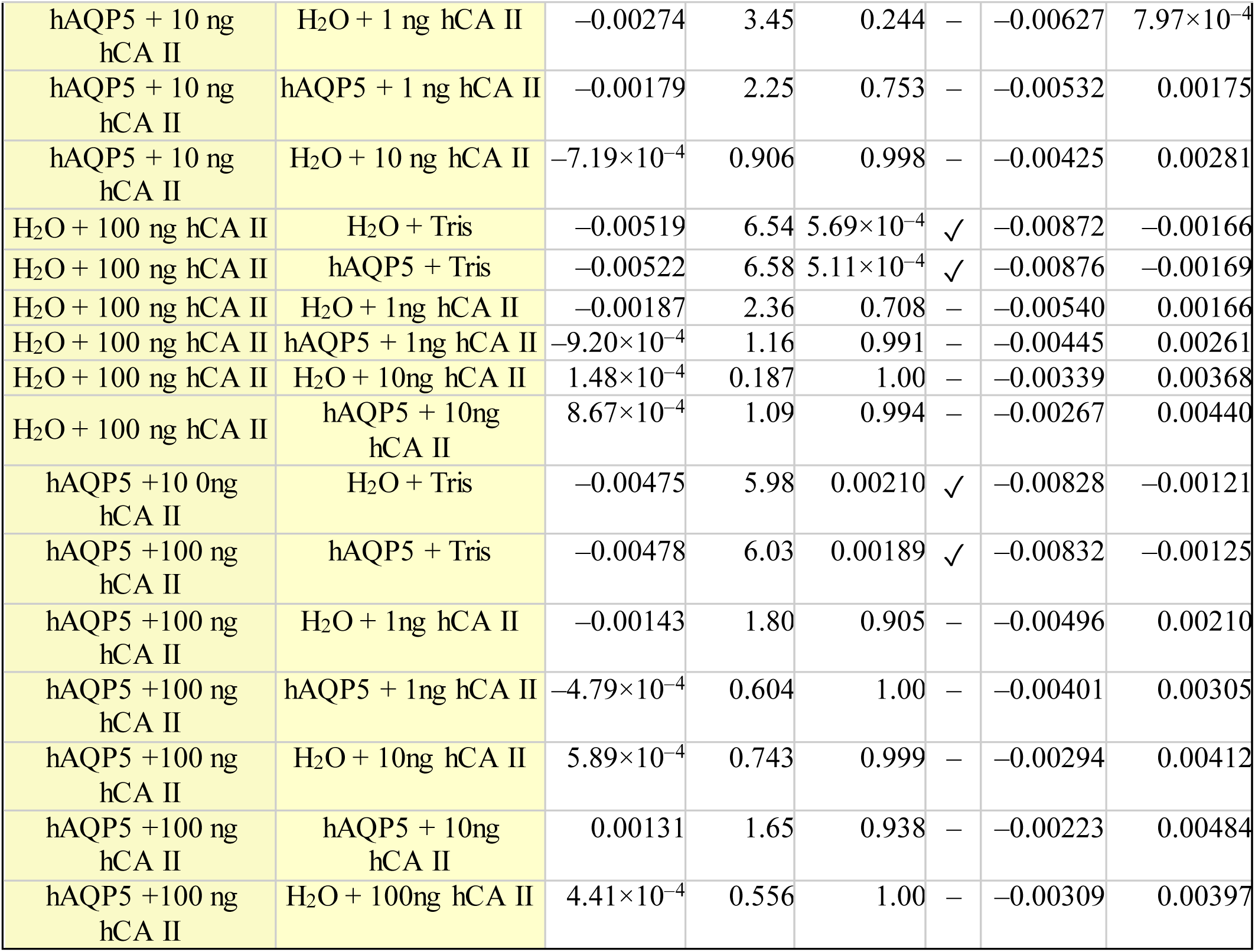
Statistics Summary for **Figure 3*C*, i**) Descriptive Statistics for data presented in Figure 3*C*, (dpH_i_/dt)_Max_ upon CO_2_ addition. ii) Overall one-way ANOVA results for data presented in Figure 3*C*. DF is degrees of freedom. iii) Tukey’s means comparison analysis for all data presented in Figure 3*C*. Threshold for significance α=0.05. Standard error of the mean = 0.0112. Mean difference for each comparison (Mean Diff.), lower confidence limit (LCL), upper confidence limit (UCL).

**Supplemental Table 3D.** Statistics Summary for **Figure 3*D***, **i**) Descriptive Statistics for data presented in Figure 3*D*, (dpH_i_/dt)_Max_ upon CO_2_ removal. ii) Overall one-way ANOVA results for data presented in Figure 3*D*. DF is degrees of freedom. iii) Tukey’s means comparison analysis for all data presented in Figure 3*D*. Threshold for significance α=0.05. Standard error of the mean = 6.27×10^−4^. Mean difference for each comparison (Mean Diff.), lower confidence limit (LCL), upper confidence limit (UCL).

|  | n | Mean | Standard Deviation | SE of Mean |
| --- | --- | --- | --- | --- |
| H<sub>2</sub>O + Tris | 8 | $8.35 \times 10^{-4}$ | $1.34 \times 10^{-4}$ | $4.74 \times 10^{-5}$ |
| hAQP5 + Tris | 8 | $8.07 \times 10^{-4}$ | $6.85 \times 10^{-5}$ | $2.42 \times 10^{-5}$ |
| H<sub>2</sub>O + 1 ng hCA II | 8 | 0.00263 | 0.00119 | $4.19 \times 10^{-4}$ |
| hAQP5 + 1 ng hCA II | 8 | 0.00306 | $8.73 \times 10^{-4}$ | $3.08 \times 10^{-4}$ |
| H<sub>2</sub>O + 10 ng hCA II | 8 | 0.00355 | 0.00121 | $4.29 \times 10^{-4}$ |
| hAQP5 + 10 ng hCA II | 8 | 0.00399 | 0.00139 | $4.92 \times 10^{-4}$ |
| H<sub>2</sub>O + 100 ng hCA II | 8 | 0.00373 | 0.00266 | $9.39 \times 10^{-4}$ |
| hAQP5 + 100 ng hCA II | 8 | 0.00297 | 0.00134 | $4.74 \times 10^{-4}$ |

ii. Overall ANOVA
| Variation | DF | Sum of Squares | Mean Square | F-value | P-value |
| --- | --- | --- | --- | --- | --- |
| Between Groups | 7 | $8.57 \times 10^{-5}$ | $1.22 \times 10^{-5}$ | 6.78 | $7.89 \times 10^{-6}$ |
| Within Groups | 56 | $1.01 \times 10^{-4}$ | $1.81 \times 10^{-6}$ | | |
| Total | 63 | $1.87 \times 10^{-4}$ | | | |

iii. Tukey Test
| Condition A | Condition B | Mean Diff. | Q-value | P-value | Sig. | LCL | UCL |
| --- | --- | --- | --- | --- | --- | --- | --- |
| hAQP5 + Tris | H<sub>2</sub>O + Tris | $-2.75 \times 10^{-5}$ | 0.0579 | 1.00 | – | $-0.00214$ | 0.00209 |
| H<sub>2</sub>O + 1 ng hCA II | H<sub>2</sub>O + Tris | 0.00179 | 3.77 | 0.154 | – | $-3.22 \times 10^{-4}$ | 0.00391 |
| H<sub>2</sub>O + 1 ng hCA II | hAQP5 + Tris | 0.00182 | 3.83 | 0.141 | – | $-2.95 \times 10^{-4}$ | 0.00394 |
| hAQP5 + 1 ng hCA II | H<sub>2</sub>O + Tris | 0.00222 | 4.68 | 0.0329 | ✓ | $1.07 \times 10^{-4}$ | 0.00434 |
| hAQP5 + 1 ng hCA II | hAQP5 + Tris | 0.00225 | 4.74 | 0.0295 | ✓ | $1.34 \times 10^{-4}$ | 0.00437 |
| hAQP5 + 1 ng hCA II | H<sub>2</sub>O + 1 ng hCA II | $4.29 \times 10^{-4}$ | 0.903 | 0.998 | – | $-0.00169$ | 0.00255 |
| H<sub>2</sub>O + 10 ng hCA II | H<sub>2</sub>O + Tris | 0.00272 | 5.72 | 0.00381 | ✓ | $6.02 \times 10^{-4}$ | 0.00483 |
| H<sub>2</sub>O + 10 ng hCA II | hAQP5 + Tris | 0.00275 | 5.78 | 0.00335 | ✓ | $6.29 \times 10^{-4}$ | 0.00486 |
| H<sub>2</sub>O + 10 ng hCA II | H<sub>2</sub>O + 1 ng hCA II | $9.24 \times 10^{-4}$ | 1.94 | 0.865 | – | $-0.00119$ | 0.00304 |
| H<sub>2</sub>O + 10 ng hCA II | hAQP5 + 1 ng hCA II | $4.95 \times 10^{-4}$ | 1.04 | 0.995 | – | $-0.00162$ | 0.00261 |
| hAQP5 + 10 ng hCA II | H<sub>2</sub>O + Tris | 0.00315 | 6.63 | $4.57 \times 10^{-4}$ | ✓ | 0.00103 | 0.00527 |
| hAQP5 + 10 ng hCA II | hAQP5 + Tris | 0.00318 | 6.69 | $3.97 \times 10^{-4}$ | ✓ | 0.00106 | 0.00529 |
| hAQP5 + 10 ng hCA II | H<sub>2</sub>O + 1 ng hCA II | 0.00136 | 2.85 | 0.479 | – | –7.59×10<sup>–4</sup> | 0.00347 |
| hAQP5 + 10 ng hCA II | hAQP5 + 1 ng hCA II | 9.28×10<sup>–4</sup> | 1.95 | 0.862 | – | –0.00119 | 0.00304 |
| hAQP5 + 10 ng hCA II | H<sub>2</sub>O + 10 ng hCA II | 4.33×10<sup>–4</sup> | 0.911 | 0.998 | – | –0.00168 | 0.00255 |
| H<sub>2</sub>O + 100ng hCA II | H<sub>2</sub>O + Tris | 0.00290 | 6.10 | 0.00160 | ✓ | 7.84×10<sup>–4</sup> | 0.00502 |
| H<sub>2</sub>O + 100ng hCA II | hAQP5 + Tris | 0.00293 | 6.16 | 0.00140 | ✓ | 8.11×10<sup>–4</sup> | 0.00504 |
| H<sub>2</sub>O + 100ng hCA II | H<sub>2</sub>O + 1 ng hCA II | 0.00111 | 2.33 | 0.721 | – | –0.00101 | 0.00322 |
| H<sub>2</sub>O + 100ng hCA II | hAQP5 + 1 ng hCA II | 6.77×10<sup>–4</sup> | 1.42 | 0.972 | – | –0.00144 | 0.00279 |
| H<sub>2</sub>O + 100ng hCA II | H<sub>2</sub>O + 10 ng hCA II | 1.82×10<sup>–4</sup> | 0.383 | 1.00 | – | –0.00193 | 0.00230 |
| H<sub>2</sub>O + 100ng hCA II | hAQP5 + 10 ng hCA II | –2.51×10<sup>–4</sup> | 0.528 | 1.00 | – | –0.00237 | 0.00187 |
| hAQP5 +100ng hCA II | H<sub>2</sub>O + Tris | 0.00214 | 4.50 | 0.0460 | ✓ | 2.19×10<sup>–5</sup> | 0.00425 |
| hAQP5 +100ng hCA II | hAQP5 + Tris | 0.00217 | 4.56 | 0.0413 | ✓ | 4.94×10<sup>–5</sup> | 0.00428 |
| hAQP5 +100ng hCA II | H<sub>2</sub>O + 1 ng hCA II | 3.44×10<sup>–4</sup> | 0.724 | 1.00 | – | –0.00177 | 0.00246 |
| hAQP5 +100ng hCA II | hAQP5 + 1 ng hCA II | –8.49×10<sup>–5</sup> | 0.179 | 1.00 | – | –0.00220 | 0.00203 |
| hAQP5 +100ng hCA II | H<sub>2</sub>O + 10 ng hCA II | –5.80×10<sup>–4</sup> | 1.22 | 0.988 | – | –0.00270 | 0.00154 |
| hAQP5 +100ng hCA II | hAQP5 + 10 ng hCA II | –0.00101 | 2.13 | 0.801 | – | –0.00313 | 0.00110 |
| hAQP5 +100ng hCA II | H<sub>2</sub>O + 100 ng hCA II | –7.62×10<sup>–4</sup> | 1.60 | 0.947 | – | –0.00288 | 0.00135 |

**Supplemental Table 5A.**
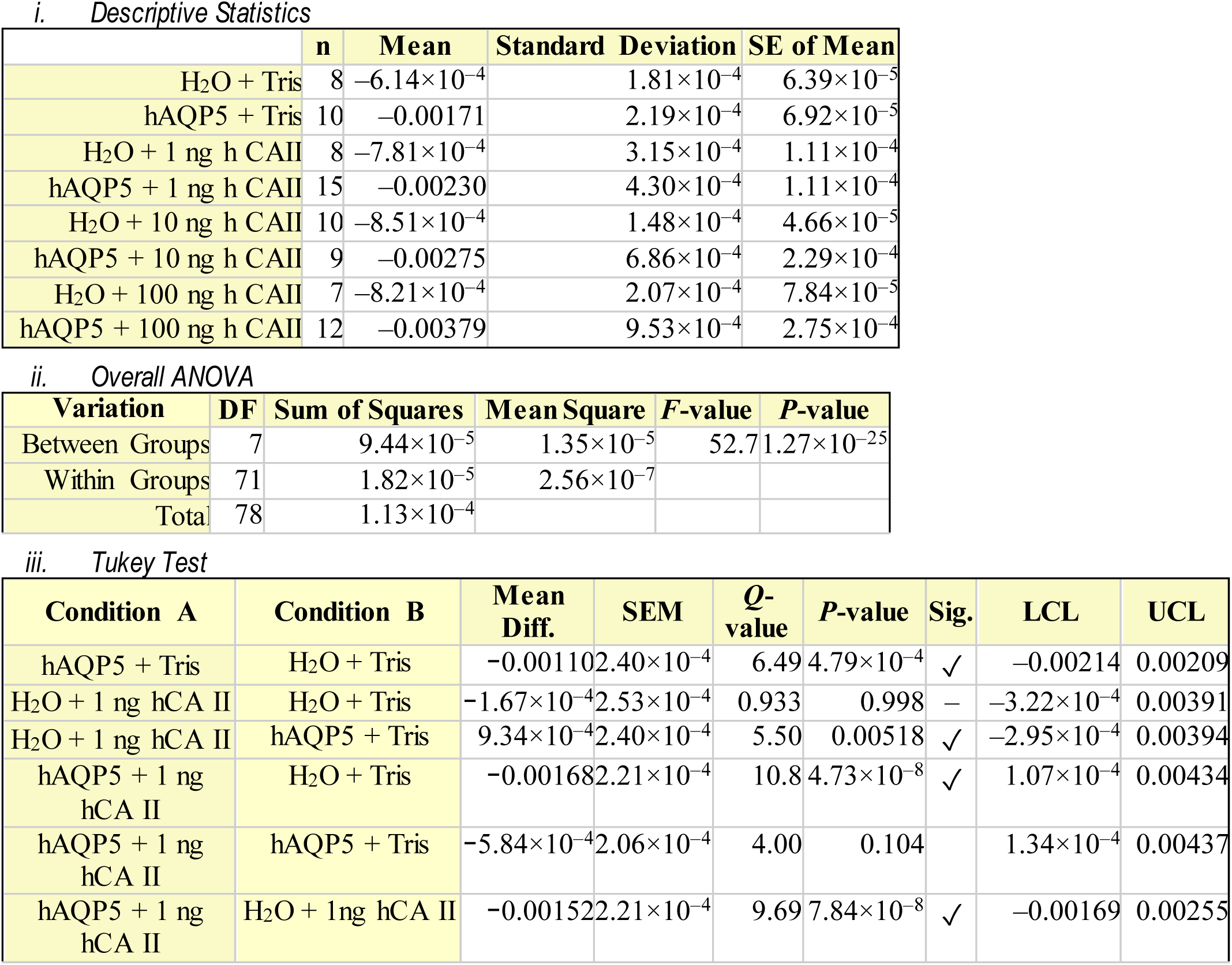
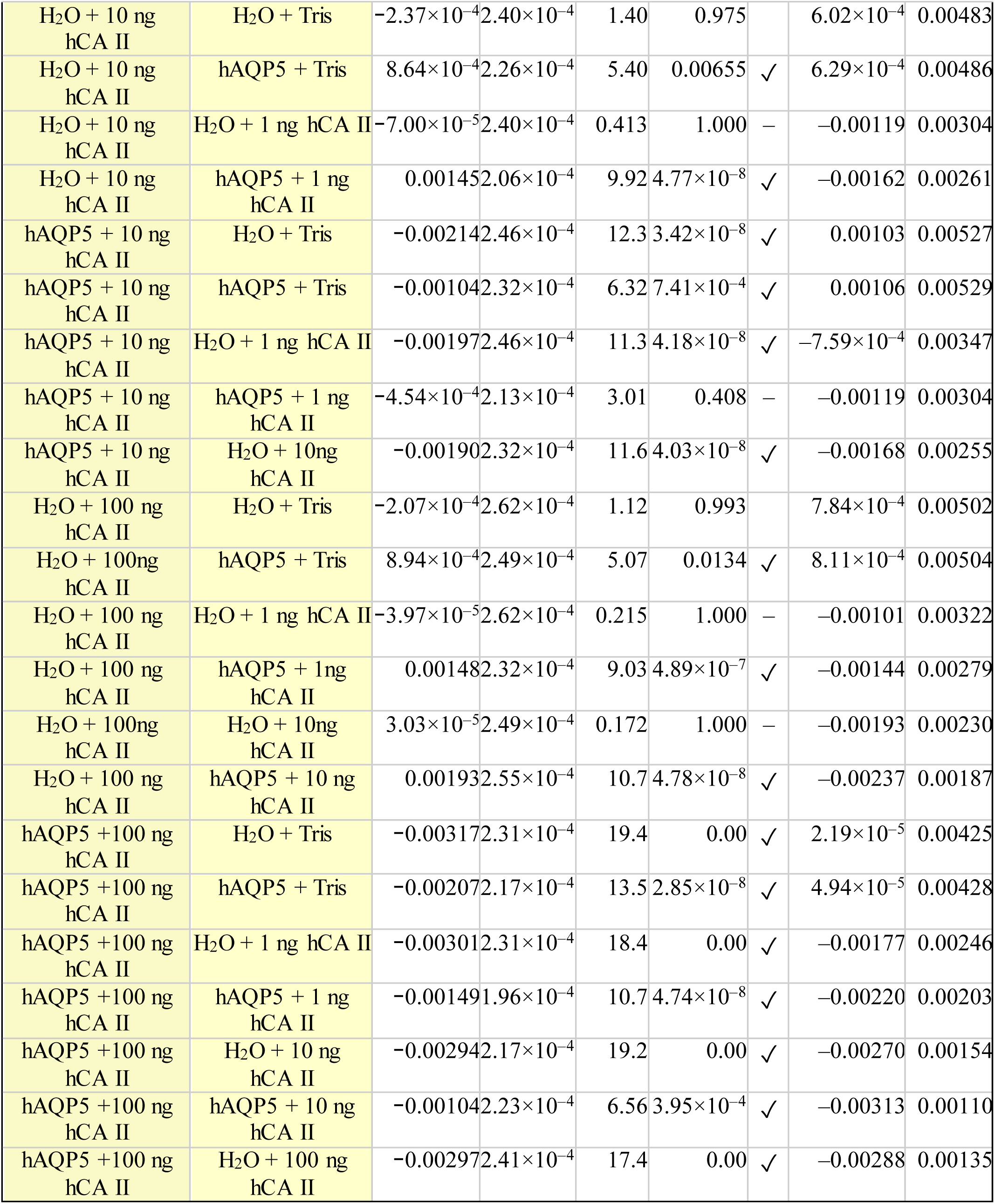
Statistics Summary for **Figure 5*A***, i) Descriptive Statistics for data presented in Figure 5*A*, (dpH_S_/dt)_Max_ upon CO_2_ addition. ii) Overall one-way ANOVA results for data presented in Figure 5*A*. DF is degrees of freedom. iii) Tukey’s means comparison analysis for all data presented in Figure 5*A*. Threshold for significance α=0.05. Standard error of the mean (SEM). Mean difference for each comparison (Mean Diff.), lower confidence limit (LCL), upper confidence limit (UCL). Where the displayed *P*-values is 0.00, this indicates a values less than <2.22×10**^−^**^308^. ^13^

|  | n | Mean | Standard Deviation | SE of Mean |
| --- | --- | --- | --- | --- |
| H<sub>2</sub>O + Tris | 8 | $-6.14 \times 10^{-4}$ | $1.81 \times 10^{-4}$ | $6.39 \times 10^{-5}$ |
| hAQP5 + Tris | 10 | -0.00171 | $2.19 \times 10^{-4}$ | $6.92 \times 10^{-5}$ |
| H<sub>2</sub>O + 1 ng h CA II | 8 | $-7.81 \times 10^{-4}$ | $3.15 \times 10^{-4}$ | $1.11 \times 10^{-4}$ |
| hAQP5 + 1 ng h CA II | 15 | -0.00230 | $4.30 \times 10^{-4}$ | $1.11 \times 10^{-4}$ |
| H<sub>2</sub>O + 10 ng h CA II | 10 | $-8.51 \times 10^{-4}$ | $1.48 \times 10^{-4}$ | $4.66 \times 10^{-5}$ |
| hAQP5 + 10 ng h CA II | 9 | -0.00275 | $6.86 \times 10^{-4}$ | $2.29 \times 10^{-4}$ |
| H<sub>2</sub>O + 100 ng h CA II | 7 | $-8.21 \times 10^{-4}$ | $2.07 \times 10^{-4}$ | $7.84 \times 10^{-5}$ |
| hAQP5 + 100 ng h CA II | 12 | -0.00379 | $9.53 \times 10^{-4}$ | $2.75 \times 10^{-4}$ |

ii. Overall ANOVA
| Variation | DF | Sum of Squares | Mean Square | F-value | P-value |
| --- | --- | --- | --- | --- | --- |
| Between Groups | 7 | $9.44 \times 10^{-5}$ | $1.35 \times 10^{-5}$ | 52.7 | $1.27 \times 10^{-25}$ |
| Within Groups | 71 | $1.82 \times 10^{-5}$ | $2.56 \times 10^{-7}$ | | |
| Total | 78 | $1.13 \times 10^{-4}$ | | | |

iii. Tukey Test
| Condition A | Condition B | Mean Diff. | SEM | Q-value | P-value | Sig. | LCL | UCL |
| --- | --- | --- | --- | --- | --- | --- | --- | --- |
| hAQP5 + Tris | H<sub>2</sub>O + Tris | -0.00110 | $2.40 \times 10^{-4}$ | 6.49 | $4.79 \times 10^{-4}$ | ✓ | -0.00214 | 0.00209 |
| H<sub>2</sub>O + 1 ng hCA II | H<sub>2</sub>O + Tris | $-1.67 \times 10^{-4}$ | $2.53 \times 10^{-4}$ | 0.933 | 0.998 | — | $-3.22 \times 10^{-4}$ | 0.00391 |
| H<sub>2</sub>O + 1 ng hCA II | hAQP5 + Tris | $9.34 \times 10^{-4}$ | $2.40 \times 10^{-4}$ | 5.50 | 0.00518 | ✓ | $-2.95 \times 10^{-4}$ | 0.00394 |
| hAQP5 + 1 ng hCA II | H<sub>2</sub>O + Tris | -0.00168 | $2.21 \times 10^{-4}$ | 10.8 | $4.73 \times 10^{-8}$ | ✓ | $1.07 \times 10^{-4}$ | 0.00434 |
| hAQP5 + 1 ng hCA II | hAQP5 + Tris | $-5.84 \times 10^{-4}$ | $2.06 \times 10^{-4}$ | 4.00 | 0.104 | | $1.34 \times 10^{-4}$ | 0.00437 |
| hAQP5 + 1 ng hCA II | H<sub>2</sub>O + 1 ng hCA II | -0.00152 | $2.21 \times 10^{-4}$ | 9.69 | $7.84 \times 10^{-8}$ | ✓ | -0.00169 | 0.00255 |

<sup>13</sup> $2.22 \times 10^{-308}$ is the smallest possible value for double type data that a 64-bit system is able to distinguish

|  |  |  |  |  |  |  |  |  |
| --- | --- | --- | --- | --- | --- | --- | --- | --- |
| H<sub>2</sub>O + 10 ng hCA II | H<sub>2</sub>O + Tris | -2.37×10<sup>-4</sup> | 2.40×10<sup>-4</sup> | 1.40 | 0.975 |  | 6.02×10<sup>-4</sup> | 0.00483 |
| H<sub>2</sub>O + 10 ng hCA II | hAQP5 + Tris | 8.64×10<sup>-4</sup> | 2.26×10<sup>-4</sup> | 5.40 | 0.00655 | ✓ | 6.29×10<sup>-4</sup> | 0.00486 |
| H<sub>2</sub>O + 10 ng hCA II | H<sub>2</sub>O + 1 ng hCA II | -7.00×10<sup>-5</sup> | 2.40×10<sup>-4</sup> | 0.413 | 1.000 | – | –0.00119 | 0.00304 |
| H<sub>2</sub>O + 10 ng hCA II | hAQP5 + 1 ng hCA II | 0.00145 | 2.06×10<sup>-4</sup> | 9.92 | 4.77×10<sup>-8</sup> | ✓ | –0.00162 | 0.00261 |
| hAQP5 + 10 ng hCA II | H<sub>2</sub>O + Tris | –0.00214 | 2.46×10<sup>-4</sup> | 12.3 | 3.42×10<sup>-8</sup> | ✓ | 0.00103 | 0.00527 |
| hAQP5 + 10 ng hCA II | hAQP5 + Tris | –0.00104 | 2.32×10<sup>-4</sup> | 6.32 | 7.41×10<sup>-4</sup> | ✓ | 0.00106 | 0.00529 |
| hAQP5 + 10 ng hCA II | H<sub>2</sub>O + 1 ng hCA II | –0.00197 | 2.46×10<sup>-4</sup> | 11.3 | 4.18×10<sup>-8</sup> | ✓ | –7.59×10<sup>-4</sup> | 0.00347 |
| hAQP5 + 10 ng hCA II | hAQP5 + 1 ng hCA II | –4.54×10<sup>-4</sup> | 2.13×10<sup>-4</sup> | 3.01 | 0.408 | – | –0.00119 | 0.00304 |
| hAQP5 + 10 ng hCA II | H<sub>2</sub>O + 10ng hCA II | –0.00190 | 2.32×10<sup>-4</sup> | 11.6 | 4.03×10<sup>-8</sup> | ✓ | –0.00168 | 0.00255 |
| H<sub>2</sub>O + 100 ng hCA II | H<sub>2</sub>O + Tris | –2.07×10<sup>-4</sup> | 2.62×10<sup>-4</sup> | 1.12 | 0.993 |  | 7.84×10<sup>-4</sup> | 0.00502 |
| H<sub>2</sub>O + 100ng hCA II | hAQP5 + Tris | 8.94×10<sup>-4</sup> | 2.49×10<sup>-4</sup> | 5.07 | 0.0134 | ✓ | 8.11×10<sup>-4</sup> | 0.00504 |
| H<sub>2</sub>O + 100 ng hCA II | H<sub>2</sub>O + 1 ng hCA II | –3.97×10<sup>-5</sup> | 2.62×10<sup>-4</sup> | 0.215 | 1.000 | – | –0.00101 | 0.00322 |
| H<sub>2</sub>O + 100 ng hCA II | hAQP5 + 1ng hCA II | 0.00148 | 2.32×10<sup>-4</sup> | 9.03 | 4.89×10<sup>-7</sup> | ✓ | –0.00144 | 0.00279 |
| H<sub>2</sub>O + 100ng hCA II | H<sub>2</sub>O + 10ng hCA II | 3.03×10<sup>-5</sup> | 2.49×10<sup>-4</sup> | 0.172 | 1.000 | – | –0.00193 | 0.00230 |
| H<sub>2</sub>O + 100 ng hCA II | hAQP5 + 10 ng hCA II | 0.00193 | 2.55×10<sup>-4</sup> | 10.7 | 4.78×10<sup>-8</sup> | ✓ | –0.00237 | 0.00187 |
| hAQP5 + 100 ng hCA II | H<sub>2</sub>O + Tris | –0.00317 | 2.31×10<sup>-4</sup> | 19.4 | 0.00 | ✓ | 2.19×10<sup>-5</sup> | 0.00425 |
| hAQP5 + 100 ng hCA II | hAQP5 + Tris | –0.00207 | 2.17×10<sup>-4</sup> | 13.5 | 2.85×10<sup>-8</sup> | ✓ | 4.94×10<sup>-5</sup> | 0.00428 |
| hAQP5 + 100 ng hCA II | H<sub>2</sub>O + 1 ng hCA II | –0.00301 | 2.31×10<sup>-4</sup> | 18.4 | 0.00 | ✓ | –0.00177 | 0.00246 |
| hAQP5 + 100 ng hCA II | hAQP5 + 1 ng hCA II | –0.00149 | 1.96×10<sup>-4</sup> | 10.7 | 4.74×10<sup>-8</sup> | ✓ | –0.00220 | 0.00203 |
| hAQP5 + 100 ng hCA II | H<sub>2</sub>O + 10 ng hCA II | –0.00294 | 2.17×10<sup>-4</sup> | 19.2 | 0.00 | ✓ | –0.00270 | 0.00154 |
| hAQP5 + 100 ng hCA II | hAQP5 + 10 ng hCA II | –0.00104 | 2.23×10<sup>-4</sup> | 6.56 | 3.95×10<sup>-4</sup> | ✓ | –0.00313 | 0.00110 |
| hAQP5 + 100 ng hCA II | H<sub>2</sub>O + 100 ng hCA II | –0.00297 | 2.41×10<sup>-4</sup> | 17.4 | 0.00 | ✓ | –0.00288 | 0.00135 |

**Supplemental Table 5B.**
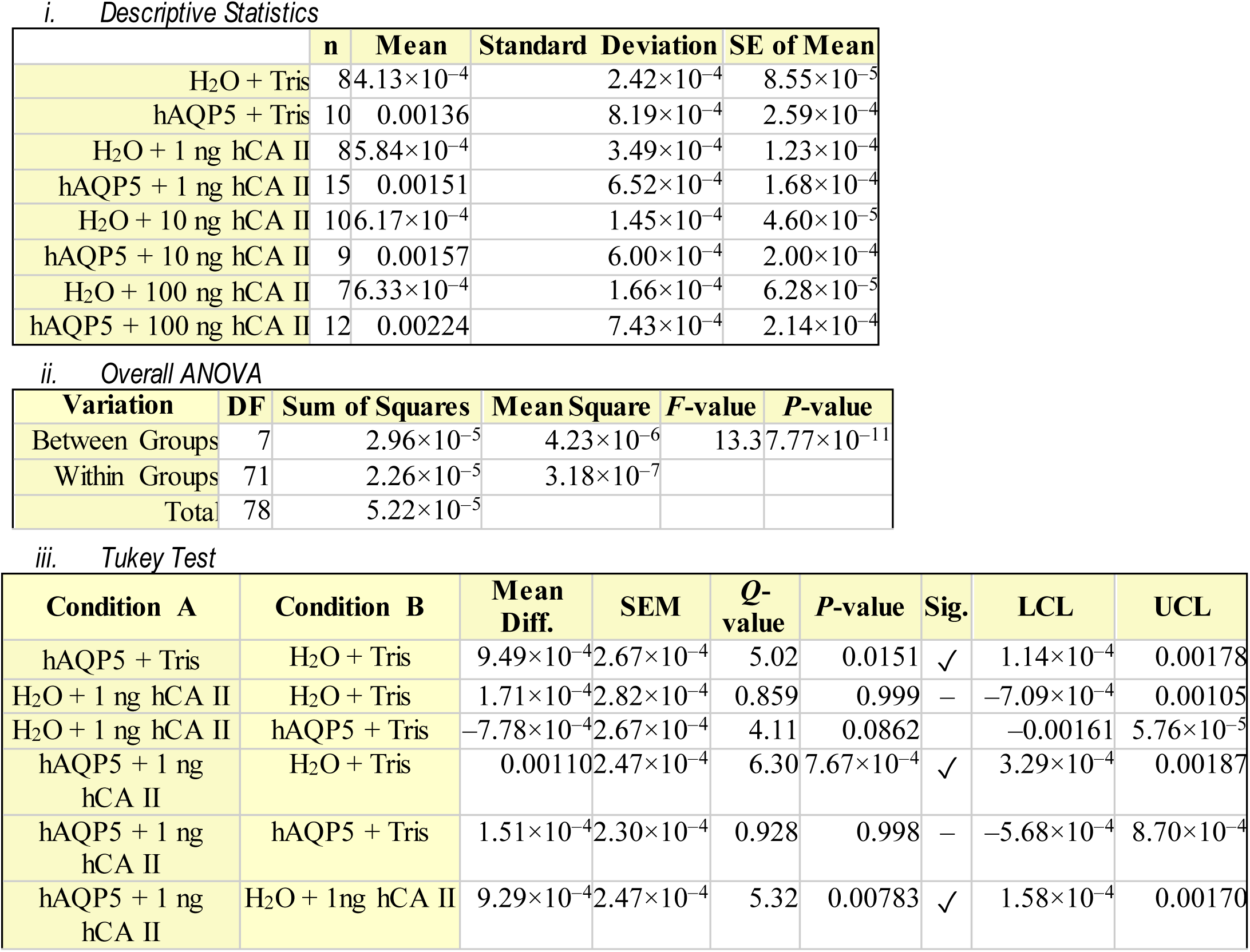
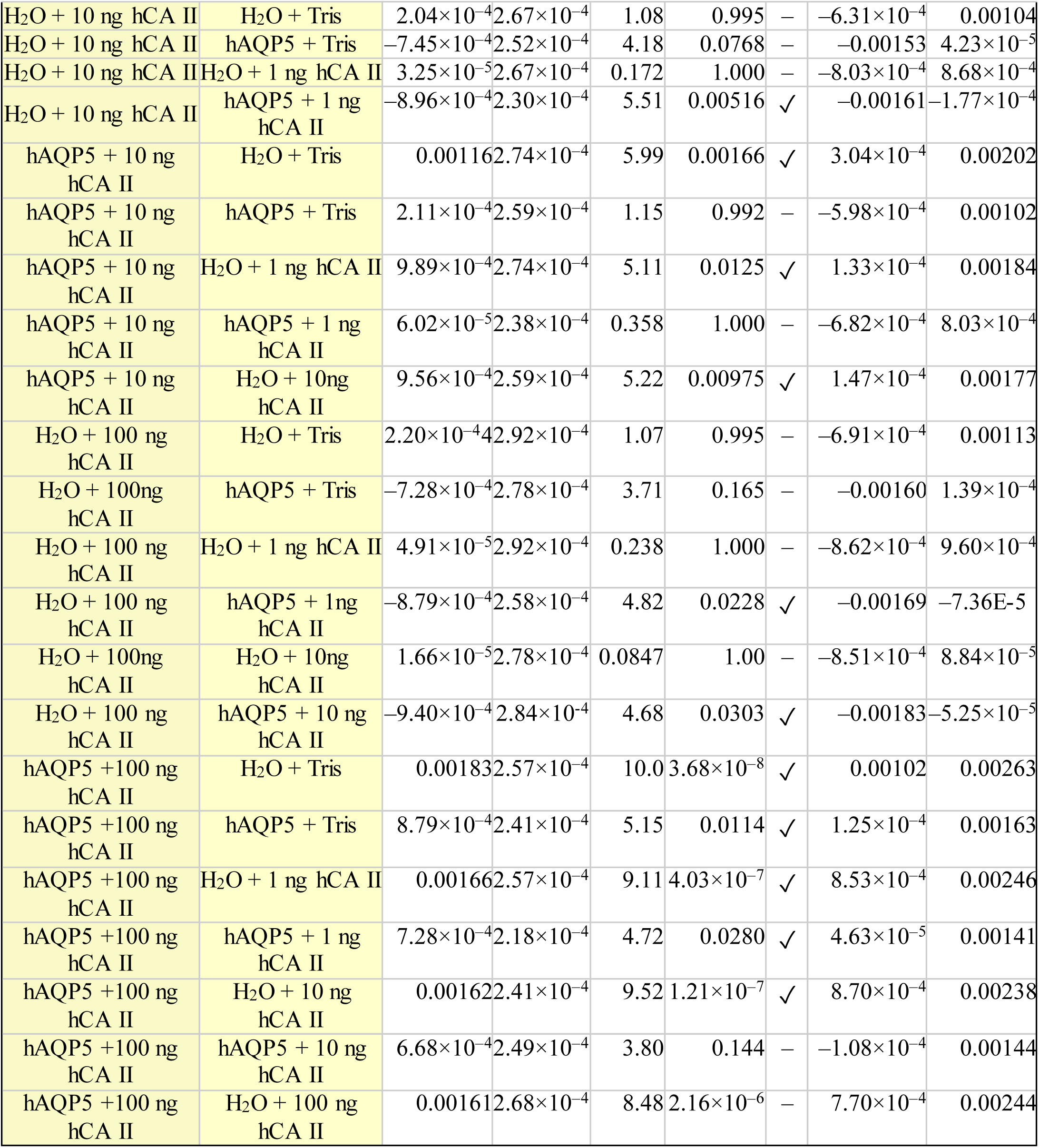
Statistics Summary for **Figure 5*B***, **i**) Descriptive Statistics for data presented in Figure 5*B*, (dpH_S_/dt)_Max_ upon CO_2_ removal. ii) Overall one-way ANOVA results for data presented in Figure 5*A*. DF is degrees of freedom. iii) Tukey’s means comparison analysis for all data presented in Figure 5*B*. Threshold for significance α=0.05. Standard error of the mean (SEM). Mean difference for each comparison (Mean Diff.), lower confidence limit (LCL), upper confidence limit (UCL). Where the displayed *P*-values is 0.00, this indicates a values less than <2.22×10**^−^**^308^. ^14^

**Supplemental Table 6A.** Statistics Summary for **Figure 6*A***, **i**) Descriptive Statistics for data presented in Figure 6A, Initial pH_i_. ii) Overall one-way ANOVA results for data presented in Figure 6*A*. DF is degrees of freedom. iii) Tukey’s means comparison analysis for all data presented in Figure 6*A*. Threshold for significance α=0.05. Standard error of the mean = 0.0262. Mean difference for each comparison (Mean Diff.), lower confidence limit (LCL), upper confidence limit (UCL).

|  | n | Mean | Standard Deviation | SE of Mean |
| --- | --- | --- | --- | --- |
| H<sub>2</sub>O + Tris | 8 | 7.27 | 0.0486 | 0.0172 |
| hAQP5 + Tris | 8 | 7.31 | 0.0512 | 0.0181 |
| H<sub>2</sub>O + 1 ng hCA II | 8 | 7.30 | 0.0614 | 0.0217 |
| hAQP5 + 1 ng hCA II | 8 | 7.32 | 0.0512 | 0.0181 |
| H<sub>2</sub>O + 10 ng hCA II | 8 | 7.26 | 0.0452 | 0.0160 |
| hAQP5 + 10 ng hCA II | 8 | 7.31 | 0.0352 | 0.0124 |
| H<sub>2</sub>O + 100 ng hCA II | 8 | 7.26 | 0.0614 | 0.0217 |
| hAQP5 + 100 ng hCA II | 8 | 7.29 | 0.0593 | 0.0210 |

*ii. Overall ANOVA Table*
| Variation | DF | Sum of Squares | Mean Square | F-value | P-value |
| --- | --- | --- | --- | --- | --- |
| Between Groups | 7 | 0.0324 | 0.00462 | 1.68 | 0.131 |
| Within Groups | 56 | 0.154 | 0.00274 |  |  |
| Total | 63 | 0.186 |  |  |  |

*iii. Tukey Test*
| Condition A | Condition B | Mean Diff. | Q-value | P-value | Sig. | LCL | UCL |
| --- | --- | --- | --- | --- | --- | --- | --- |
| hAQP5 + Tris | H<sub>2</sub>O + Tris | 0.0440 | 2.37 | 0.701 | – | –0.0385 | 0.126 |
| H<sub>2</sub>O + 1 ng hCA II | H<sub>2</sub>O + Tris | 0.0264 | 1.43 | 0.971 | – | –0.0561 | 0.109 |
| H<sub>2</sub>O + 1 ng hCA II | hAQP5 + Tris | –0.0176 | 0.948 | 0.997 | – | –0.100 | 0.0649 |
| hAQP5 + 1 ng hCA II | H<sub>2</sub>O + Tris | 0.0515 | 2.78 | 0.512 | – | –0.0309 | 0.134 |
| hAQP5 + 1 ng hCA II | hAQP5 + Tris | 0.00759 | 0.410 | 1.000 | – | –0.0749 | 0.0900 |
| hAQP5 + 1 ng hCA II | H<sub>2</sub>O + 1 ng hCA II | 0.0252 | 1.36 | 0.978 | – | –0.0573 | 0.108 |
| H<sub>2</sub>O + 10 ng hCA II | H<sub>2</sub>O + Tris | –0.00884 | 0.477 | 1.000 | – | –0.0913 | 0.0736 |
| H<sub>2</sub>O + 10 ng hCA II | hAQP5 + Tris | –0.0528 | 2.85 | 0.481 | – | –0.135 | 0.0297 |
| H<sub>2</sub>O + 10 ng hCA II | H<sub>2</sub>O + 1 ng hCA II | –0.0352 | 1.90 | 0.877 | – | –0.118 | 0.0472 |
| H<sub>2</sub>O + 10 ng hCA II | hAQP5 + 1 ng hCA II | –0.0604 | 3.26 | 0.309 | – | –0.143 | 0.0221 |
| hAQP5 + 10 ng hCA II | H<sub>2</sub>O + Tris | 0.0439 | 2.37 | 0.703 | – | –0.0386 | 0.126 |
| hAQP5 + 10 ng hCA II | hAQP5 + Tris | –1.08×10<sup>–4</sup> | 0.00586 | 1.00 | – | –0.0826 | 0.0824 |
| hAQP5 + 10 ng hCA II | H<sub>2</sub>O + 1 ng hCA II | 0.0175 | 0.943 | 0.998 | – | –0.0650 | 0.0999 |
| hAQP5 + 10 ng hCA II | hAQP5 + 1 ng hCA II | –0.00770 | 0.416 | 1.000 | – | –0.0902 | 0.0748 |
| hAQP5 + 10 ng hCA II | H<sub>2</sub>O + 10ng hCA II | 0.0527 | 2.84 | 0.484 | – | –0.0298 | 0.135 |
| H<sub>2</sub>O + 100 ng hCA II | H<sub>2</sub>O + Tris | –0.00531 | 0.287 | 1.000 | – | –0.0878 | 0.0772 |
| H<sub>2</sub>O + 100ng hCA II | hAQP5 + Tris | –0.0493 | 2.66 | 0.569 | – | –0.132 | 0.0332 |
| H<sub>2</sub>O + 100 ng hCA II | H<sub>2</sub>O + 1 ng hCA II | –0.0317 | 1.71 | 0.925 | – | –0.114 | 0.0508 |
| H<sub>2</sub>O + 100 ng hCA II | hAQP5 + 1 ng hCA II | –0.0569 | 3.07 | 0.385 | – | –0.139 | 0.0256 |
| H<sub>2</sub>O + 100ng hCA II | H<sub>2</sub>O + 10 ng hCA II | 0.00353 | 0.191 | 1.000 | – | –0.0789 | 0.0860 |
| H<sub>2</sub>O + 100 ng hCA II | hAQP5 + 10 ng hCA II | –0.0492 | 2.65 | 0.572 | – | –0.132 | 0.0333 |
| hAQP5 +100 ng hCA II | H<sub>2</sub>O + Tris | 0.0177 | 0.957 | 0.997 | – | –0.0647 | 0.100 |
| hAQP5 +100 ng hCA II | hAQP5 + Tris | –0.0262 | 1.42 | 0.972 | – | –0.109 | 0.0562 |
| hAQP5 +100 ng hCA II | H<sub>2</sub>O + 1 ng hCA II | –0.00866 | 0.468 | 1.000 | – | –0.0911 | 0.0738 |
| hAQP5 +100 ng hCA II | hAQP5 + 1 ng hCA II | –0.0338 | 1.83 | 0.898 | – | –0.116 | 0.0486 |
| hAQP5 +100 ng hCA II | H<sub>2</sub>O + 10 ng hCA II | 0.0266 | 1.43 | 0.970 | – | –0.0559 | 0.109 |
| hAQP5 +100 ng hCA II | hAQP5 + 10 ng hCA II | –0.0261 | 1.41 | 0.973 | – | –0.109 | 0.0563 |
| hAQP5 +100 ng hCA II | H<sub>2</sub>O + 100 ng hCA II | 0.0230 | 1.24 | 0.987 | – | –0.0594 | 0.105 |

**Supplemental Table 6B.** Statistics Summary for **Figure 6B, i**) Descriptive Statistics for data presented in Figure 6B, ΔpH_i_. ii) Overall one-way ANOVA results for data presented in Figure 6*B*. DF is degrees of freedom. iii) Tukey’s means comparison analysis for all data presented in Figure 6*B*. Threshold for significance α=0.05. Standard error of the mean = 0.0137. Mean difference for each comparison (Mean Diff.), lower confidence limit (LCL), upper confidence limit (UCL).

|  | n | Mean | Standard Deviation | SE of Mean |
| --- | --- | --- | --- | --- |
| H<sub>2</sub>O + Tris | 8 | 0.251 | 0.0220 | 0.00778 |
| hAQP5 + Tris | 8 | 0.257 | 0.0253 | 0.00894 |
| H<sub>2</sub>O + 1 ng hCA II | 8 | 0.252 | 0.0296 | 0.0105 |
| hAQP5 + 1 ng hCA II | 8 | 0.274 | 0.0234 | 0.00828 |
| H<sub>2</sub>O + 10 ng hCA II | 8 | 0.264 | 0.0209 | 0.00740 |
| hAQP5 + 10 ng hCA II | 8 | 0.292 | 0.0238 | 0.00841 |
| H<sub>2</sub>O + 100 ng hCA II | 8 | 0.252 | 0.0254 | 0.00898 |
| hAQP5 + 100 ng hCA II | 8 | 0.278 | 0.0422 | 0.0149 |

*ii. Overall ANOVA Table*
| Variation | DF | Sum of Squares | Mean Square | F-value | P-value |
| --- | --- | --- | --- | --- | --- |
| Between Groups | 7 | 0.0125 | 0.00178 | 2.39 | 0.0329 |
| Within Groups | 56 | 0.0419 | $7.47 \times 10^{-4}$ | | |
| Total | 63 | 0.0543 |  |  |  |

*iii. Tukey Test*
| Condition A | Condition B | Mean Diff. | Q-value | P-value | Sig. | LCL | UCL |
| --- | --- | --- | --- | --- | --- | --- | --- |
| hAQP5 + Tris | H<sub>2</sub>O + Tris | 0.00615 | 0.636 | 1.000 | – | –0.0369 | 0.0492 |
| H<sub>2</sub>O + 1 ng hCA II | H<sub>2</sub>O + Tris | 0.00173 | 0.179 | 1.000 | – | –0.0413 | 0.0448 |
| H<sub>2</sub>O + 1 ng hCA II | hAQP5 + Tris | –0.00442 | 0.457 | 1.000 | – | –0.0475 | 0.0386 |
| hAQP5 + 1 ng hCA II | H<sub>2</sub>O + Tris | 0.0234 | 2.42 | 0.679 | – | –0.0196 | 0.0664 |
| hAQP5 + 1 ng hCA II | hAQP5 + Tris | 0.0173 | 1.79 | 0.909 | – | –0.0258 | 0.0603 |
| hAQP5 + 1 ng hCA II | H<sub>2</sub>O + 1 ng hCA II | 0.0217 | 2.24 | 0.757 | – | –0.0214 | 0.0647 |
| H<sub>2</sub>O + 10 ng hCA II | H<sub>2</sub>O + Tris | 0.0129 | 1.34 | 0.980 | – | –0.0301 | 0.0560 |
| H<sub>2</sub>O + 10 ng hCA II | hAQP5 + Tris | 0.00677 | 0.700 | 1.000 | – | –0.0363 | 0.0498 |
| H<sub>2</sub>O + 10 ng hCA II | H<sub>2</sub>O + 1 ng hCA II | 0.0112 | 1.16 | 0.991 | – | –0.0319 | 0.0542 |
| H<sub>2</sub>O + 10 ng hCA II | hAQP5 + 1 ng hCA II | –0.0105 | 1.09 | 0.994 | – | –0.0535 | 0.0325 |
| hAQP5 + 10 ng hCA II | H<sub>2</sub>O + Tris | 0.0409 | 4.23 | 0.0737 | – | –0.00212 | 0.0840 |
| hAQP5 + 10 ng hCA II | hAQP5 + Tris | 0.0348 | 3.60 | 0.199 | – | –0.00828 | 0.0778 |
| hAQP5 + 10 ng hCA II | H<sub>2</sub>O + 1 ng hCA II | 0.0392 | 4.05 | 0.0995 | – | –0.00386 | 0.0822 |
| hAQP5 + 10 ng hCA II | hAQP5 + 1 ng hCA II | 0.0175 | 1.81 | 0.902 | – | –0.0255 | 0.0605 |
| hAQP5 + 10 ng hCA II | H<sub>2</sub>O + 10ng hCA II | 0.0280 | 2.90 | 0.460 | – | –0.0150 | 0.0710 |
| H<sub>2</sub>O + 100 ng hCA II | H<sub>2</sub>O + Tris | 0.00154 | 0.160 | 1.000 | – | –0.0415 | 0.0446 |
| H<sub>2</sub>O + 100ng hCA II | hAQP5 + Tris | –0.00461 | 0.477 | 1.000 | – | –0.0476 | 0.0384 |
| H<sub>2</sub>O + 100 ng hCA II | H<sub>2</sub>O + 1 ng hCA II | $-1.90 \times 10^{-4}$ | 0.0197 | 1.00 | – | –0.0432 | 0.0428 |
| H<sub>2</sub>O + 100 ng hCA II | hAQP5 + 1ng hCA II | –0.0219 | 2.26 | 0.748 | – | –0.0649 | 0.0212 |
| H<sub>2</sub>O + 100ng hCA II | H<sub>2</sub>O + 10ng hCA II | –0.0114 | 1.18 | 0.990 | – | –0.0544 | 0.0317 |
| H<sub>2</sub>O + 100 ng hCA II | hAQP5 + 10 ng hCA II | –0.0394 | 4.07 | 0.0963 | – | –0.0824 | 0.00367 |
| hAQP5 +100 ng hCA II | H<sub>2</sub>O + Tris | 0.0276 | 2.85 | 0.480 | – | –0.0155 | 0.0706 |
| hAQP5 +100 ng hCA II | hAQP5 + Tris | 0.0214 | 2.22 | 0.767 | – | –0.0216 | 0.0645 |
| hAQP5 +100 ng hCA II | H<sub>2</sub>O + 1 ng hCA II | 0.0258 | 2.67 | 0.563 | – | –0.0172 | 0.0689 |
| hAQP5 +100 ng hCA II | hAQP5 + 1 ng hCA II | 0.00417 | 0.431 | 1.000 | – | –0.0389 | 0.0472 |
| hAQP5 +100 ng hCA II | H<sub>2</sub>O + 10 ng hCA II | 0.0147 | 1.52 | 0.960 | – | –0.0284 | 0.0577 |
| hAQP5 +100 ng hCA II | hAQP5 + 10 ng hCA II | –0.0133 | 1.38 | 0.976 | – | –0.0564 | 0.0297 |
| hAQP5 +100 ng hCA II | H<sub>2</sub>O + 100 ng hCA II | 0.0260 | 2.69 | 0.554 | – | –0.0170 | 0.0691 |

**Supplemental Table 6C.** Statistics Summary for **Figure 6*C*, i**) Descriptive Statistics for data presented in Figure 6*C*, β_i_. ii) Overall one-way ANOVA results for data presented in Figure 6*C*. DF is degrees of freedom. iii) Tukey’s means comparison analysis for all data presented in Figure 6*C*. Threshold for significance α=0.05. Standard error of the mean = 0.942. Mean difference for each comparison (Mean Diff.), lower confidence limit (LCL), upper confidence limit (UCL).

|  | n | Mean | Standard Deviation | SE of Mean |
| --- | --- | --- | --- | --- |
| H<sub>2</sub>O + Tris | 8 | 13.4 | 2.20 | 0.779 |
| hAQP5 + Tris | 8 | 14.2 | 2.14 | 0.758 |
| H<sub>2</sub>O + 1 ng hCA II | 8 | 14.1 | 2.07 | 0.731 |
| hAQP5 + 1 ng hCA II | 8 | 12.9 | 1.21 | 0.427 |
| H<sub>2</sub>O + 10 ng hCA II | 8 | 12.1 | 2.03 | 0.718 |
| hAQP5 + 10 ng hCA II | 8 | 11.5 | 1.23 | 0.435 |
| H<sub>2</sub>O + 100 ng hCA II | 8 | 13.1 | 2.26 | 0.799 |
| hAQP5 + 100 ng hCA II | 8 | 11.8 | 1.57 | 0.556 |

*ii. Overall ANOVA Table*
| Variation | DF | Sum of Squares | Mean Square | F-value | P-value |
| --- | --- | --- | --- | --- | --- |
| Between Groups | 7 | 57.6 | 8.22 | 2.32 | 0.0379 |
| Within Groups | 56 | 199 | 3.55 |  |  |
| Total | 63 | 256 |  |  |  |

*iii. Tukey Test*
| Condition A | Condition B | Mean Diff. | Q-value | P-value | Sig. | LCL | UCL |
| --- | --- | --- | --- | --- | --- | --- | --- |
| hAQP5 + Tris | H<sub>2</sub>O + Tris | 0.873 | 1.31 | 0.982 | – | –2.09 | 3.84 |
| H<sub>2</sub>O + 1 ng hCA II | H<sub>2</sub>O + Tris | 0.699 | 1.05 | 0.995 | – | –2.27 | 3.67 |
| H<sub>2</sub>O + 1 ng hCA II | hAQP5 + Tris | –0.173 | 0.260 | 1.000 | – | –3.14 | 2.79 |
| hAQP5 + 1 ng hCA II | H<sub>2</sub>O + Tris | –0.414 | 0.621 | 1.000 | – | –3.38 | 2.55 |
| hAQP5 + 1 ng hCA II | hAQP5 + Tris | –1.29 | 1.93 | 0.869 | – | –4.25 | 1.68 |
| hAQP5 + 1 ng hCA II | H<sub>2</sub>O + 1 ng hCA II | –1.11 | 1.67 | 0.934 | – | –4.08 | 1.85 |
| H<sub>2</sub>O + 10 ng hCA II | H<sub>2</sub>O + Tris | –1.27 | 1.91 | 0.876 | – | –4.24 | 1.70 |
| H<sub>2</sub>O + 10 ng hCA II | hAQP5 + Tris | –2.14 | 3.22 | 0.326 | – | –5.11 | 0.824 |
| H<sub>2</sub>O + 10 ng hCA II | H<sub>2</sub>O + 1 ng hCA II | –1.97 | 2.96 | 0.434 | – | –4.93 | 0.997 |
| H<sub>2</sub>O + 10 ng hCA II | hAQP5 + 1 ng hCA II | –0.856 | 1.28 | 0.984 | – | –3.82 | 2.11 |
| hAQP5 + 10 ng hCA II | H<sub>2</sub>O + Tris | –1.87 | 2.80 | 0.504 | – | –4.83 | 1.10 |
| hAQP5 + 10 ng hCA II | hAQP5 + Tris | –2.74 | 4.11 | 0.0905 | – | –5.70 | 0.227 |
| hAQP5 + 10 ng hCA II | H<sub>2</sub>O + 1 ng hCA II | –2.57 | 3.85 | 0.137 | – | –5.53 | 0.401 |
| hAQP5 + 10 ng<br>hCA II | hAQP5 + 1 ng<br>hCA II | -1.45 | 2.18 | 0.782 | - | -4.42 | 1.51 |
| hAQP5 + 10 ng<br>hCA II | H<sub>2</sub>O + 10ng hCA II | -0.597 | 0.895 | 0.998 | - | -3.56 | 2.37 |
| H<sub>2</sub>O + 100 ng hCA II | H<sub>2</sub>O + Tris | -0.261 | 0.392 | 1.000 | - | -3.23 | 2.71 |
| H<sub>2</sub>O + 100ng hCA II | hAQP5 + Tris | -1.13 | 1.70 | 0.928 | - | -4.10 | 1.83 |
| H<sub>2</sub>O + 100 ng hCA II | H<sub>2</sub>O + 1 ng hCA II | -0.960 | 1.44 | 0.970 | - | -3.93 | 2.01 |
| H<sub>2</sub>O + 100 ng hCA II | hAQP5 + 1ng hCA II | 0.153 | 0.229 | 1.000 | - | -2.81 | 3.12 |
| H<sub>2</sub>O + 100ng hCA II | H<sub>2</sub>O + 10ng hCA II | 1.01 | 1.51 | 0.960 | - | -1.96 | 3.97 |
| H<sub>2</sub>O + 100 ng hCA II | hAQP5 + 10 ng<br>hCA II | 1.60 | 2.41 | 0.685 | - | -1.36 | 4.57 |
| hAQP5 +100 ng<br>hCA II | H<sub>2</sub>O + Tris | -1.55 | 2.33 | 0.720 | - | -4.52 | 1.41 |
| hAQP5 +100 ng<br>hCA II | hAQP5 + Tris | -2.42 | 3.64 | 0.188 | - | -5.39 | 0.542 |
| hAQP5 +100 ng<br>hCA II | H<sub>2</sub>O + 1 ng hCA II | -2.25 | 3.38 | 0.267 | - | -5.22 | 0.715 |
| hAQP5 +100 ng<br>hCA II | hAQP5 + 1 ng<br>hCA II | -1.14 | 1.71 | 0.926 | - | -4.10 | 1.83 |
| hAQP5 +100 ng<br>hCA II | H<sub>2</sub>O + 10 ng hCA II | -0.282 | 0.424 | 1.000 | - | -3.25 | 2.68 |
| hAQP5 +100 ng<br>hCA II | hAQP5 + 10 ng<br>hCA II | 0.314 | 0.472 | 1.000 | - | -2.65 | 3.28 |
| hAQP5 +100 ng<br>hCA II | H<sub>2</sub>O + 100 ng hCA II | -1.29 | 1.94 | 0.867 | - | -4.26 | 1.68 |

**Supplemental Table 6D.** Statistics Summary for **Figure 6*D*, i**) Descriptive Statistics for data presented in Figure 6*D*, *P*_f_. ii) Overall one-way ANOVA results for data presented in Figure 6*D*. DF is degrees of freedom. iii) Tukey’s means comparison analysis for all data presented in Figure 6*D*. Threshold for significance α=0.05. Standard error of the mean = 1.68×10^−4^. Mean difference for each comparison (Mean Diff.), lower confidence limit (LCL), upper confidence limit (UCL).

|  | n | Mean | Standard Deviation | SE of Mean |
| --- | --- | --- | --- | --- |
| H<sub>2</sub>O + Tris | 8 | $1.60 \times 10^{-4}$ | $9.95 \times 10^{-5}$ | $3.52 \times 10^{-5}$ |
| hAQP5 + Tris | 8 | 0.00373 | $4.41 \times 10^{-4}$ | $1.56 \times 10^{-4}$ |
| H<sub>2</sub>O + 1 ng hCA II | 8 | $1.90 \times 10^{-4}$ | $1.10 \times 10^{-4}$ | $3.90 \times 10^{-5}$ |
| hAQP5 + 1 ng hCA II | 8 | 0.00376 | $4.48 \times 10^{-4}$ | $1.58 \times 10^{-4}$ |
| H<sub>2</sub>O + 10 ng hCA II | 8 | $2.35 \times 10^{-4}$ | $2.01 \times 10^{-4}$ | $7.11 \times 10^{-5}$ |
| hAQP5 + 10 ng hCA II | 8 | 0.00388 | $5.32 \times 10^{-4}$ | $1.88 \times 10^{-4}$ |
| H<sub>2</sub>O + 100 ng hCA II | 8 | $1.82 \times 10^{-4}$ | $1.50 \times 10^{-4}$ | $5.31 \times 10^{-5}$ |
| hAQP5 + 100 ng hCA II | 8 | 0.00378 | $3.80 \times 10^{-4}$ | $1.34 \times 10^{-4}$ |

*ii. Overall ANOVA Table*
| Variation | DF | Sum of Squares | Mean Square | F-value | P-value |
| --- | --- | --- | --- | --- | --- |
| Between Groups | 7 | $2.07 \times 10^{-4}$ | $2.96 \times 10^{-5}$ | 261 | $2.53 \times 10^{-4}$ |
| Within Groups | 56 | $6.35 \times 10^{-6}$ | $1.13 \times 10^{-7}$ | | |
| Total | 63 | $2.13 \times 10^{-4}$ | | | |

*iii. Tukey Test*
| Condition A | Condition B | Mean Diff. | Q-value | P-value | Sig. | LCL | UCL |
| --- | --- | --- | --- | --- | --- | --- | --- |
| hAQP5 + Tris | H<sub>2</sub>O + Tris | 0.00357 | 30.0 | $6.86 \times 10^{-8}$ | ✓ | 0.00304 | 0.00410 |
| H<sub>2</sub>O + 1 ng hCA II | H<sub>2</sub>O + Tris | $3.02 \times 10^{-5}$ | 0.254 | 1.00 | – | $-5.00 \times 10^{-4}$ | $5.60 \times 10^{-4}$ |
| H<sub>2</sub>O + 1 ng hCA II | hAQP5 + Tris | -0.00354 | 29.8 | $8.93 \times 10^{-7}$ | ✓ | -0.00407 | -0.00301 |
| hAQP5 + 1 ng hCA II | H<sub>2</sub>O + Tris | 0.00360 | 30.2 | $1.87 \times 10^{-6}$ | ✓ | 0.00307 | 0.00413 |
| hAQP5 + 1 ng hCA II | hAQP5 + Tris | $2.33 \times 10^{-5}$ | 0.196 | 1.00 | – | $-5.07 \times 10^{-4}$ | $5.53 \times 10^{-4}$ |
| hAQP5 + 1 ng hCA II | H<sub>2</sub>O + 1 ng hCA II | 0.00357 | 30.0 | $2.79 \times 10^{-7}$ | ✓ | 0.00304 | 0.00410 |
| H<sub>2</sub>O + 10 ng hCA II | H<sub>2</sub>O + Tris | $7.45 \times 10^{-5}$ | 0.626 | 1.00 | – | $-4.56 \times 10^{-4}$ | $6.05 \times 10^{-4}$ |
| H<sub>2</sub>O + 10 ng hCA II | hAQP5 + Tris | -0.00350 | 29.4 | $8.76 \times 10^{-7}$ | ✓ | -0.00403 | -0.00297 |
| H<sub>2</sub>O + 10 ng hCA II | H<sub>2</sub>O + 1 ng hCA II | $4.43 \times 10^{-5}$ | 0.372 | 1.00 | – | $-4.86 \times 10^{-4}$ | $5.74 \times 10^{-4}$ |
| H<sub>2</sub>O + 10 ng hCA II | hAQP5 + 1 ng hCA II | -0.00352 | 29.6 | $8.85 \times 10^{-7}$ | ✓ | -0.00405 | -0.00299 |
| hAQP5 + 10 ng hCA II | H<sub>2</sub>O + Tris | 0.00372 | 31.2 | $7.30 \times 10^{-8}$ | ✓ | 0.00319 | 0.00425 |
| hAQP5 + 10 ng hCA II | hAQP5 + Tris | $1.46 \times 10^{-4}$ | 1.23 | 0.988 | – | $-3.84 \times 10^{-4}$ | $6.77 \times 10^{-4}$ |
| hAQP5 + 10 ng hCA II | H<sub>2</sub>O + 1 ng hCA II | 0.00369 | 31.0 | $7.23 \times 10^{-8}$ | ✓ | 0.00316 | 0.00422 |
| hAQP5 + 10 ng hCA II | hAQP5 + 1 ng hCA II | $1.23 \times 10^{-4}$ | 1.03 | 0.996 | – | $-4.07 \times 10^{-4}$ | $6.53 \times 10^{-4}$ |
| hAQP5 + 10 ng hCA II | H<sub>2</sub>O + 10ng hCA II | 0.00364 | 30.6 | $7.16 \times 10^{-8}$ | ✓ | 0.00311 | 0.00418 |
| H<sub>2</sub>O + 100 ng hCA II | H<sub>2</sub>O + Tris | $2.18 \times 10^{-5}$ | 0.183 | 1.000 | – | $-5.08 \times 10^{-4}$ | $5.52 \times 10^{-4}$ |
| H<sub>2</sub>O + 100ng hCA II | hAQP5 + Tris | –0.00355 | 29.8 | $8.96 \times 10^{-7}$ | ✓ | –0.00408 | –0.00302 |
| H<sub>2</sub>O + 100 ng hCA II | H<sub>2</sub>O + 1 ng hCA II | $-8.37 \times 10^{-6}$ | 0.0703 | 1.00 | – | $-5.38 \times 10^{-4}$ | $5.22 \times 10^{-4}$ |
| H<sub>2</sub>O + 100 ng hCA II | hAQP5 + 1 ng hCA II | –0.00357 | 30.0 | $6.86 \times 10^{-8}$ | ✓ | –0.00410 | –0.00304 |
| H<sub>2</sub>O + 100ng hCA II | H<sub>2</sub>O + 10 ng hCA II | $-5.27 \times 10^{-5}$ | 0.442 | 1.00 | – | $-5.83 \times 10^{-4}$ | $4.77 \times 10^{-4}$ |
| H<sub>2</sub>O + 100 ng hCA II | hAQP5 + 10 ng hCA II | –0.00370 | 31.1 | $7.25 \times 10^{-8}$ | ✓ | –0.00423 | –0.00317 |
| hAQP5 +100 ng hCA II | H<sub>2</sub>O + Tris | 0.00362 | 30.4 | $7.08 \times 10^{-8}$ | ✓ | 0.00309 | 0.00415 |
| hAQP5 +100 ng hCA II | hAQP5 + Tris | $4.36 \times 10^{-5}$ | 0.366 | 1.00 | – | $-4.87 \times 10^{-4}$ | $5.74 \times 10^{-4}$ |
| hAQP5 +100 ng hCA II | H<sub>2</sub>O + 1 ng hCA II | 0.00359 | 30.1 | $1.86 \times 10^{-6}$ | ✓ | 0.00306 | 0.00412 |
| hAQP5 +100 ng hCA II | hAQP5 + 1 ng hCA II | $2.03 \times 10^{-5}$ | 0.170 | 1.00 | – | $-5.10 \times 10^{-4}$ | $5.50 \times 10^{-4}$ |
| hAQP5 +100 ng hCA II | H<sub>2</sub>O + 10 ng hCA II | 0.00354 | 29.7 | $8.93 \times 10^{-7}$ | ✓ | 0.00301 | 0.00407 |
| hAQP5 +100 ng hCA II | hAQP5 + 10 ng hCA II | $-1.03 \times 10^{-4}$ | 0.864 | 0.999 | – | $-6.33 \times 10^{-4}$ | $4.27 \times 10^{-4}$ |
| hAQP5 +100 ng hCA II | H<sub>2</sub>O + 100 ng hCA II | 0.00359 | 30.2 | $1.87 \times 10^{-6}$ | ✓ | 0.00306 | 0.00412 |

**Supplemental Table 8A.** Statistics Summary for **Figure 8*A*, i**) Descriptive Statistics for data presented in Figure 8*A*, ΔpHS upon CO2 addition. ii) Overall one-way ANOVA results for data presented in **Figure 8***A*. DF is degrees of freedom. iii) Tukey’s means comparison analysis for all data presented in **Figure 8***A*. Threshold for significance α=0.05. Standard error of the mean = 0.0235. Mean difference for each comparison (Mean Diff.), lower confidence limit (LCL), upper confidence limit (UCL).

|  | n | Mean | Standard Deviation | SE of Mean |
| --- | --- | --- | --- | --- |
| H<sub>2</sub>O + Tris | 6 | 0.235 | 0.235 | 0.0112 |
| hAQP5 + Tris | 6 | 0.236 | 0.236 | 0.00784 |
| H<sub>2</sub>O + 1ng hCA II | 6 | 0.324 | 0.324 | 0.0196 |
| hAQP5 + 1ng hCA II | 6 | 0.347 | 0.347 | 0.0230 |

ii. *Overall ANOVA Table*
| Variation | DF | Sum of Squares | Mean Square | F-value | P-value |
| --- | --- | --- | --- | --- | --- |
| Between Groups | 3 | 0.0614 | 0.0205 | 12.4 | 8.35×10<sup>-5</sup> |
| Within Groups | 20 | 0.0330 | 0.00165 |  |  |
| Total | 23 | 0.0944 |  |  |  |

iii. *Tukey Test*
| Condition A | Condition B | Mean Diff. | Q-value | P-value | Sig. | LCL | UCL |
| --- | --- | --- | --- | --- | --- | --- | --- |
| hAQP5 + Tris | H<sub>2</sub>O + Tris | 0.00134 | 0.0810 | 1.000 | — | -0.0643 | 0.0670 |
| H<sub>2</sub>O + 1 ng hCA II | H<sub>2</sub>O + Tris | 0.0889 | 5.36 | 0.00582 | ✓ | 0.0232 | 0.155 |
| H<sub>2</sub>O + 1 ng hCA II | hAQP5 + Tris | 0.0875 | 5.28 | 0.00663 | ✓ | 0.0219 | 0.153 |
| hAQP5 + 1 ng hCA II | H<sub>2</sub>O + Tris | 0.112 | 6.76 | 6.13×10<sup>-4</sup> | ✓ | 0.0464 | 0.178 |
| hAQP5 + 1 ng hCA II | hAQP5 + Tris | 0.111 | 6.68 | 6.99×10<sup>-4</sup> | ✓ | 0.0451 | 0.176 |
| hAQP5 + 1 ng hCA II | H<sub>2</sub>O + 1 ng hCA II | 0.0232 | 1.40 | 0.757 | — | -0.0425 | 0.0888 |

**Supplemental Table 8B.** Statistics Summary for **Figure 8*B*, i**) Descriptive Statistics for data presented in Figure 8*B*, ΔpH_S_ upon CO_2_ removal. ii) Overall one-way ANOVA results for data presented in Figure 8*B*. DF is degrees of freedom. iii) Tukey’s means comparison analysis for all data presented in Figure 8*B*. Threshold for significance α=0.05. Standard error of the mean = 0.0282. Mean difference for each comparison (Mean Diff.), lower confidence limit (LCL), upper confidence limit (UCL).

|  | n | Mean | Standard Deviation | SE of Mean |
| --- | --- | --- | --- | --- |
| H<sub>2</sub>O + Tris | 6 | -0.335 | 0.0523 | 0.0213 |
| hAQP5 + Tris | 6 | -0.341 | 0.0314 | 0.0128 |
| H<sub>2</sub>O + 1 ng hCA II | 6 | -0.418 | 0.0546 | 0.0223 |
| hAQP5 + 1 ng hCA II | 6 | -0.450 | 0.0531 | 0.0217 |

*ii. Overall ANOVA Table*
| Variation | DF | Sum of Squares | Mean Square | F-value | P-value |
| --- | --- | --- | --- | --- | --- |
| Between Groups | 3 | 0.0590 | 0.0197 | 8.26 | $9.03 \times 10^{-4}$ |
| Within Groups | 20 | 0.0476 | 0.00238 |  |  |
| Total | 23 | 0.107 |  |  |  |

*iii. Tukey Test*
| Condition A | Condition B | Mean Diff. | Q-value | P-value | Sig. | LCL | UCL |
| --- | --- | --- | --- | --- | --- | --- | --- |
| hAQP5 + Tris | H<sub>2</sub>O + Tris | -0.00633 | 0.318 | 0.996 | — | -0.0852 | 0.0725 |
| H<sub>2</sub>O + 1 ng hCA II | H<sub>2</sub>O + Tris | -0.0836 | 4.20 | 0.0352 | ✓ | -0.162 | -0.00478 |
| H<sub>2</sub>O + 1 ng hCA II | hAQP5 + Tris | -0.0773 | 3.88 | 0.0560 | — | -0.156 | 0.00156 |
| hAQP5 + 1 ng hCA II | H<sub>2</sub>O + Tris | -0.116 | 5.80 | 0.00287 | ✓ | -0.194 | -0.0367 |
| hAQP5 + 1 ng hCA II | hAQP5 + Tris | -0.109 | 5.48 | 0.00478 | ✓ | -0.188 | -0.0304 |
| hAQP5 + 1 ng hCA II | H<sub>2</sub>O + 1 ng hCA II | -0.0319 | 1.60 | 0.674 | — | -0.111 | 0.0469 |

**Supplemental Table 8C.** Statistics Summary for **Figure 8*C*, i**) Descriptive Statistics for data presented in Figure 8*C*, (dpH_i_/dt)_Max_ upon CO_2_ addition. ii) Overall one-way ANOVA results for data presented in Figure 8*C*. DF is degrees of freedom. iii) Tukey’s means comparison analysis for all data presented in Figure 8*C*. Threshold for significance α=0.05. Standard error of the mean = 6.72×10^−4^. Mean difference for each comparison (Mean Diff.), lower confidence limit (LCL), upper confidence limit (UCL).

|  | n | Mean | Standard Deviation | SE of Mean |
| --- | --- | --- | --- | --- |
| H<sub>2</sub>O + Tris | 6 | -0.00250 | $7.09 \times 10^{-4}$ | $2.89 \times 10^{-4}$ |
| hAQP5 + Tris | 6 | -0.00451 | 0.00110 | $4.48 \times 10^{-4}$ |
| H<sub>2</sub>O + 1 ng hCA II | 6 | -0.00742 | $9.98 \times 10^{-4}$ | $4.08 \times 10^{-4}$ |
| hAQP5 + 1 ng hCA II | 6 | -0.00670 | 0.00165 | $6.73 \times 10^{-4}$ |

*ii. Overall ANOVA Table*
| Variation | DF | Sum of Squares | Mean Square | F-value | P-value |
| --- | --- | --- | --- | --- | --- |
| Between Groups | 3 | $8.95 \times 10^{-5}$ | $2.98 \times 10^{-5}$ | 22.0 | $1.50 \times 10^{-6}$ |
| Within Groups | 20 | $2.71 \times 10^{-5}$ | $1.35 \times 10^{-6}$ | | |
| Total | 23 | $1.17 \times 10^{-4}$ | | | |

*iii. Tukey Test*
| Condition A | Condition B | Mean Diff. | Q-value | P-value | Sig. | LCL | UCL |
| --- | --- | --- | --- | --- | --- | --- | --- |
| hAQP5 + Tris | H<sub>2</sub>O + Tris | -0.00201 | 4.24 | 0.0331 | ✓ | -0.00389 | $-1.33 \times 10^{-4}$ |
| H<sub>2</sub>O + 1 ng hCA II | H<sub>2</sub>O + Tris | -0.00492 | 10.4 | $2.42 \times 10^{-6}$ | ✓ | -0.00680 | -0.00304 |
| H<sub>2</sub>O + 1 ng hCA II | hAQP5 + Tris | -0.00291 | 6.12 | 0.00171 | ✓ | -0.00479 | -0.00103 |
| hAQP5 + 1 ng hCA II | H<sub>2</sub>O + Tris | -0.00420 | 8.84 | $2.33 \times 10^{-5}$ | ✓ | -0.00608 | -0.00232 |
| hAQP5 + 1 ng hCA II | hAQP5 + Tris | -0.00218 | 4.60 | 0.0192 | ✓ | -0.00407 | $-3.05 \times 10^{-4}$ |
| hAQP5 + 1 ng hCA II | H<sub>2</sub>O + 1 ng hCA II | $7.23 \times 10^{-4}$ | 1.52 | 0.707 | — | -0.00116 | 0.00260 |

**Supplemental Table 8D.**
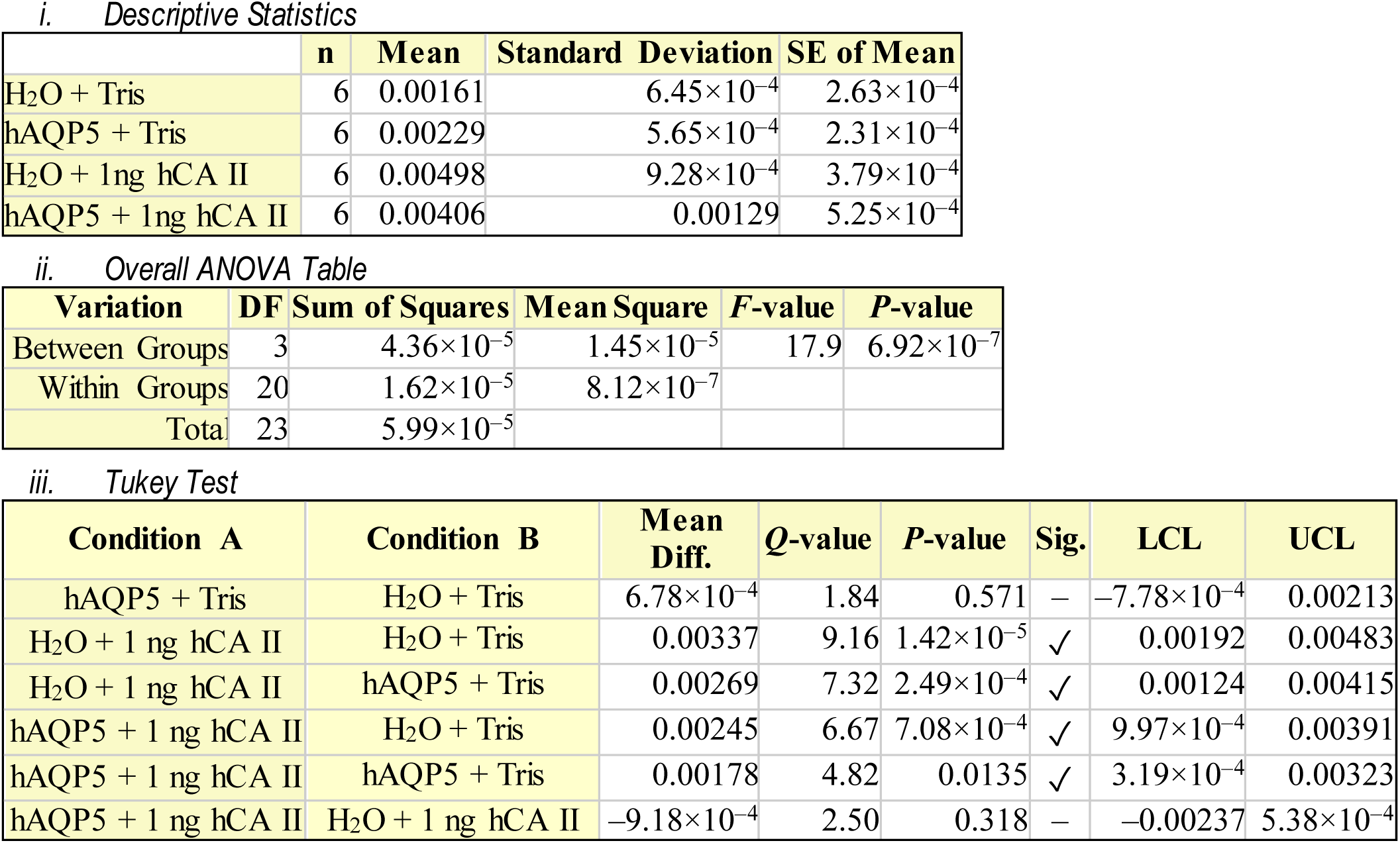
Statistics Summary for **Figure 8*D*, i**) Descriptive Statistics for data presented in Figure 8*D*, (dpH_i_/dt)_Max_ upon CO_2_ removal. ii) Overall one-way ANOVA results for data presented in Figure 8*D*. DF is degrees of freedom. iii) Tukey’s means comparison analysis for all data presented in Figure 8*D*. Threshold for significance α=0.05. Standard error of the mean = 5.20×10^−4^. Mean difference for each comparison (Mean Diff.), lower confidence limit (LCL), upper confidence limit (UCL).

|  | n | Mean | Standard Deviation | SE of Mean |
| --- | --- | --- | --- | --- |
| H<sub>2</sub>O + Tris | 6 | 0.00161 | $6.45 \times 10^{-4}$ | $2.63 \times 10^{-4}$ |
| hAQP5 + Tris | 6 | 0.00229 | $5.65 \times 10^{-4}$ | $2.31 \times 10^{-4}$ |
| H<sub>2</sub>O + 1ng hCA II | 6 | 0.00498 | $9.28 \times 10^{-4}$ | $3.79 \times 10^{-4}$ |
| hAQP5 + 1ng hCA II | 6 | 0.00406 | 0.00129 | $5.25 \times 10^{-4}$ |

*ii. Overall ANOVA Table*
| Variation | DF | Sum of Squares | Mean Square | F-value | P-value |
| --- | --- | --- | --- | --- | --- |
| Between Groups | 3 | $4.36 \times 10^{-5}$ | $1.45 \times 10^{-5}$ | 17.9 | $6.92 \times 10^{-7}$ |
| Within Groups | 20 | $1.62 \times 10^{-5}$ | $8.12 \times 10^{-7}$ | | |
| Total | 23 | $5.99 \times 10^{-5}$ | | | |

*iii. Tukey Test*
| Condition A | Condition B | Mean Diff. | Q-value | P-value | Sig. | LCL | UCL |
| --- | --- | --- | --- | --- | --- | --- | --- |
| hAQP5 + Tris | H<sub>2</sub>O + Tris | $6.78 \times 10^{-4}$ | 1.84 | 0.571 | — | $-7.78 \times 10^{-4}$ | 0.00213 |
| H<sub>2</sub>O + 1 ng hCA II | H<sub>2</sub>O + Tris | 0.00337 | 9.16 | $1.42 \times 10^{-5}$ | ✓ | 0.00192 | 0.00483 |
| H<sub>2</sub>O + 1 ng hCA II | hAQP5 + Tris | 0.00269 | 7.32 | $2.49 \times 10^{-4}$ | ✓ | 0.00124 | 0.00415 |
| hAQP5 + 1 ng hCA II | H<sub>2</sub>O + Tris | 0.00245 | 6.67 | $7.08 \times 10^{-4}$ | ✓ | $9.97 \times 10^{-4}$ | 0.00391 |
| hAQP5 + 1 ng hCA II | hAQP5 + Tris | 0.00178 | 4.82 | 0.0135 | ✓ | $3.19 \times 10^{-4}$ | 0.00323 |
| hAQP5 + 1 ng hCA II | H<sub>2</sub>O + 1 ng hCA II | $-9.18 \times 10^{-4}$ | 2.50 | 0.318 | — | -0.00237 | $5.38 \times 10^{-4}$ |

**Supplemental Table 9A.** Statistics Summary for **Figure 9*A*, i**) Descriptive Statistics for data presented in **Figure 9*A***, (dpH_S_/d_t_)_Max_ upon CO_2_ addition. ii) Overall one-way ANOVA results for data presented in **Figure 9*A***. DF is degrees of freedom. iii) Tukey’s means comparison analysis for all data presented in **Figure 9*A***. Threshold for significance α=0.05. Standard error of the mean = 8.10 ×10–4. Mean difference for each comparison (Mean Diff.), lower confidence limit (LCL), upper confidence limit (UCL).

|  | n | Mean | Standard Deviation | SE of Mean |
| --- | --- | --- | --- | --- |
| H<sub>2</sub>O + Tris | 6 | -0.00502 | 0.00137 | $5.58 \times 10^{-4}$ |
| hAQP5 + Tris | 6 | -0.00541 | 0.00185 | $7.53 \times 10^{-4}$ |
| H<sub>2</sub>O + 1 ng hCA II | 6 | -0.00910 | $9.11 \times 10^{-4}$ | $3.72 \times 10^{-4}$ |
| hAQP5 + 1 ng hCA II | 6 | -0.00843 | 0.00133 | $5.43 \times 10^{-4}$ |

ii. *Overall ANOVA Table*
| Variation | DF | Sum of Squares | Mean Square | F-value | P-value |
| --- | --- | --- | --- | --- | --- |
| Between Groups | 3 | $7.74 \times 10^{-5}$ | $2.58 \times 10^{-5}$ | 13.1 | $5.83 \times 10^{-5}$ |
| Within Groups | 20 | $3.94 \times 10^{-5}$ | $1.97 \times 10^{-5}$ | | |
| Total | 23 | $1.17 \times 10^{-5}$ | | | |

iii. *Tukey Test*
| Condition A | Condition B | Mean Diff. | Q-value | P-value | Sig. | LCL | UCL |
| --- | --- | --- | --- | --- | --- | --- | --- |
| hAQP5 + Tris | H<sub>2</sub>O + Tris | $-3.98 \times 10^{-4}$ | 0.695 | 0.960 | — | -0.00267 | 0.00187 |
| H<sub>2</sub>O + 1 ng hCA II | H<sub>2</sub>O + Tris | -0.00408 | 7.13 | $3.38 \times 10^{-4}$ | ✓ | -0.00635 | -0.00182 |
| H<sub>2</sub>O + 1 ng hCA II | hAQP5 + Tris | -0.00369 | 6.43 | 0.00103 | ✓ | -0.00595 | -0.00142 |
| hAQP5 + 1 ng hCA II | H<sub>2</sub>O + Tris | -0.00342 | 5.96 | 0.00221 | ✓ | -0.00568 | -0.00115 |
| hAQP5 + 1 ng hCA II | hAQP5 + Tris | -0.00302 | 5.27 | 0.00674 | ✓ | -0.00528 | $-7.50 \times 10^{-4}$ |
| hAQP5 + 1 ng hCA II | H<sub>2</sub>O + 1 ng hCA II | $6.68 \times 10^{-4}$ | 1.17 | 0.842 | — | -0.00160 | 0.00294 |

**Supplemental Table 9B.** Statistics Summary for **Figure 9*B***, i) Descriptive Statistics for data presented in **Figure 9*B***, (dpH_S_/dt)_Max_ upon CO_2_ removal. ii) Overall one-way ANOVA results for data presented in **Figure 9*B***. DF is degrees of freedom. iii) Tukey’s means comparison analysis for all data presented in **Figure 9*B***. Threshold for significance α=0.05. Standard error of the mean = 7.71 ×10^−4^. Mean difference for each comparison (Mean Diff.), lower confidence limit (LCL), upper confidence limit (UCL).

|  | n | Mean | Standard Deviation | SE of Mean |
| --- | --- | --- | --- | --- |
| H<sub>2</sub>O + Tris | 6 | 0.00271 | $9.14 \times 10^{-4}$ | $3.73 \times 10^{-4}$ |
| hAQP5 + Tris | 6 | 0.00507 | 0.00180 | $7.34 \times 10^{-4}$ |
| H<sub>2</sub>O + 1 ng hCA II | 6 | 0.00364 | 0.00127 | $5.18 \times 10^{-4}$ |
| hAQP5 + 1 ng hCA II | 6 | 0.00479 | 0.00120 | $4.91 \times 10^{-4}$ |

ii. *Overall ANOVA Table*
| Variation | DF | Sum of Squares | Mean Square | F-value | P-value |
| --- | --- | --- | --- | --- | --- |
| Between Groups | 3 | $2.13 \times 10^{-5}$ | $7.12 \times 10^{-6}$ | 3.99 | 0.0222 |
| Within Groups | 20 | $3.56 \times 10^{-5}$ | $1.78 \times 10^{-6}$ | | |
| Total | 23 | $5.70 \times 10^{-5}$ | | | |

iii. *Tukey Test*
| Condition A | Condition B | Mean Diff. | Q-value | P-value | Sig. | LCL | UCL |
| --- | --- | --- | --- | --- | --- | --- | --- |
| hAQP5 + Tris | H<sub>2</sub>O + Tris | 0.00236 | 4.33 | 0.0288 | ✓ | $2.04 \times 10^{-4}$ | 0.00452 |
| H<sub>2</sub>O + 1 ng hCA II | H<sub>2</sub>O + Tris | $9.36 \times 10^{-4}$ | 1.72 | 0.625 | – | –0.00122 | 0.00309 |
| H<sub>2</sub>O + 1 ng hCA II | hAQP5 + Tris | –0.00143 | 2.62 | 0.281 | – | –0.00358 | $7.32 \times 10^{-4}$ |
| hAQP5 + 1 ng hCA II | H<sub>2</sub>O + Tris | 0.00209 | 3.83 | 0.0604 | – | $-7.19 \times 10^{-5}$ | 0.00424 |
| hAQP5 + 1 ng hCA II | hAQP5 + Tris | $-2.76 \times 10^{-4}$ | 0.506 | 0.984 | – | –0.00243 | 0.00188 |
| hAQP5 + 1 ng hCA II | H<sub>2</sub>O + 1 ng hCA II | 0.00115 | 2.11 | 0.461 | – | –0.00101 | 0.00331 |

**Supplemental Table 10A.** Statistics Summary for **Figure 10*A*, i**) Descriptive Statistics for data presented in Figure 10*A*, initial pH_i_ values. ii) Overall one-way ANOVA results for data presented in Figure 10*A*. DF is degrees of freedom. iii) Tukey’s means comparison analysis for all data presented in Figure 10*A*. Threshold for significance α=0.05. Standard error of the mean = 0.0353. Mean difference for each comparison (Mean Diff.), lower confidence limit (LCL), upper confidence limit (UCL).

|  | **n** | **Mean** | **Standard Deviation** | **SE of Mean** |
| --- | --- | --- | --- | --- |
| H<sub>2</sub>O + Tris | 6 | 7.29 | 0.0530 | 0.0216 |
| hAQP5 + Tris | 6 | 7.32 | 0.0741 | 0.0303 |
| H<sub>2</sub>O + 1 ng hCA II | 6 | 7.28 | 0.0685 | 0.0280 |
| hAQP5 + 1 ng hCA II | 6 | 7.31 | 0.0441 | 0.0180 |

*ii. Overall ANOVA Table*
| **Variation** | **DF** | **Sum of Squares** | **Mean Square** | **F-value** | **P-value** |
| --- | --- | --- | --- | --- | --- |
| Between Groups | 3 | 0.00556 | 0.00185 | 0.497 | 0.689 |
| Within Groups | 20 | 0.0747 | 0.00373 |  |  |
| Total | 23 | 0.0802 |  |  |  |

*iii. Tukey Test*
| **Condition A** | **Condition B** | **Mean Diff.** | **Q-value** | **P-value** | **Sig.** | **LCL** | **UCL** |
| --- | --- | --- | --- | --- | --- | --- | --- |
| hAQP5 + Tris | H<sub>2</sub>O + Tris | 0.0352 | 1.41 | 0.752 | – | –0.0635 | 0.134 |
| H<sub>2</sub>O + 1 ng hCA II | H<sub>2</sub>O + Tris | –0.00157 | 0.0629 | 1.000 | – | –0.100 | 0.0972 |
| H<sub>2</sub>O + 1 ng hCA II | hAQP5 + Tris | –0.0368 | 1.47 | 0.727 | – | –0.136 | 0.0620 |
| hAQP5 + 1 ng hCA II | H<sub>2</sub>O + Tris | 0.0203 | 0.813 | 0.939 | – | –0.0785 | 0.119 |
| hAQP5 + 1 ng hCA II | hAQP5 + Tris | –0.0149 | 0.599 | 0.974 | – | –0.114 | 0.0838 |
| hAQP5 + 1 ng hCA II | H<sub>2</sub>O + 1 ng hCA II | 0.0218 | 0.876 | 0.925 | – | –0.0769 | 0.121 |

**Supplemental Table 10B.** Statistics Summary for **Figure 10*B*, i**) Descriptive Statistics for data presented in Figure 10*B*, ΔpH_i_ values. ii) Overall one-way ANOVA results for data presented in Figure 10*B*. DF is degrees of freedom. iii) Tukey’s means comparison analysis for all data presented in Figure 10*B*. Threshold for significance α=0.05. Standard error of the mean = 0.0177. Mean difference for each comparison (Mean Diff.), lower confidence limit (LCL), upper confidence limit (UCL).

|  | n | Mean | Standard Deviation | SE of Mean |
| --- | --- | --- | --- | --- |
| H<sub>2</sub>O + Tris | 6 | -0.275 | 0.0274 | 0.0112 |
| hAQP5 + Tris | 6 | -0.291 | 0.0117 | 0.00477 |
| H<sub>2</sub>O + 1 ng hCA II | 6 | -0.275 | 0.0374 | 0.0153 |
| hAQP5 + 1 ng hCA II | 6 | -0.301 | 0.0383 | 0.0157 |

*ii. Overall ANOVA Table*
| Variation | DF | Sum of Squares | Mean Square | F-value | P-value |
| --- | --- | --- | --- | --- | --- |
| Between Groups | 3 | 0.00297 | $9.90 \times 10^{-4}$ | 1.05 | 0.391 |
| Within Groups | 20 | 0.0188 | $9.40 \times 10^{-4}$ | | |
| Total | 23 | 0.0218 |  |  |  |

*iii. Tukey Test*
| Condition A | Condition B | Mean Diff. | Q-value | P-value | Sig. | LCL | UCL |
| --- | --- | --- | --- | --- | --- | --- | --- |
| hAQP5 + Tris | H<sub>2</sub>O + Tris | -0.0161 | 1.29 | 0.798 | — | -0.0334 | 0.0657 |
| H<sub>2</sub>O + 1 ng hCA II | H<sub>2</sub>O + Tris | $6.38 \times 10^{-4}$ | 0.0510 | 1.000 | — | -0.0502 | 0.0489 |
| H<sub>2</sub>O + 1 ng hCA II | hAQP5 + Tris | 0.0168 | 1.34 | 0.779 | — | -0.0663 | 0.0327 |
| hAQP5 + 1 ng hCA II | H<sub>2</sub>O + Tris | -0.0256 | 2.05 | 0.486 | — | -0.0239 | 0.0752 |
| hAQP5 + 1 ng hCA II | hAQP5 + Tris | -0.00948 | 0.758 | 0.949 | — | -0.0401 | 0.0590 |
| hAQP5 + 1 ng hCA II | H<sub>2</sub>O + 1ng hCA II | -0.0263 | 2.10 | 0.465 | — | -0.0233 | 0.0758 |

**Supplemental Table 10C.** Statistics Summary for **Figure 10*C*, i**) Descriptive Statistics for data presented in Figure 10*C*, intrinsic buffering power (β_i_). ii) Overall one-way ANOVA results for data presented in Figure 10*C*. DF is degrees of freedom. iii) Tukey’s means comparison analysis for all data presented in Figure 10*C*. Threshold for significance α=0.05. Standard error of the mean = 1.02. Mean difference for each comparison (Mean Diff.), lower confidence limit (LCL), upper confidence limit (UCL).

|  | n | Mean | Standard Deviation | SE of Mean |
| --- | --- | --- | --- | --- |
| H<sub>2</sub>O + Tris | 6 | 12.0 | 2.10 | 0.855 |
| hAQP5 + Tris | 6 | 11.7 | 1.63 | 0.664 |
| H<sub>2</sub>O + 1 ng hCA II | 6 | 12.0 | 1.87 | 0.762 |
| hAQP5 + 1 ng hCA II | 6 | 10.8 | 1.43 | 0.586 |

*ii. Overall ANOVA Table*
| Variation | DF | Sum of Squares | Mean Square | F-value | P-value |
| --- | --- | --- | --- | --- | --- |
| Between Groups | 3 | 5.77 | 1.92 | 0.612 | 0.615 |
| Within Groups | 20 | 62.9 | 3.14 |  |  |
| Total | 23 | 68.6 |  |  |  |

*iii. Tukey Test*
| Condition A | Condition B | Mean Diff. | Q-value | P-value | Sig. | LCL | UCL |
| --- | --- | --- | --- | --- | --- | --- | --- |
| hAQP5 + Tris | H<sub>2</sub>O + Tris | -0.265 | 0.366 | 0.994 | — | -3.13 | 2.60 |
| H<sub>2</sub>O + 1 ng hCA II | H<sub>2</sub>O + Tris | -0.0111 | 0.0154 | 1.000 | — | -2.88 | 2.85 |
| H<sub>2</sub>O + 1 ng hCA II | hAQP5 + Tris | 0.253 | 0.350 | 0.994 | — | -2.61 | 3.12 |
| hAQP5 + 1 ng hCA II | H<sub>2</sub>O + Tris | -1.20 | 1.65 | 0.652 | — | -4.06 | 1.67 |
| hAQP5 + 1 ng hCA II | hAQP5 + Tris | -0.932 | 1.29 | 0.799 | — | -3.80 | 1.93 |
| hAQP5 + 1 ng hCA II | H<sub>2</sub>O + 1 ng hCA II | -1.19 | 1.64 | 0.659 | — | -4.05 | 1.68 |

**Supplemental Table 10D.**
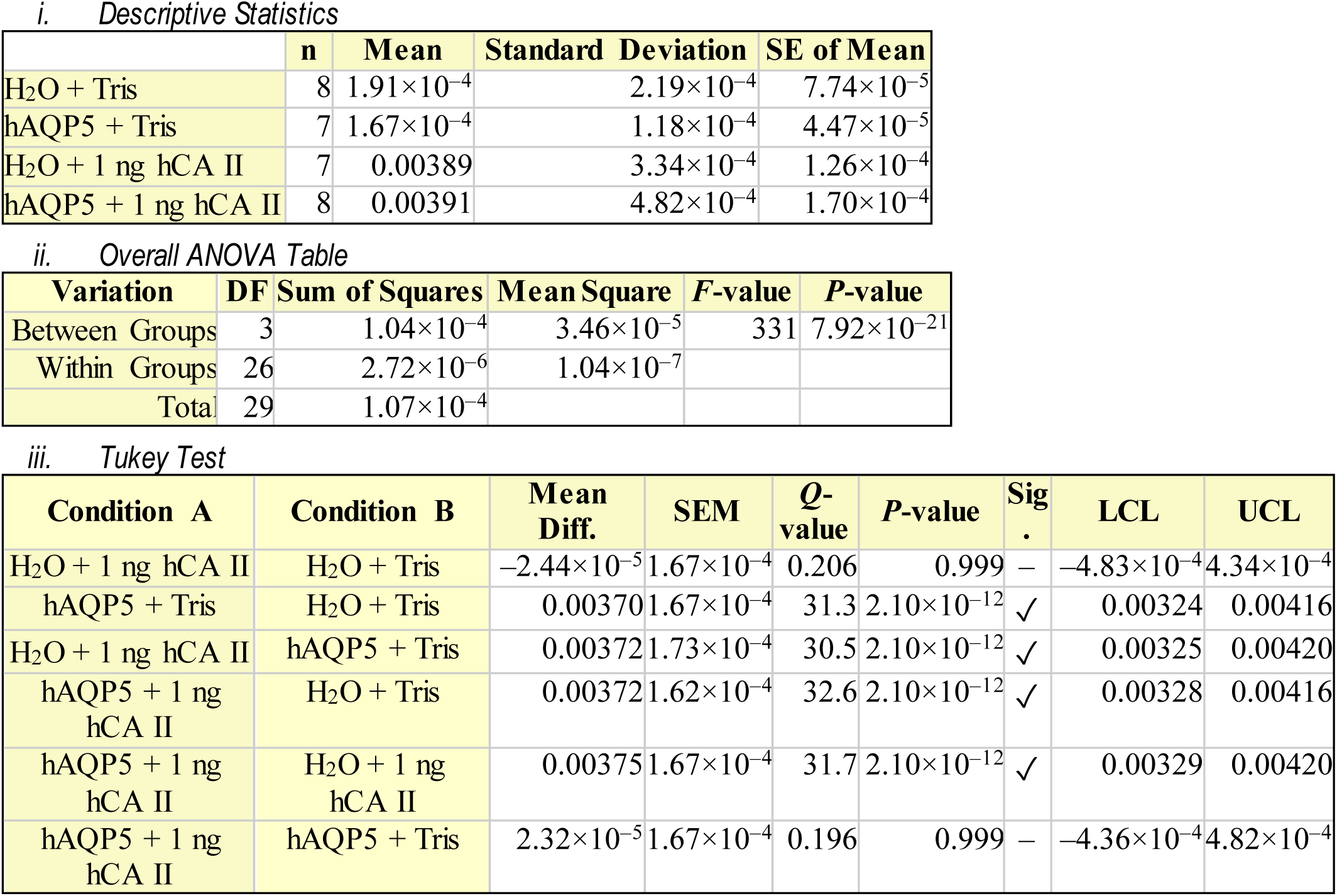
Statistics Summary for **Figure 10*D*, i**) Descriptive Statistics for data presented in Figure 10*D*, *P*_f_. ii) Overall one-way ANOVA results for data presented in Figure 10*D*. DF is degrees of freedom. iii) Tukey’s means comparison analysis for all data presented in Figure 10*D*. Threshold for significance α=0.05. Standard error of the mean for Tukey’s Test (SEM). Mean difference for each comparison (Mean Diff.), lower confidence limit (LCL), upper confidence limit (UCL).

**Supplemental Table 11A.** Statistics Summary for **Figure 11*A*, i**) Descriptive Statistics for data presented in Figure 11*A*, which are the ΔpH_S_ upon CO_2_ addition data rearranged from Figure 3*A* to juxtapose two hAQP5 absent bars and two heterologously expressed hAQP5 bars. ii) Overall one-way ANOVA results for only data bars presented in Figure 11*A* extracted from Figure 3*A*. DF is degrees of freedom. iii) Tukey’s means comparison analysis for all data presented in Figure 11*A*. iv) Descriptive Statistics for data presented in Figure 11*A*, to analyze ΔΔ_Base_ vs. ΔΔ_AQP5_ when adding 1 ng hCA II, on the “trans” side of the membrane to the pH_S_ measurement. v) One-way ANOVA results for ΔΔ_Base_ vs. ΔΔ_AQP5_ data presented in Figure 11*A*. vi) Tukey’s means comparison analysis for ΔΔ_Base_ vs. ΔΔ_AQP5_ data presented in Figure 11*A*. Threshold for significance α=0.05. Standard error of the mean for Tukey’s Test (SEM). Mean difference for each comparison (Mean Diff.), lower confidence limit (LCL), upper confidence limit (UCL).

|  | n | Mean | Standard Deviation | SE of Mean |
| --- | --- | --- | --- | --- |
| H<sub>2</sub>O + Tris | 8 | 0.0285 | 0.00895 | 0.00316 |
| hAQP5 + Tris | 8 | 0.0787 | 0.0181 | 0.00641 |
| H<sub>2</sub>O + 1 ng hCA II | 8 | 0.0411 | 0.0115 | 0.00406 |
| hAQP5 + 1 ng hCA II | 8 | 0.127 | 0.0246 | 0.00869 |

ii. Overall ANOVA Table for $\Delta pH$ s on CO<sub>2</sub> addition
| Variation | DF | Sum of Squares | Mean Square | F-value | P-value |
| --- | --- | --- | --- | --- | --- |
| Between Groups | 3 | 0.0471 | 0.0157 | 54.8 | $7.63 \times 10^{-12}$ |
| Within Groups | 28 | 0.00802 | $2.86 \times 10^{-4}$ | | |
| Total | 31 | 0.0552 |  |  |  |

iii. Tukey Test for $\Delta pH$ s on CO<sub>2</sub> addition
| Condition A | Condition B | Mean Diff. | SEM | Q-value | P-value | Sig. | LCL | UCL |
| --- | --- | --- | --- | --- | --- | --- | --- | --- |
| hAQP5 + Tris | H<sub>2</sub>O + Tris | 0.0502 | 0.00846 | 8.39 | $1.25 \times 10^{-5}$ | ✓ | 0.0271 | 0.0733 |
| H<sub>2</sub>O + 1 ng hCA II | H<sub>2</sub>O + Tris | 0.0126 | 0.00846 | 2.11 | 0.455 | – | –0.0105 | 0.0357 |
| H<sub>2</sub>O + 1 ng hCA II | hAQP5 + Tris | –0.0376 | 0.00846 | 6.28 | $7.00 \times 10^{-4}$ | ✓ | –0.0607 | –0.0145 |
| hAQP5 + 1 ng hCA II | H<sub>2</sub>O + Tris | 0.0986 | 0.00846 | 16.5 | 0.00 | ✓ | 0.0755 | 0.122 |
| hAQP5 + 1 ng hCA II | hAQP5 + Tris | 0.0484 | 0.00846 | 8.09 | $2.20 \times 10^{-5}$ | ✓ | 0.0253 | 0.0715 |
| hAQP5 + 1 ng hCA II | H<sub>2</sub>O + 1 ng hCA II | 0.0860 | 0.00846 | 14.4 | $6.50 \times 10^{-7}$ | ✓ | 0.0629 | 0.109 |

iv. Descriptive Statistics for $\Delta\Delta pH_s$ on CO<sub>2</sub> addition
|  | n | Mean | Standard Deviation | SE of Mean |
| --- | --- | --- | --- | --- |
| $\Delta\Delta_{Base}$ | 8 | 0.0126 | 0.0115 | 0.00406 |
| $\Delta\Delta_{hAQP5}$ | 8 | 0.0484 | 0.0246 | 0.00869 |

v. Overall ANOVA Table for $\Delta\Delta pH_s$ on CO<sub>2</sub> addition
| Variation | DF | Sum of Squares | Mean Square | F-value | P-value |
| --- | --- | --- | --- | --- | --- |
| Between Groups | 1 | 0.00513 | 0.00513 | 13.9 | 0.00223 |
| Within Groups | 14 | 0.00516 | $3.68 \times 10^{-4}$ | | |
| Total | 15 | 0.0103 |  |  |  |

vi. Tukey Test for $\Delta\Delta pH_s$ on CO<sub>2</sub> addition
| Condition A | Condition B | Mean Diff. | SEM | Q-value | P-value | Sig. | LCL | UCL |
| --- | --- | --- | --- | --- | --- | --- | --- | --- |
| $\Delta\Delta_{Base}$ | $\Delta\Delta_{AQP5}$ | 0.0358 | 0.00960 | 5.28 | 0.00224 | ✓ | 0.0152 | 0.0564 |

**Supplemental Table 11B.** Statistics Summary for **Figure 11*B*, i**) Descriptive Statistics for data presented in Figure 11*B*, which are the ΔpH_S_ upon CO_2_ removal data rearranged from Figure 3*B* to juxtapose two hhAQP5 absent bars and two heterologously expressed hAQP5 bars. ii) Overall one-way ANOVA results for only data bars presented in Figure 11*B* extracted from Figure 3*B*. DF is degrees of freedom. iii) Tukey’s means comparison analysis for all data presented in Figure 11*B*. iv) Descriptive Statistics for data presented in Figure 11*B*, to analyze ΔΔ_Base_ vs. ΔΔ_AQP5_ when adding 1 ng hCA II, on the “trans” side of the membrane to the pH_S_ measurement. v) One-way ANOVA results for ΔΔ_Base_ vs. ΔΔ_AQP5_ data presented in Figure 11*B*. vi) Tukey’ s means comparison analysis for ΔΔ_Base_ vs. ΔΔ_AQP5_ data presented in Figure 11*B*. Threshold for significance α=0.05. Standard error of the mean for Tukey’s Test (SEM). Mean difference for each comparison (Mean Diff.), lower confidence limit (LCL), upper confidence limit (UCL).

|  | n | Mean | Standard Deviation | SE of Mean |
| --- | --- | --- | --- | --- |
| H<sub>2</sub>O + Tris | 8 | 0.0285 | 0.00895 | 0.00316 |
| hAQP5 + Tris | 8 | 0.0787 | 0.0181 | 0.00641 |
| H<sub>2</sub>O + 100 ng hCA II | 8 | 0.0416 | 0.00948 | 0.00335 |
| hAQP5 + 100 ng hCA II | 8 | 0.160 | 0.0478 | 0.0169 |

ii. Overall ANOVA Table for $\Delta$ pHs on CO<sub>2</sub> addition
| Variation | DF | Sum of Squares | Mean Square | F-value | P-value |
| --- | --- | --- | --- | --- | --- |
| Between Groups | 3 | 0.0845 | 0.0282 | 40.4 | $2.61 \times 10^{-10}$ |
| Within Groups | 28 | 0.0195 | $6.97 \times 10^{-4}$ | | |
| Total | 31 | 0.104 |  |  |  |

iii. Tukey Test for $\Delta$ pHs on CO<sub>2</sub> addition
| Condition A | Condition B | Mean Diff. | SEM | Q-value | P-value | Sig. | LCL | UCL |
| --- | --- | --- | --- | --- | --- | --- | --- | --- |
| hAQP5 + Tris | H<sub>2</sub>O + Tris | 0.0502 | 0.0132 | 5.38 | 0.00374 | ✓ | 0.0142 | 0.0862 |
| H<sub>2</sub>O + 100 ng hCA II | H<sub>2</sub>O + Tris | 0.0131 | 0.0132 | 1.41 | 0.754 | – | -0.0229 | 0.0492 |
| H<sub>2</sub>O + 100 ng hCA II | hAQP5 + Tris | -0.0371 | 0.0132 | 3.97 | 0.0419 | ✓ | -0.0731 | -0.00104 |
| hAQP5 + 100 ng hCA II | H<sub>2</sub>O + Tris | 0.132 | 0.0132 | 14.1 | $2.23 \times 10^{-7}$ | ✓ | 0.0959 | 0.168 |
| hAQP5 + 100 ng hCA II | hAQP5 + Tris | 0.0817 | 0.0132 | 8.76 | $6.27 \times 10^{-6}$ | ✓ | 0.0457 | 0.118 |
| hAQP5 + 100 ng hCA II | H<sub>2</sub>O + 100 ng hCA II | 0.119 | 0.0132 | 12.7 | 0.00 | ✓ | 0.0828 | 0.155 |

iv. Descriptive Statistics for $\Delta\Delta\text{pH}_s$ on CO<sub>2</sub> addition
|  | n | Mean | Standard Deviation | SE of Mean |
| --- | --- | --- | --- | --- |
| $\Delta\Delta_{\text{Base}}$ | 8 | 0.0131 | 0.00948 | 0.00335 |
| $\Delta\Delta_{\text{hAQP5}}$ | 8 | 0.0817 | 0.0478 | 0.0169 |

v. Overall ANOVA Table
| Variation | DF | Sum of Squares | Mean Square | F-value | P-value |
| --- | --- | --- | --- | --- | --- |
| Between Groups | 1 | 0.0188 | 0.0188 | 15.8 | 0.00137 |
| Within Groups | 14 | 0.0166 | 0.00119 |  |  |
| Total | 15 | 0.0355 |  |  |  |

vi. Tukey Test
| Condition A | Condition B | Mean Diff. | SEM | Q-value | P-value | Sig. | LCL | UCL |
| --- | --- | --- | --- | --- | --- | --- | --- | --- |
| $\Delta\Delta_{\text{Base}}$ | $\Delta\Delta_{\text{AQP5}}$ | 0.0686 | 0.0172 | 5.63 | 0.00137 | ✓ | 0.0316 | 0.106 |

**Supplemental Table 12A.** Statistics Summary for **Figure 12*A*, i**) Descriptive Statistics for data presented in Figure 12*A*, for comparison of select –(dpH_i_/dt)_Max_ bars extracted from Figure 3*C* and Figure 8*C* to analyze –(dpH_i_/dt)_Max_ upon CO_2_ addition in the absence or presence of bCA in bECF, on the “trans” side of the membrane to the pH_i_ measurement. ii) Overall one-way ANOVA results for –(dpH_i_/dt)_Max_ data presented in Figure 12*A*. DF is degrees of freedom. iii) Tukey’ s means comparison analysis for –(dpH_i_/dt)_Max_ the subset of data presented in Figure 12*A*. iv) Descriptive Statistics for data presented in Figure 12*A*, to analyze Δ_Base_ vs. Δ_AQP5_. v) One-way ANOVA results for Δ_Base_ vs. Δ_AQP5_ data presented in Figure 12*A*. vi) Tukey’s means comparison analysis for ΔΔ_Base_ vs. ΔΔ_AQP5_ data presented in Figure 12*A*. Threshold for significance α=0.05. Standard error of the mean for Tukey’s Test (SEM). Mean difference for each comparison (Mean Diff.), lower confidence limit (LCL), upper confidence limit (UCL).

|  | n | Mean | Standard Deviation | SE of Mean |
| --- | --- | --- | --- | --- |
| H<sub>2</sub>O + Tris | 8 | -0.00126 | $2.08 \times 10^{-4}$ | $7.35 \times 10^{-5}$ |
| H<sub>2</sub>O + Tris + bCA<sub>bECF</sub> | 6 | -0.00250 | $7.09 \times 10^{-4}$ | $2.89 \times 10^{-4}$ |
| hAQP5 + Tris | 8 | -0.00122 | $1.57 \times 10^{-4}$ | $5.55 \times 10^{-5}$ |
| hAQP5 + Tris+ bCA<sub>bECF</sub> | 6 | -0.00451 | 0.00110 | $4.48 \times 10^{-4}$ |

ii. Overall ANOVA Table for $-(dpH/dt)_{Max}$ on CO<sub>2</sub> addition
| Variation | DF | Sum of Squares | Mean Square | F-value | P-value |
| --- | --- | --- | --- | --- | --- |
| Between Groups | 3 | $4.73 \times 10^{-5}$ | $1.58 \times 10^{-5}$ | 42.0 | $1.04 \times 10^{-9}$ |
| Within Groups | 24 | $9.00 \times 10^{-6}$ | $3.75 \times 10^{-7}$ | | |
| Total | 27 | $5.63 \times 10^{-5}$ | | | |

iii. Tukey Test for $-(dpH/dt)_{Max}$ on CO<sub>2</sub> addition
| Condition A | Condition B | Mean Diff. | SEM | Q-value | P-value | Sig. | LCL | UCL |
| --- | --- | --- | --- | --- | --- | --- | --- | --- |
| H<sub>2</sub>O + Tris + bCA<sub>bECF</sub> | H<sub>2</sub>O + Tris | -0.00124 | $3.31 \times 10^{-4}$ | 5.30 | 0.00513 | ✓ | -0.00215 | $-3.27 \times 10^{-4}$ |
| hAQP5 + Tris | H<sub>2</sub>O + Tris | $3.53 \times 10^{-5}$ | $3.06 \times 10^{-4}$ | 0.163 | 0.999 | – | $-8.09 \times 10^{-4}$ | $8.80 \times 10^{-4}$ |
| hAQP5 + Tris | H<sub>2</sub>O + Tris + bCA<sub>bECF</sub> | 0.00127 | $3.31 \times 10^{-4}$ | 5.45 | 0.00396 | ✓ | $3.62 \times 10^{-4}$ | 0.00219 |
| hAQP5 + Tris+ bCA<sub>bECF</sub> | H<sub>2</sub>O + Tris | -0.00325 | $3.31 \times 10^{-4}$ | 13.9 | $4.43 \times 10^{-7}$ | ✓ | -0.00417 | -0.00234 |
| hAQP5 + Tris+ bCA<sub>bECF</sub> | H<sub>2</sub>O + Tris + bCA<sub>bECF</sub> | -0.00201 | $3.54 \times 10^{-4}$ | 8.05 | $4.07 \times 10^{-5}$ | ✓ | -0.00299 | -0.00104 |
| hAQP5 + Tris+ bCA<sub>bECF</sub> | hAQP5 + Tris | -0.00329 | $3.31 \times 10^{-4}$ | 14.1 | $4.47 \times 10^{-7}$ | ✓ | -0.00420 | -0.00238 |

iv. Descriptive Statistics for $\Delta_{hhAQP5}$ VS $\Delta_{Base}$ for $-(dpH/dt)_{Max}$ on CO<sub>2</sub> addition
|  | n | Mean | Standard Deviation | SE of Mean |
| --- | --- | --- | --- | --- |
| $\Delta_{Base}$ | 6 | -0.00124 | $7.09 \times 10^{-4}$ | $2.89 \times 10^{-4}$ |
| $\Delta_{AQP5}$ | 6 | -0.00329 | 0.00110 | $4.48 \times 10^{-4}$ |

v. Overall ANOVA Table
| Variation | DF | Sum of Squares | Mean Square | F-value | P-value |
| --- | --- | --- | --- | --- | --- |
| Between Groups | 1 | $1.26 \times 10^{-5}$ | $1.26 \times 10^{-5}$ | 14.8 | 0.00325 |
| Within Groups | 10 | $8.53 \times 10^{-6}$ | $8.53 \times 10^{-7}$ | | |
| Total | 11 | $2.11 \times 10^{-5}$ | | | |

vi. Tukey Test
| Condition A | Condition B | Mean Diff. | SEM | Q-value | P-value | Sig. | LCL | UCL |
| --- | --- | --- | --- | --- | --- | --- | --- | --- |
| $\Delta_{Base}$ | $\Delta_{AQP5}$ | -0.00205 | $5.33 \times 10^{-4}$ | 5.43 | 0.00325 | ✓ | -0.00324 | $-8.61 \times 10^{-4}$ |

**Supplemental Table 12B.**
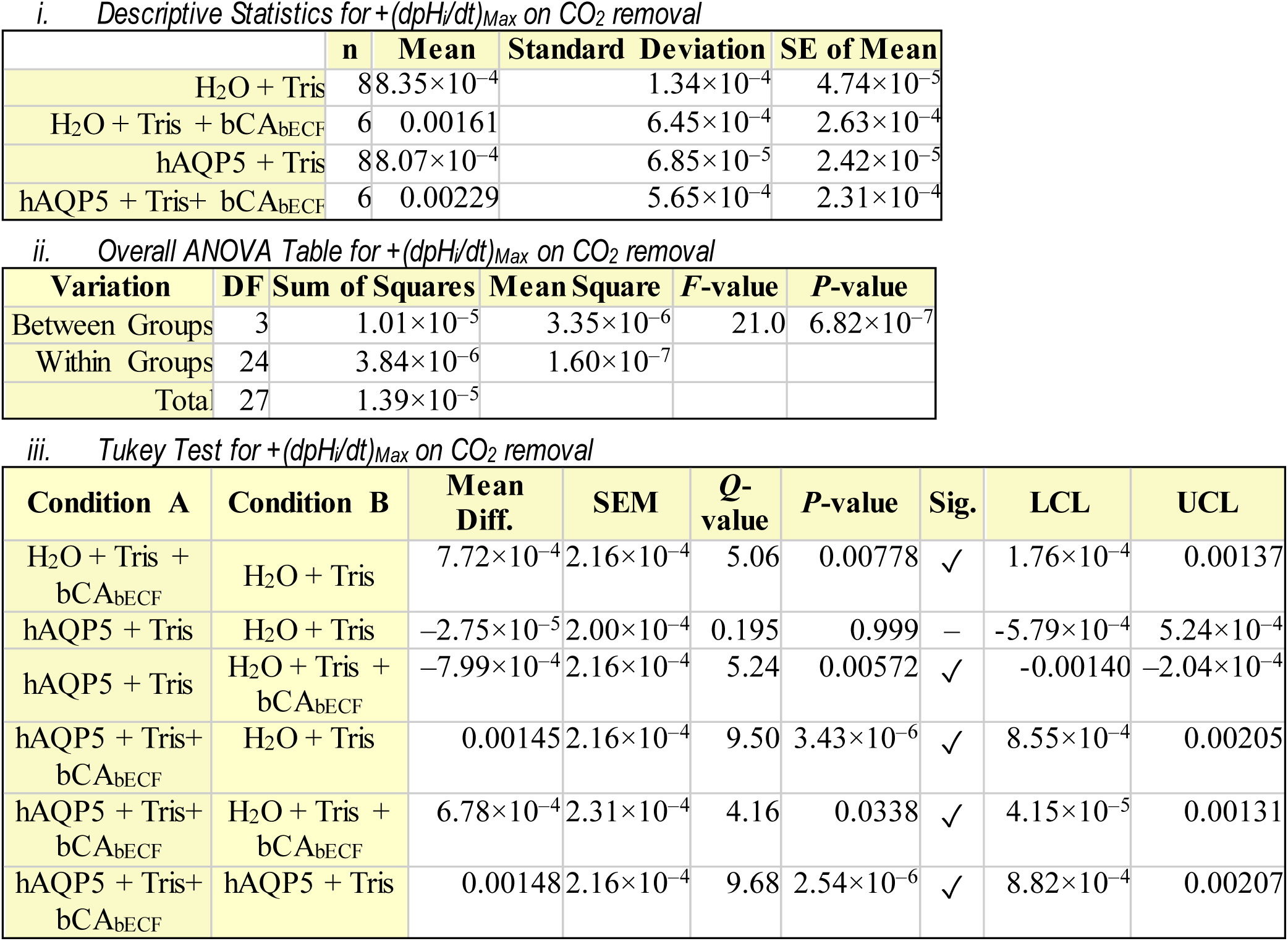
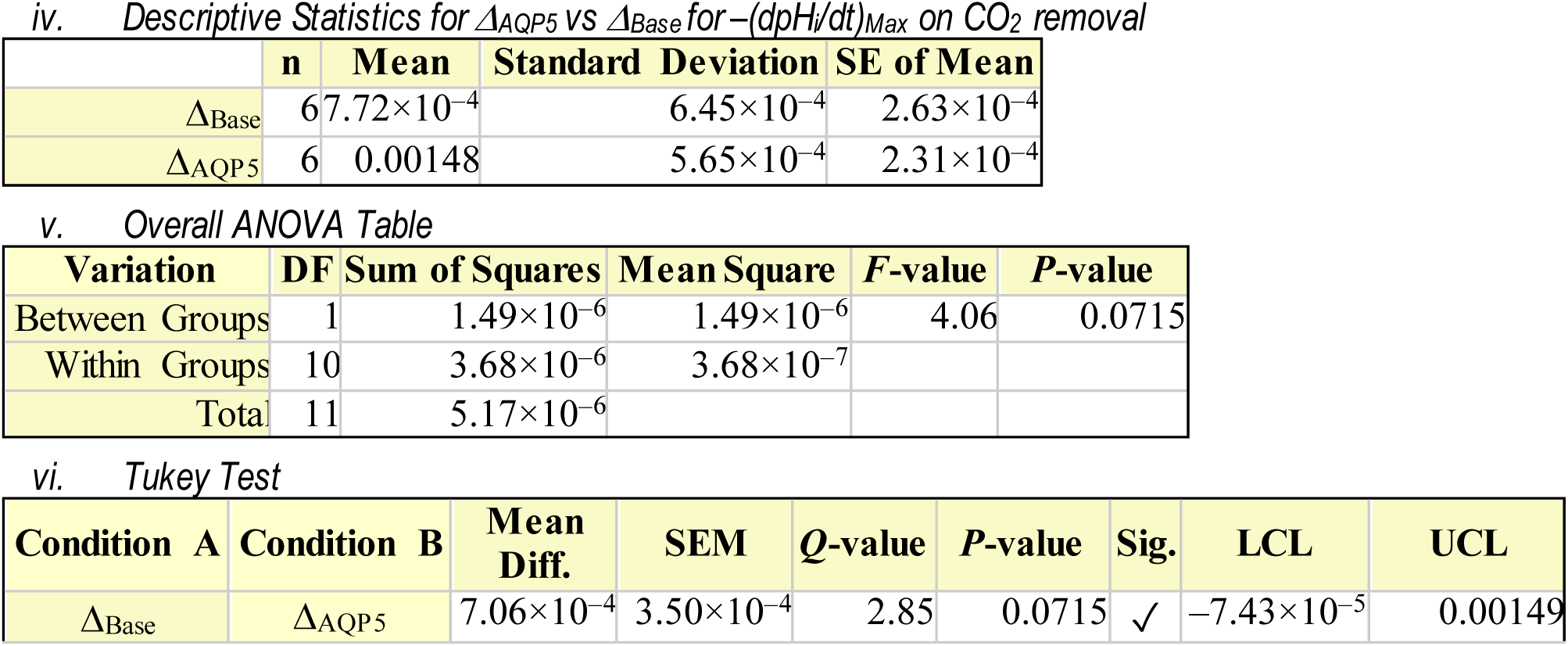
Statistics Summary for **Figure 12*B*, i**) Descriptive Statistics for data presented in Figure 12*B*, for comparison of select –(dpH_i_/dt)_Max_ bars extracted from Figure 3*D* and Figure 8*D* to analyze +(dpH_i_/dt)_Max_ upon CO_2_ removal in the absence or presence of bCA in bECF, on the “trans” side of the membrane to the pH_i_ measurement. ii) Overall one-way ANOVA results for +(dpH_i_/dt)_Max_ data presented in Figure 12*B*. DF is degrees of freedom. iii) Tukey’ s means comparison analysis for +(dpH_i_/dt)_Max_ data presented in Figure 12*B*. iv) Descriptive Statistics for data presented in Figure 12*B*, to analyze Δ_Base_ vs. Δ_AQP5_. v) One-way ANOVA results for Δ_Base_ vs. Δ_AQP5_ data presented in Figure 12*B*. vi) Tukey’s means comparison analysis for ΔΔ_Base_ vs. ΔΔ_AQP5_ data presented in Figure 12*B*. Threshold for significance α=0.05. Standard error of the mean for Tukey’s Test (SEM). Mean difference for each comparison (Mean Diff.), lower confidence limit (LCL), upper confidence limit (UCL).

## Footnotes

1 For simplicity, we refer to concentrations in the bECF using the subscript “o”.

2 Here, ±hAQP5 indicates that experiments in this section compare pH responses in the absence vs. presence of hAQP5.

3 Here, “Δ” indicates that, in this section, we compare a series of injected doses of hCA II; “–bCA” indicates that extracellular bCA was absent in all experiments.

4 In lists of points in the Summary and Conclusion, we use (a), (b), (c)…. The (a) in one list corresponds to the (a) in another list, etc. However, in the Interpretation, we list points as (1), (2), (3)…because the interpretations do not necessarily correspond to lettered points in a one-to-one fashion.

5 See “Results” > “±hAQP5, ΔhCA II, –bCA: Effects on ΔpH_S_ & (dpH_i_/dt)_Max_” > “ΔpH_S_” > “Interpretation: ΔpH_S_ (colorful bars) in Figure 3A,B; ±hAQP5, ΔhCA II, –bCA”.

6 See “Results” > “±hAQP5 in absence of exogenous CAs: Effects on ΔpH_S_ & (dpH_i_/dt)_Max_” > “(dpH_i_/dt)_Max_” > “Interpretation: (dpH_i_/dt)_Max_ (gray vs. black bars) in Figure 3*C,D*; ±hAQP5, –hCA II, –bCA”.

7 See Results > “±hAQP5in absence of exogenous CAs: Effects on ΔpH_S_ & (dpH_i_/dt)_Max_” _>_ “pH_S_” > “Interpretation: ΔpHS (gray vs. black bars) in Figure 3A,B; ±hAQP5, –hCA II, –bCA”. Also see Results > “±hAQP5, ΔhCA II,7 –bCA: Effects on ΔpH_S_ & (dpH_i_/dt)_Max_” > “pH_S_” > “Interpretation: ΔpH_S_ (colorful bars) in Figure 3A,B; ±hAQP5, –hCA II, –bCA”.

8 See Results > “±hAQP5, –hCA II, +bCA: Effects on ΔpH_S_ & (dpH_i_/dt)_Max_” > *“pH_S_” > “Interpretation: ΔpH_S_ (gray vs. black bars) in Figure 8A,B: ±hAQP5, –hCA II, +bCA”*.

9 See Results > “±hAQP5, –hCA II, +bCA: Effects on ΔpH_S_ & (dpH_i_/dt)_Max_” > *“pH_S_” > “Interpretation: ΔpH_S_ (gray vs. black bars) in Figure 8A,B: ±hAQP5, –hCA II, +bCA”*.

10 See Results > “±hAQP5, +hCA II, +bCA: Effects on ΔpH_S_ & (dpH_i_/dt)_Max_” > *“pH_S_” > “Interpretation: ΔpH_S_ (colorful bars) in Figure 8A,B: ±hAQP5, +hCA II, +bCA”*.

11 See Results > “±hAQP5, +hCA II, +bCA: Effects on ΔpH_S_ & (dpH_i_/dt)_Max_” > *“pH_S_” > “Interpretation: ΔpH_S_ (colorful bars) in Figure 8A,B: ±hAQP5, +hCA II, +bCA”*.

12 2.22×10^−308^ is the smallest possible value for double type data that a 64-bit systemis able to distinguish.

13 2.22×10^−308^ is the smallest possible value for double type data that a 64-bit systemis able to distinguish

14 2.22×10^−308^ is the smallest possible value for double type data that a 64-bit systemis able to distinguish

15 See Methods” > “Calculation of ΔpH_S_”, “Methods” > “Determination of “maximal” dpH_i_/dt”, “Methods” > “Calculation of intrinsic intracellular buffering power”

## References

Alishahi M & Kamali R (2019). A novel molecular dynamics study of CO_2_ permeation through aquaporin-5. Eur Phys J E 42, 151.

Blaydon DC, Lind LK, Plagnol V, Linton KJ, Smith FJD, Wilson NJ, McLean WHI, Munro CS, South AP, Leigh IM, O’Toole EA, Lundström A & Kelsell DP (2013). Mutations in AQP5, encoding a water-channel protein, cause autosomal-dominant diffuse nonepidermolytic palmoplantar keratoderma. Am J Hum Genet 93, 330–335.

Boone CD, Pinard M, McKenna R & Silverman D (2014). Catalytic Mechanism of α-Class Carbonic Anhydrases: CO_2_ Hydration and Proton Transfer. In Carbonic Anhydrase: Mechanism, Regulation, Links to Disease, and Industrial Applications, ed. Frost SC & McKenna R, Subcellular Biochemistry, pp. 31–52. Springer Netherlands, Dordrecht. Available at: 10.1007/978-94-007-7359-2_3.

Boron WF (1977). Intracellular pH transients in giant barnacle muscle fibers. Am J Physiol 233, C61–73.

Boron WF (2010). Sharpey-Schafer lecture: Gas channels. Exp Physiol 95, 1107–1130.

Boron WF & De Weer P (1976). Intracellular pH transients in squid giant axons caused by CO_2_, NH_3_, and metabolic inhibitors. J Gen Physiol 67, 91–112.

Calvanese L, Pellegrini-Calace M & Oliva R (2013). In silico study of human aquaporin AQP11 and AQP12 channels. Protein Sci 22, 455–466.

Calvetti D, Prezioso J, Occhipinti R, Boron WF & Somersalo E (2020). Computational model of electrode-induced microenvironmental effects on pH measurements near a cell membrane. SIAM Multiscale Model Simul 18, 1053–1075.

Chai RC, Jiang JH, Wong AYK, Jiang F, Gao K, Vatcher G & Yu ACH (2013). AQP5 is differentially regulated in astrocytes during metabolic and traumatic injuries. Glia 61, 1748–1765.

Chandy G, Zampighi GA, Kreman M & Hall JE (1997). Comparison of the water transporting properties of MIP and AQP1. J Membr Biol 159, 29–39.

Chen J, Lecuona E, Briva A, Welch LC & Sznajder JI (2008). Carbonic anhydrase II and alveolar fluid reabsorption during hypercapnia. Am J Respir Cell Mol Biol 38, 32–37.

Cooper GJ & Boron WF (1998). Effect of pCMBS on CO_2_ permeability of *Xenopus* oocytes expressing aquaporin 1 or its C189S mutant. Am J Physiol 275, C1481–1486.

Endeward V, Musa-Aziz R, Cooper GJ, Chen L-M, Pelletier MF, Virkki LV, Supuran CT, King LS, Boron WF & Gros G (2006). Evidence that aquaporin 1 is a major pathway for CO_2_ transport across the human erythrocyte membrane. FASEB J 20, 1974–1981.

Funaki H, Yamamoto T, Koyama Y, Kondo D, Yaoita E, Kawasaki K, Kobayashi H, Sawaguchi S, Abe H & Kihara I (1998). Localization and expression of AQP5 in cornea, serous salivary glands, and pulmonary epithelial cells. Am J Physiol 275, C1151–1157.

Geyer RR, Musa-Aziz R, Qin X & Boron WF (2013). Relative CO_2_/NH_3_ selectivities of mammalian aquaporins 0-9. Am J Physiol Cell Physiol 304, C985–994.

Gresz V, Kwon TH, Hurley PT, Varga G, Zelles T, Nielsen S, Case RM & Steward MC (2001). Identification and localization of aquaporin water channels in human salivary glands. Am J Physiol Gastrointest Liver Physiol 281, G247–254.

Gutknecht J, Bisson MA & Tosteson FC (1977). Diffusion of carbon dioxide through lipid bilayer membranes: effects of carbonic anhydrase, bicarbonate, and unstirred layers. J Gen Physiol 69, 779–794.

Heming TA, Geers C, Gros G, Bidani A & Crandall ED (1986). Effects of dextran-bound inhibitors on carbonic anhydrase activity in isolated rat lungs. J Appl Physiol Bethesda Md 1985 61, 1849–1856.

Heymann JB & Engel A (1999). Aquaporins: Phylogeny, structure, and physiology of water channels. News Physiol Sci Int J Physiol Prod Jointly Int Union Physiol Sci Am Physiol Soc 14, 187–193.

Ishibashi K (2009). New members of mammalian aquaporins: AQP10-AQP12. Handb Exp Pharmacol 251–262.

Ishibashi K, Kuwahara M, Gu Y, Kageyama Y, Tohsaka A, Suzuki F, Marumo F & Sasaki S (1997). Cloning and functional expression of a new water channel abundantly expressed in the testis permeable to water, glycerol, and urea. J Biol Chem 272, 20782–20786.

Ishibashi K, Morishita Y & Tanaka Y (2017). The evolutionary aspects of aquaporin family. In *Aquaporins*, ed. Yang B, Advances in Experimental Medicine and Biology, pp. 35–50. Springer Netherlands, Dordrecht. Available at: 10.1007/978-94-024-1057-0_2.

Ishida N, Hirai SI & Mita S (1997). Immunolocalization of aquaporin homologs in mouse lacrimal glands. Biochem Biophys Res Commun 238, 891–895.

King LS, Kozono D & Agre P (2004). From structure to disease: the evolving tale of aquaporin biology. Nat Rev Mol Cell Biol 5, 687–698.

Krane CM, Fortner CN, Hand AR, McGraw DW, Lorenz JN, Wert SE, Towne JE, Paul RJ, Whitsett JA & Menon AG (2001). Aquaporin 5-deficient mouse lungs are hyperresponsive to choline rgic stimulation. Proc Natl Acad Sci U S A 98, 14114–14119.

Kreda SM, Gynn MC, Fenstermacher DA, Boucher RC & Gabriel SE (2001). Expression and localization of epithelial aquaporins in the adult human lung. Am J Respir Cell Mol Biol 24, 224–234.

Kumari SS, Varadaraj M, Yerramilli VS, Menon AG & Varadaraj K (2012). Spatial expression of aquaporin 5 in mammalian cornea and lens, and regulation of its localization by phosphokinase A. Mol Vis 18, 957–967.

Larsen HS, Aure MH, Peters SB, Larsen M, Messelt EB & Kanli Galtung H (2011). Localization of AQP5 during development of the mouse submandibular salivary gland. J Mol Histol 42, 71–81.

Li C & Wang W (2017). Molecular Biology of Aquaporins. In *Aquaporins*, ed. Yang B, Advances in Experimental Medicine and Biology, pp. 1–34. Springer Netherlands, Dordrecht. Available at: 10.1007/978-94-024-1057-0_1.

Lien YH & Lai LW (1998). Respiratory acidosis in carbonic anhydrase II-deficient mice. Am J Physiol 274, L301–304.

Lu J, Daly CM, Parker MD, Gill HS, Piermarini PM, Pelletier MF & Boron WF (2006). Effect of human carbonic anhydrase II on the activity of the human electrogenic Na/HCO_3_ cotransporter NBCe1-A in *Xenopus* oocytes. J Biol Chem 281, 19241–19250.

Ma T, Song Y, Gillespie A, Carlson EJ, Epstein CJ & Verkman AS (1999). Defective secretion of saliva in transgenic mice lacking aquaporin-5 water channels. J Biol Chem 274, 20071–20074.

Matsuzaki T, Hata H, Ozawa H & Takata K (2009). Immunohistochemical localization of the aquaporins AQP1, AQP3, AQP4, and AQP5 in the mouse respiratory system. Acta Histochem Cytochem 42, 159–169.

Matsuzaki T, Tajika Y, Suzuki T, Aoki T, Hagiwara H & Takata K (2003). Immunolocalization of the water channel, aquaporin-5 (AQP5), in the rat digestive system. Arch Histol Cytol 66, 307–315.

Maurel C, Boursiac Y, Luu D-T, Santoni V, Shahzad Z & Verdoucq L (2015). Aquaporins in plants. Physiol Rev 95, 1321–1358.

Michalek K (2016). Aquaglyceroporins in the kidney: present state of knowledge and prospects. J Physiol Pharmacol Off J Pol Physiol Soc 67, 185–193.

Moss FJ & Boron WF (2020). Carbonic anhydrases enhance activity of endogenous Na-H exchangers and not the electrogenic Na/HCO_3_ cotransporter NBCe1-A, expressed in *Xenopus* oocytes. J Physiol 598, 5821–5856.

Moss FJ, Mahinthichaichan P, Lodowski DT, Kowatz T, Tajkhorshid E, Engel A, Boron WF & Vahedi-Faridi A (2020). Aquaporin-7: A dynamic aquaglyceroporin with greater water and glycerol permeability than its bacterial homolog GlpF. Front Physiol; DOI: 10.3389/fphys.2020.00728.

Moss FJ, Zhao P, Salameh AI, Taki S, Wass AB, Jacobberger JW, Huffman DE, Meyerson HJ, Occhipinti R & Boron WF (2025). Role of channels in the O_2_ permeability of murine red blood cells II. Morphological and proteomic studies. bioRxiv; DOI: 10.1101/2025.03.05.639962.

Musa-Aziz R, Boron WF & Parker MD (2010). Using fluorometry and ion-sensitive microelectrodes to study the functional expression of heterologously-expressed ion channels and transporters in *Xenopus* oocytes. Methods San Diego Calif 51, 134–145.

Musa-Aziz R, Chen L-M & Boron WF (2007). Differential effects of DIDS on the CO_2_ vs NH_3_ permeability of AQP1, AmtB, and RhAG. Proc Int Congr Nephrol Rio Jan Braz.

Musa-Aziz R, Chen L-M, Pelletier MF & Boron WF (2009). Relative CO_2_/NH_3_ selectivities of AQP1, AQP4, AQP5, AmtB, and RhAG. Proc Natl Acad Sci U S A 106, 5406–5411.

Musa-Aziz R, Geyer RR, Moss FJ & Boron WF (2025). Mechanism of CO_2_ and NH_3_ transport through human aquaporin 1: Evidence for parallel CO_2_ pathways. bioRxiv; DOI: 10.1101/2025.02.28.640247.

Musa-Aziz R, Occhipinti R & Boron WF (2014a). Evidence from simultaneous intracellular-and surface-pH transients that carbonic anhydrase II enhances CO_2_ fluxes across *Xenopus* oocytes plasma membranes. Am J Physiol Cell Physiol 307, C791–813.

Musa-Aziz R, Occhipinti R & Boron WF (2014b). Evidence from simultaneous intracellular-and surface-pH transients that carbonic anhydrase IV enhances CO_2_ fluxes across *Xenopus* oocyte plasma membranes. Am J Physiol Cell Physiol 307, C814–840.

Nakhoul NL, Davis BA, Romero MF & Boron WF (1998). Effect of expressing the water channel aquaporin-1 on the CO_2_ permeability of *Xenopus* oocytes. Am J Physiol 274, C543–548.

Nejsum LN, Kwon T-H, Jensen UB, Fumagalli O, Frøkiaer J, Krane CM, Menon AG, King LS, Agre PC & Nielsen S (2002). Functional requirement of aquaporin-5 in plasma membranes of sweat glands. Proc Natl Acad Sci U S A 99, 511–516.

Ni Z-X, Cui J-M, Zhang N-Z & Fu B-Q (2017). Structural and evolutionary divergence of aquaporins in parasites (Review). Mol Med Rep 15, 3943–3948.

Nielsen S, King LS, Christensen BM & Agre P (1997). Aquaporins in complex tissues. II. Subcellular distribution in respiratory and glandular tissues of rat. Am J Physiol-Cell Physiol 273, C1549– C1561.

Occhipinti R & Boron WF (2019). Role of carbonic anhydrases and inhibitors in acid-base physiolo gy: Insights from mathematical modeling. Int J Mol Sci 20, 3841.

Occhipinti R, Musa-Aziz R & Boron WF (2014). Evidence from mathematical modeling that carbonic anhydrase II and IV enhance CO_2_ fluxes across *Xenopus* oocytes plasma membranes. Am J Physiol Cell Physiol 307, C841–858.

Occhipinti R, Zhao P, Moss FJ & Boron WF (2025). Role of channels in the O_2_ permeability of murine red blood cells. III. Mathematical modeling and simulations. bioRxiv; DOI: 10.1101/2025.03.05.639964.

Parker MD, Musa-Aziz R, Rojas JD, Choi I, Daly CM & Boron WF (2008). Characterization of human SLC4A10 as an electroneutral Na^+^/HCO_3_^−^ cotransporter (NBCn2) with Cl^−^ self-exchange activit y. J Biol Chem 283, 12777–12788.

Parvin MN, Tsumura K, Akamatsu T, Kanamori N & Hosoi K (2002). Expression and localization of AQP5 in the stomach and duodenum of the rat. Biochim Biophys Acta 1542, 116–124.

Procino G, Mastrofrancesco L, Sallustio F, Costantino V, Barbieri C, Pisani F, Schena FP, Svelto M & Valenti G (2011). AQP5 Is expressed In type-B intercalated cells in the collecting duct system of the rat, mouse and human kidney. Cell Physiol Biochem 28, 683–692.

Purkerson JM & Schwartz GJ (2007). The role of carbonic anhydrases in renal physiology. Kidney Int 71, 103–115.

Qin X & Boron WF (2012). A single-amino acid mutation converts human aquaporin 5 to an anion channel. FASEB J 26, 1103.7.

Raina S, Preston GM, Guggino WB & Agre P (1995). Molecular cloning and characterization of an aquaporin cDNA from salivary, lacrimal, and respiratory tissues. J Biol Chem 270, 1908–1912.

Roos A & Boron WF (1981). Intracellular pH. Physiol Rev 61, 296–434.

Shinn E, Kowatz T, Vahedi-Faridi A, Boron WF & Tajkhorshid E (2024). Permeation of CO_2_ through the central pore of human AQP5. Biophys J 123, 265a.

Somersalo E, Occhipinti R, Boron WF & Calvetti D (2012). A reaction-diffusion model of CO_2_ influx into an oocyte. J Theor Biol 309, 185–203.

Steinfeld S, Cogan E, King LS, Agre P, Kiss R & Delporte C (2001). Abnormal distribution of aquaporin-5 water channel protein in salivary glands from Sjögren’s syndrome patients. Lab Investig J Tech Methods Pathol 81, 143–148.

Taki K, Kato H & Yoshida I (1999). Elimination of CO2 in patients with carbonic anhydrase II deficienc y, with studies of respiratory function at rest. Respir Med 93, 536–539.

Tsubota K, Hirai S, King LS, Agre P & Ishida N (2001). Defective cellular trafficking of lacrimal gland aquaporin-5 in Sjögren’s syndrome. Lancet Lond Engl 357, 688–689.

Vilas G, Krishnan D, Loganathan SK, Malhotra D, Liu L, Beggs MR, Gena P, Calamita G, Jung M, Zimmermann R, Tamma G, Casey JR & Alexander RT (2015). Increased water flux induced by an aquaporin-1/carbonic anhydrase II interaction. Mol Biol Cell 26, 1106–1118.

Wang Y, Cohen J, Boron WF, Schulten K & Tajkhorshid E (2007). Exploring gas permeability of cellular membranes and membrane channels with molecular dynamics. J Struct Biol 157, 534–544.

Wroblewski S (1879). On the nature of the absorption of gases. Nature 21, 190–192.

Yoshimura S, Nakamura H, Horai Y, Nakajima H, Shiraishi H, Hayashi T, Takahashi T & Kawakami A (2016). Abnormal distribution of AQP5 in labial salivary glands is associated with poor saliva secretion in patients with Sjögren’s syndrome including neuromyelitis optica complicated patients. Mod Rheumatol 26, 384–390.

Zhang L, Yao D, Zhou F, Zhang Q, Xia Y, Wang Q, Qin A, Zhao J, Li D, Zhou L & Cao Y (2019). The structural basis for glycerol permeation by human AQP7. bioRxiv 858332.

Zhang YB & Chen LY (2013). In silico study of Aquaporin V: Effects and affinity of the central pore-occluding lipid. Biophys Chem 171, 24–30.

Zhao P, Moss FJ, Occhipinti R, Geyer RR, Huffman DE, Gros G, Meyerson HJ & Boron W F (2025). Role of channels in the O_2_ permeability of murine red blood cells. I. Stopped-flow and hematological studies. bioRxiv; DOI: 10.1101/2025.03.05.639948.

